# Distinct sequence grammars of nucleosomal and linker DNA shape mutational and methylation landscapes in cancer

**DOI:** 10.64898/2026.09.28.755196

**Authors:** Yosef Masoudi-Sobhanzadeh, Sushant Kumar

## Abstract

Nucleosomes regulate DNA accessibility and thereby influence gene expression, DNA repair, and mutagenesis. Although DNA sequence strongly determines nucleosome positioning, the higher-order sequence grammar underlying nucleosome organization and its relationship to cancer-associated mutational and epigenetic processes remain poorly understood. We developed *DeepND*, an interpretable deep learning framework that distinguishes nucleosomal from inter-nucleosomal DNA and identifies sequence features associated with nucleosome architecture. Our interpretable framework enabled analysis of sequence patterns and identified 210 sequence motifs of 2–16 bp length that were significantly enriched or depleted in nucleosomal DNA, substantially extending the previously characterized nucleosomal motifs. These motifs revealed distinct sequence grammars within nucleosomal footprints and linker regions, including patterns associated with inward- and outward-facing minor grooves. Forty-four of these motifs showed cancer-type-specific enrichment or depletion of somatic mutations, and six motifs were associated with differential DNA methylation levels. In lung and endometrial cancers, mutations exhibited a significant ∼10-bp periodic enrichment pattern across distinct minor-groove orientations, consistent with differential sensitivity of mutational and DNA repair processes to nucleosomal rotational positioning. We further identified sequence-specific methylation patterns consistent with reduced accessibility of nucleosomal DNA to DNA methyltransferases, including differential methylation at inward- and outward-facing CpG motifs. Together, these findings provide a sequence-resolved characterization of nucleosome-level organization and its link to cancer-specific mutational processes and epigenetic landscapes.

## Introduction

Nucleosomes are the fundamental structural units of chromatin that enable long DNA sequences to be compacted and fit within the nucleus of the cell^1^. Each nucleosome consists of ∼147 base pairs of DNA (referred to as the nucleosomal footprint) wrapped around an octamer of histone proteins. Adjacent nucleosomes are linked by variable-length linkers, also known as inter-nucleosomal DNA^2^. The precise arrangement of nucleosomes along the genome, often referred to as nucleosome positioning, determines DNA accessibility for transcription factors, RNA polymerase, and DNA repair machinery. Therefore, nucleosome positioning plays a critical role in regulating various biological processes, including transcription, replication initiation, and tissue-specific epigenetics^3^. As a result, disruption of nucleosome positioning is often associated with various human diseases, including neurodevelopmental disorders and multiple cancer types.

Various intrinsic DNA sequence features strongly influence nucleosome positioning across the genome. For instance, WW (A/T-rich dinucleotides) and SS (C/G-rich dinucleotides) motifs are preferentially enriched at minor-groove sites facing inward or outward relative to histone proteins^4,5^. Motivated by the dependence of nucleosome positioning on DNA sequence features, multiple computational approaches, including deep learning methods, have been developed to predict nucleosome profiles from DNA sequences. However, these approaches often lack interpretability^6^ or primarily identify short sequence motifs, typically no longer than 4 nucleotides^7^. Consequently, the sequence grammar underlying nucleosome positioning remains incompletely characterized.

Prior studies have also demonstrated a close association between nucleosome positioning and mutational processes operating in the cancer genome^8,9^. For example, it has been shown that reduced accessibility of the nucleotide excision repair machinery within nucleosome-occupied DNA in cancer genomes often results in higher mutation rates in these regions^10–12^. Moreover, the biophysical properties of nucleosomes influence DNA damage and repair outcomes, creating a distinctive mutational periodicity along nucleosomal DNA that aligns with the minor-groove orientation of the underlying DNA segment^13^. Additionally, the translational and rotational positioning of DNA around the histone octamer can influence DNA damage and repair in a position-dependent manner^8,13^. Finally, nucleosome positioning is closely associated with DNA methylation, shaping the landscape of epigenetic dysregulation in cancer^14–16^. Specifically, structural modeling studies have shown that DNMT–nucleosome interactions contribute to methylation periodicity and nucleosome stability^17^. Collectively, these findings underscore nucleosomes as key factors influencing both mutation and methylation patterns in cancer.

Identifying nucleosomal DNA-related sequence motifs offers several advantages, including precise determination of the dyad position, which approximately corresponds to the midpoint of the 147-base-pair footprint DNA^18,19^. In particular, characterizing motifs associated with somatic mutations and DNA methylation may reveal sequence features underlying cancer-type-specific mutational and epigenetic patterns. Such motifs may also help clarify how local nucleosome structure and positioning influence genome stability and epigenetic regulation^11^. Furthermore, some motifs may exhibit cancer-type-specific enrichment, suggesting their potential role as molecular hallmarks of tumor origin. In addition to serving as mutation biomarkers, nucleosome-associated motifs are detectable in cfDNA, which has been linked to several diseases^20–23^. However, a comprehensive characterization of sequence motifs associated with both nucleosomal and inter-nucleosomal regions, along with their relationships to cancer-associated mutations and DNA methylation, remains lacking.

Overall, our work addresses the limited interpretability of existing nucleosome-positioning models and the restricted sequence length of motifs characterized by previous approaches. To this end, we developed *DeepND*, an interpretable deep residual neural network framework designed to interrogate sequence motifs spanning up to 30 nucleotides. We identified 210 informative sequence motifs ranging from 2 to 16 nucleotides that showed statistically significant enrichment or depletion in nucleosomal DNA, extending beyond the 4-nucleotide sequence patterns characterized by previous approaches. We further investigated associations between these motifs and nucleosome structure, particularly minor-groove orientations at histone–DNA contacts, as well as cancer-associated genomic alterations. We then examined somatic mutations and DNA methylation within the informative sequence motifs and investigated the correspondence between mutation-associated motifs and known mutational signatures. Together, these analyses provide, to the best of our knowledge, the first sequence-resolved characterization of nucleosome-associated sequence grammar and reveal its associations with nucleosome structure, cancer-associated mutations, and epigenetic alterations.

## Results

### An overview of the method

We used high-resolution MNase-seq data to develop a multi-stage machine learning framework based on a deep residual architecture to classify genomic regions as nucleosomal or inter-nucleosomal from DNA sequence and to identify sequence features associated with nucleosomal DNA. The framework utilizes MNase-seq-derived nucleosome-protected fragments, which are first aligned to the reference genome (**Figure 1A**). Next, we smooth the fragment-density profiles to accurately detect well-positioned nucleosomes and precisely determine the dyad position within each nucleosomal region. We then use the smoothed nucleosomal profile to label genomic regions as nucleosomal or inter-nucleosomal DNA. We generate one-hot encoded representations of the underlying DNA sequences to train a residual convolutional neural network. Our model architecture includes convolutional and pooling layers connected by skip connections, enabling the network to capture both local and long-range sequence dependencies relevant to nucleosome formation (**Figure 1B**). We applied an iterative training strategy to prevent overfitting by varying hyperparameters and using different optimization algorithms. After the optimal model converged, we combined the most predictive convolutional filters from multiple independent training runs to identify recurrent sequence patterns associated with nucleosome occupancy. From the trained models, we systematically extracted both short and higher-order sequence motifs that consistently contributed to predictive performance (**Figure 1C**). Overall, this framework provides a comprehensive and unbiased approach to identifying nucleosome-related sequence motifs. After generating a catalog of well-positioned nucleosomal sequence motifs, we examined their associations with mutational and methylation features in cancer genomes.

**Figure 1.**
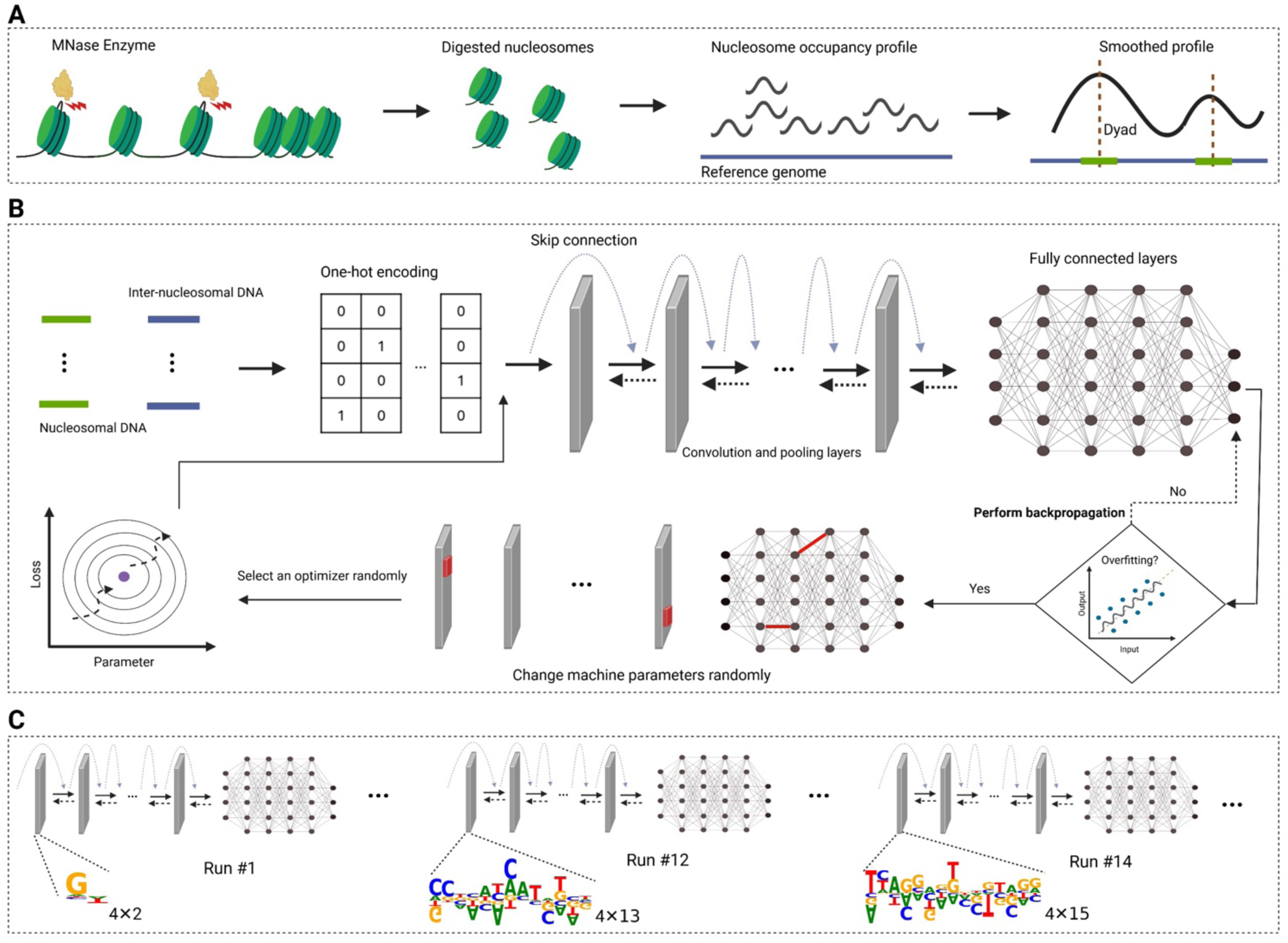
Overview of the *DeepND* framework. (**A**) MNase binds nucleosomal DNA and digests linker regions, generating DNA fragments of various lengths. The ∼147 bp mononucleosomal fragments are mapped to the human reference genome, and dyad positions are identified after smoothing the nucleosome occupancy profile. (**B**) Nucleosomal and inter-nucleosomal DNA sequences are represented using one-hot encoding, and a classifier is trained with deep residual networks. Trainable parameters highlighted in red in the kernels and fully connected layers are randomly selected and modified when overfitting occurs. (**C**) Nucleosomal DNA–associated sequence motifs are identified by running *DeepND* with different convolutional kernel sizes.

### Assessing the influence of short flanking DNA on classifier performance

Previous studies have shown that flanking linker DNA can influence nucleosome positioning, likely through its interaction with histone protein H1^24,25^. Therefore, we investigated whether incorporating short flanking linker sequences adjacent to nucleosomal DNA improves the performance of our machine learning model. To this end, we evaluated two distinct modeling strategies (**Figure 2A**). In the first scheme (Scheme 1), the input consisted solely of the canonical 147-bp nucleosomal footprint DNA. In the second scheme (Scheme 2), we extended the input to include both the 147-bp nucleosomal footprint and the flanking linker DNA (27 bp on each side of the dyad position), resulting in input DNA sequences of 201 bp in length. For the corresponding negative dataset (inter-nucleosomal DNA group), sequences of length 147 bp and 201 bp were selected for Schemes 1 and 2, respectively. Furthermore, we excluded nucleotides located 71–77 bp from the dyad position from all input sequences to minimize potential biases introduced by MNase digestion. These positions exhibit elevated MNase cleavage frequencies (**Figure 2B**); therefore, their inclusion will likely confound model training and evaluation.

**Figure 2.**
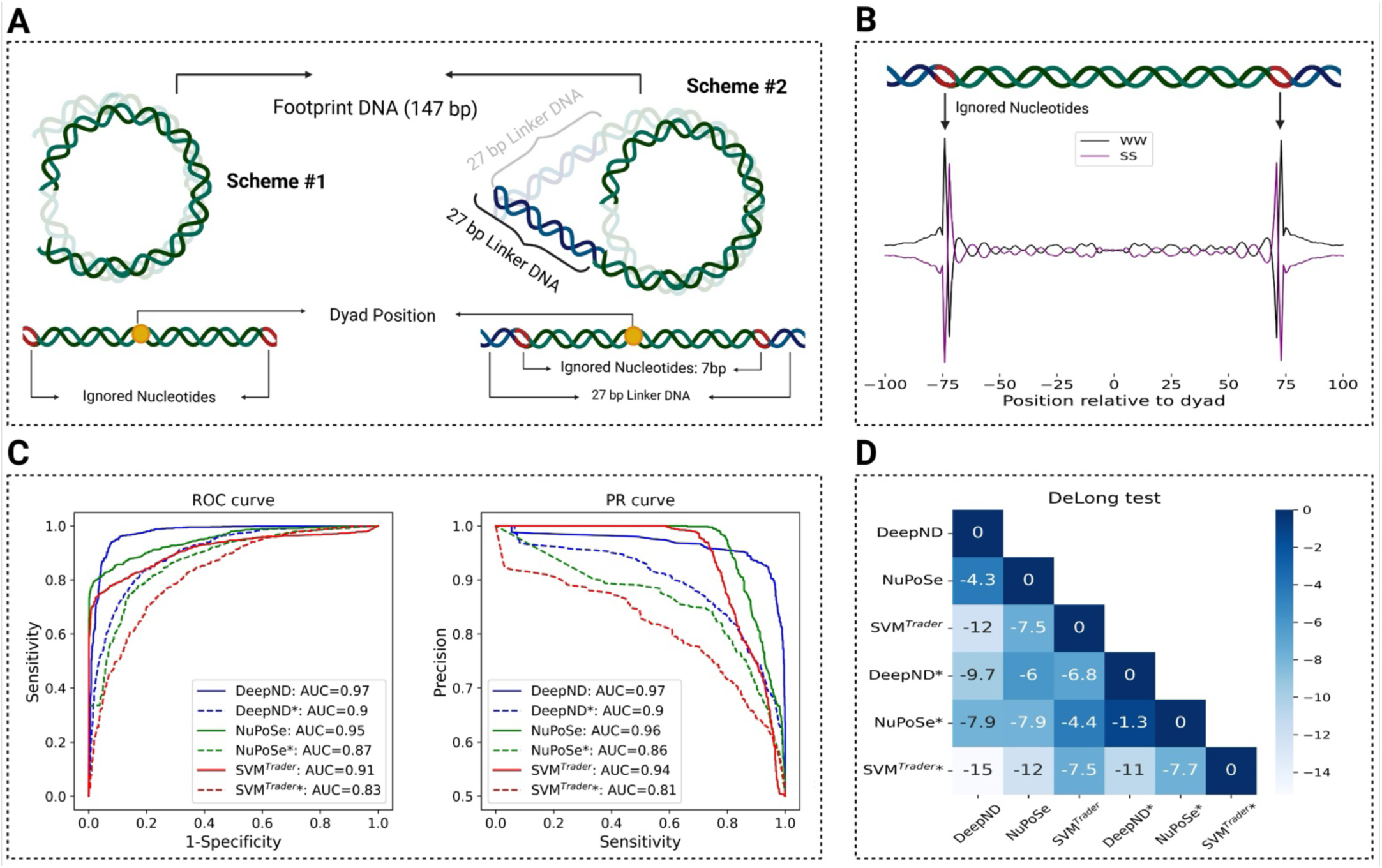
Assessing the influence of short flanking linker DNA on the performance of the classification models. (**A**) An illustration of two training schemes. Scheme #1 includes only the nucleosomal footprint DNA, whereas Scheme #2 includes both the footprint and the 27-bp flanking linker DNA. Green, red, and blue segments represent the nucleosomal footprint DNA, the 27-bp linker DNA, and ignored nucleotides, respectively. (**B**) Two prominent peaks are observed on either side of the dyad position because the MNase enzyme cleaves these sites much more frequently than other sites. (**C**) Model performance is evaluated using PR and ROC curves. Solid and dashed lines represent Schemes 1 and 2, respectively. The symbol * indicates classifiers that were not trained using short flanking linker DNA information. (**D**) The DeLong test was used to compare model performance. Negative values in the DeLong test matrix indicate that the method on the y-axis performs worse than the method on the x-axis.

A systematic comparison of our *DeepND* model with three widely used nucleosomal-versus-inter-nucleosomal DNA classifiers showed that incorporating flanking linker DNA substantially improved performance across multiple evaluation metrics (**Figure 2C**). For instance, the *DeepND* model that included linker DNA (Scheme 2) markedly outperformed the model based on Scheme 1 (MCC: 0.86 versus 0.63; AUC: 0.97 versus 0.9). Additionally, to statistically compare the AUCs of the methods, we applied the DeLong test across nucleosome classifiers and observed significant differences between the ROC curves of the various algorithms (**Figure 2D**), with z-scores greater than 2 corresponding to *p* < 0.05. Moreover, comparison of different nucleosomal and inter-nucleosomal DNA classifiers (**Table S1**) showed that the *DeepND* model based on Scheme 2 achieved the best overall performance across multiple metrics, including MCC (0.86), F1 score (0.93), and Cohen’s kappa (0.88). Together, these results highlight the strong and consistent benefit of incorporating flanking linker DNA for model performance, independent of the underlying sequence representation strategies (one-hot encoding or handcrafted sequence features) or the machine-learning algorithm employed.

Previous machine-learning methods for classifying nucleosomal and inter-nucleosomal DNA have typically been benchmarked using three gold-standard datasets derived from human (CD4+ T cells), *C. elegans*, and *D. melanogaster*^26–28^. Due to computational constraints, prior methods rely on relatively small subsets of genomes for training, with each dataset comprising approximately 5,000 DNA fragments spanning both nucleosomal and inter-nucleosomal DNA. In contrast, the MNase-Seq data utilized in our study incorporates the highest-resolution nucleosomal DNA currently available. We also evaluated and compared the performance of *DeepND* with other methods on these three benchmarking datasets across a range of flanking linker DNA lengths (**Figure S1** and **Table S2**)^24,29–32^.

Across species, the length of the flanking linker DNA was observed to affect *DeepND’s* performance. For *H. sapiens*, *C. elegans*, and *D. melanogaster*, optimal performance was achieved with linker DNA lengths of approximately 10–20 bp, 90–130 bp, and 60–90 bp, respectively (Figure S1A). These differences in optimal linker DNA length are consistent with species-specific differences in chromatin organization and nucleosome spacing. Mechanistically, chromatin compaction is modulated by linker DNA length and nucleosome interactions, with shorter spacing promoting more compact chromatin structures^33^. Additionally, we observed a decline in model performance with longer flanking linker DNA (an average decrease of 7%, as shown in Figure S1A), likely because the extended flanking regions begin to encompass both nucleosomal and inter-nucleosomal DNA, thereby affecting discriminative features. Overall, *DeepND* outperformed existing nucleosomal DNA classification methods across all three benchmarking datasets, achieving an average MCC of 0.83 based on 5-fold cross-validation (**Table S2**).

### Identification of nucleosomal DNA-related sequence motifs

We focused on identifying informative sequence motifs associated with nucleosomal DNA regions by systematically analyzing the first-layer convolutional kernels of DeepND and quantifying their associations using Fisher’s exact test (as described in the Materials and Methods section). Our analysis identified 136 sequence motifs derived from nucleosomal footprint DNA (**Figures S2–S5** and **Table S3**) and 64 sequence motifs derived from flanking linker DNA regions (**Figure S5** and **Table S4**). Notably, we identified, for the first time, long sequence motifs ranging from 2 to 16 bp that are significantly enriched or depleted in nucleosomal DNA, whereas previous studies have largely been limited to motifs of length 2–4 bp. Most informative sequence motifs ranged from 4–6 bp in footprint DNA and 2–9 bp in flanking linker DNA regions (Figure S6). Among these motifs, AGTACAGGCACAG (OR: 1.89) and AGTCGGCGCCAGGACG (OR: 0.23) exhibited the highest enrichment and depletion in nucleosomal DNA, respectively. Here, odds ratios greater than 1 indicate enrichment of a motif in nucleosomal DNA, while those less than 1 indicate depletion.

We next evaluated the relative importance of the identified sequence motifs and prioritized those that contributed most to our classifier’s performance (**Figure 3** and **Figure S7**). Because multiple motifs exhibited strong pairwise Pearson correlations (r > 0.5), we grouped them accordingly (**Figure 3A** and **Figure 7A**). We observed that the distribution of informative sequence motifs differed significantly between nucleosomal and inter-nucleosomal DNA groups (**Figure 3B** and **Figure S7B**). We ranked these motifs by their contribution to model accuracy (see Methods section) by measuring the reduction in classification accuracy after removing each motif individually (**Figure 3C** and **Figure 3D**). Among these groups, Group 3 (G3) motifs had the greatest impact on classifier performance, as their exclusion from our model resulted in the largest decline in AUC (from ∼0.91 to ∼0.76). This group comprised eight sequence motifs spanning both nucleosomal footprint and flanking linker DNA regions. Footprint DNA-derived motifs included several A/T-rich patterns (e.g., AAAAA/TTTTT and AA/TT), whereas linker DNA-associated motifs showed simpler dinucleotide and trinucleotide patterns such as TA and TAA/TTA.

**Figure 3.**
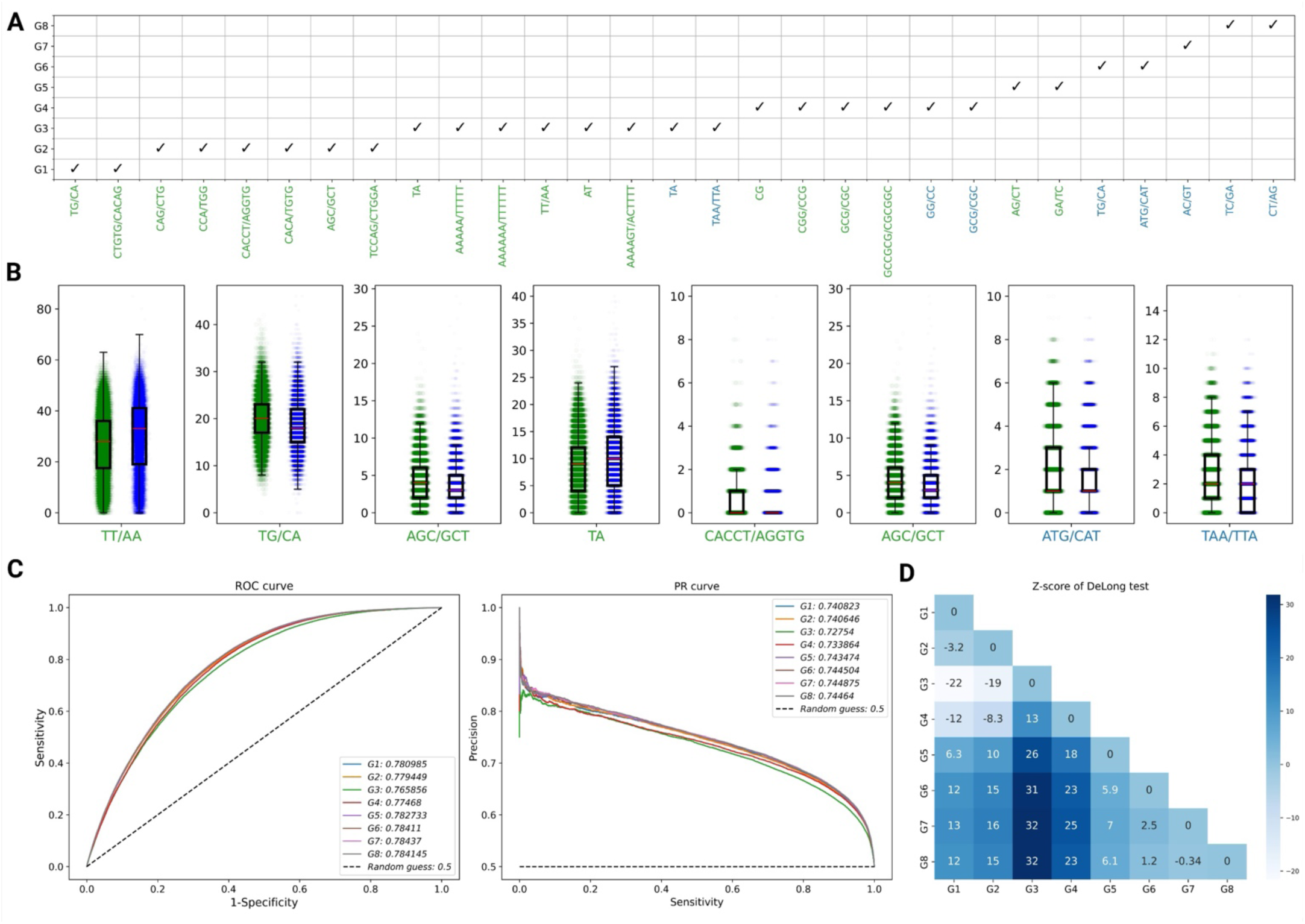
Filtering informative sequence motifs and ranking them by their impact on the classifier. (**A**) Selected informative sequence motifs grouped by Pearson correlation coefficients (>0.5). (**B**) Frequencies of the selected informative sequence motifs in nucleosomal and inter-nucleosomal DNA regions. A complete list of all features is provided in the supplement figure S7B. (**C**) Effect of removing each motif group on classifier performance. A larger decrease in AUC upon removal indicates greater importance of the corresponding group. (**D**) The DeLong test compares classifier performance after removing different motif groups; positive values indicate that the group on the x-axis has a greater impact than the group on the y-axis.

Although we trained the *DeepND* framework using convolutional kernels with sizes ranging from 2 to 30 bp, kernels larger than 16 bp were not directly interpretable as discrete sequence motifs. However, these long kernels exhibited a distinctive structural pattern, frequently consisting of repeated or identical subregions arranged at approximately 10 bp intervals. Nucleosomal DNA is known to exhibit a ∼10–11 bp periodicity at minor-groove sites (**Figure 4A)**. This periodicity arises from the alternating orientation of minor-groove sites facing inward and outward relative to the histone octamer^34^. We hypothesized that these repeated subregions correspond to minor-groove-associated sequence features within nucleosomal DNA. Accordingly, we identified these repeated subregions as periodicity-related features and quantified their periodic behavior using Fast Fourier Transform (FFT) analysis. Using this approach, we identified 80 periodicity-related sequence motifs with lengths ranging from 4 to 6 bp that were significantly enriched at minor-groove sites, as determined by Fisher’s exact test (**Figures 4B** and **S8-S13**; and **Table S5**). Previous studies have primarily reported nucleosomal periodicity for dinucleotide motifs and, more rarely, for 3–4 bp motifs. For instance, we found the GGAGGT/ACCTCC sequence motif with the longest periodicity, which is associated with minor-groove sites facing the histone octamer and exhibits a strong periodicity pattern across nucleosomal DNA (**Figures 4C** and **4D**).

**Figure 4.**
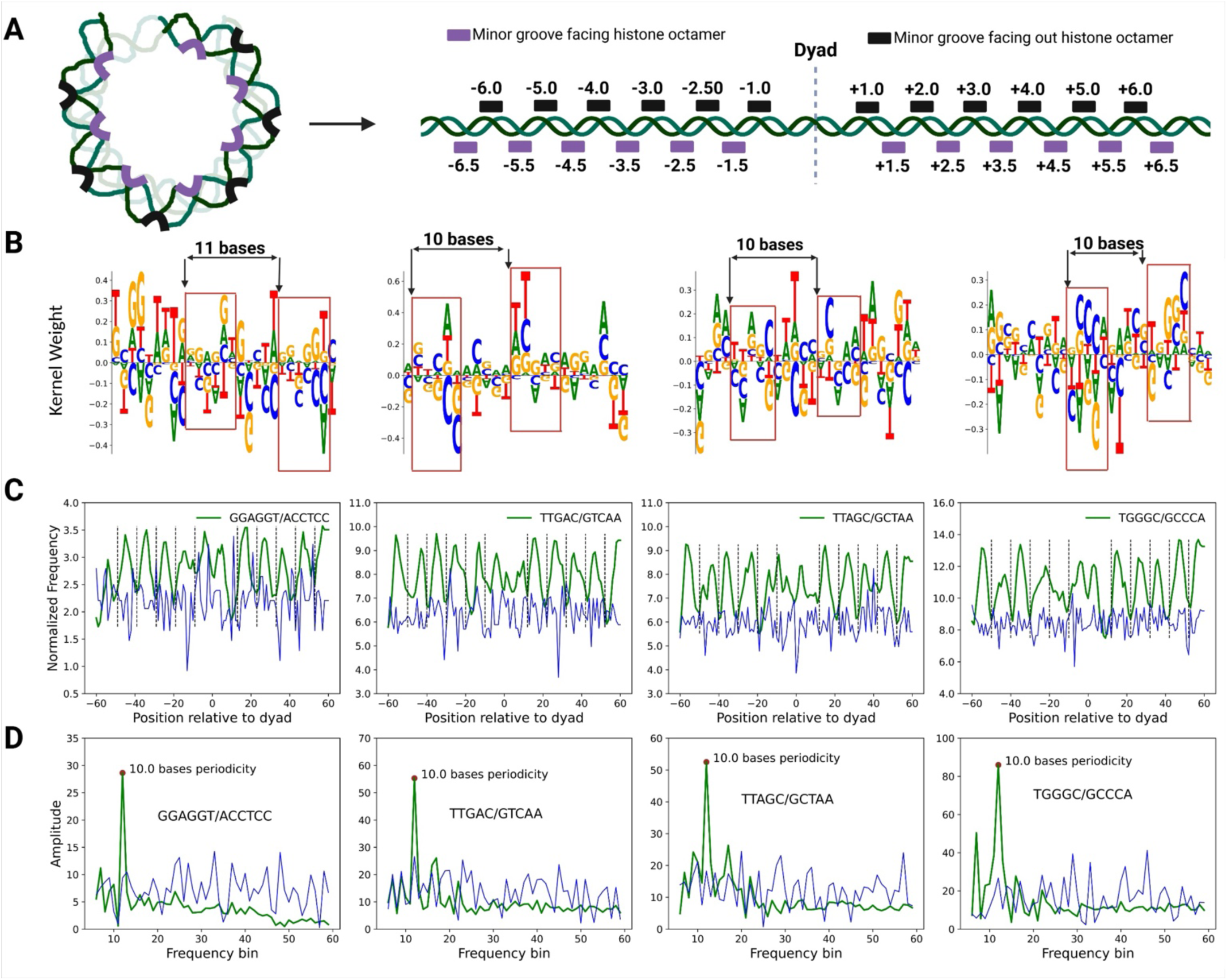
Periodicity-based sequence motifs learned by DeepND. (**A**) Structure of nucleosomal DNA, with minor-groove sites facing inward and outward relative to the histone octamer. Purple and black rectangles correspond to minor-groove sites facing outward from and inward toward the histone octamer, respectively. (**B**) Sequence logos of four kernels containing identical subregions spaced 10–11 bp apart. These identical subregions correspond to periodicity-related sequence motifs. All identified periodicity-related sequence motifs are provided in Figures S8–S13 and Table S5. (**C**) Frequencies of periodicity-related sequence motifs in nucleosomal (green) and inter-nucleosomal (blue) DNA sites. The dashed lines show minor-groove sites facing outward from the histone octamer. (**D**) Fast Fourier transform analysis of motif frequencies, showing a strong ∼10 bp periodic pattern in the nucleosomal DNA dataset.

### Detecting mutational motifs and their associations with different cancer types

We investigated whether nucleosome positioning is significantly associated with somatic mutation burden and underlying mutational processes in cancer genomes. Specifically, we examined whether somatic mutational patterns within nucleosomal DNA differ from those in inter-nucleosomal regions, whether somatic mutations preferentially occupy specific nucleosome-associated sequence motifs (hereafter referred to as mutational motifs), and whether these motif-level mutational patterns vary across cancer types and relate to known mutational signatures. Since nucleosome positioning is influenced by various biological and epigenetic factors, our primary analyses focused on mutations affecting well-positioned nucleosomes, whose positions are largely consistent across cell/tissue types^35^. Furthermore, we primarily relied on Mutect2-based mutation calls generated for the CPTAC study^36,37^ for this analysis. We also repeated this analysis for non-well-positioned nucleosomes and mutations identified by tools other than MuTect2 (**Figures S14–S20** and **Tables S6–S21)**.

A close inspection of single-nucleotide variant (SNV) pattern comparisons at the pan-cancer level between nucleosomal and inter-nucleosomal DNA revealed a significant association between specific SNV types and nucleosomal footprint DNA of well-positioned nucleosomes (**Figure S14**). In particular, T>C mutations were significantly depleted (OR: ∼0.8, adjusted p < 0.01), whereas C>A mutations were significantly enriched (OR: ∼1.1, adjusted p < 0.05) within nucleosomal footprint DNA (**Figure 5A and Tables S6-S8**). Although we observed small differences in mean fractions for other SNV types (C>G, T>A, etc.), these differences were not statistically significant after multiple-hypothesis corrections. Interestingly, analysis of somatic mutations in both well-positioned and non-well-positioned nucleosomes revealed additional significant associations between SNV patterns and nucleosomal DNA (**Table S6**). We also observed that mutational enrichment and depletion patterns varied across chromosomal contexts and genomic compositions. For example, T>A mutations were significantly enriched on chromosome 9 and depleted on chromosome 6 (**Tables S6–S8**). Compared with nucleosomal footprint DNA (**Figures S14–S15 and Tables S6–S11**), some SNV patterns were enriched in flanking linker DNA regions but depleted in nucleosomal DNA regions (e.g., T>G mutations in nucleosomal and inter-nucleosomal regions), underscoring the context-dependent influence of nucleosome organization on mutational processes.

**Figure 5.**
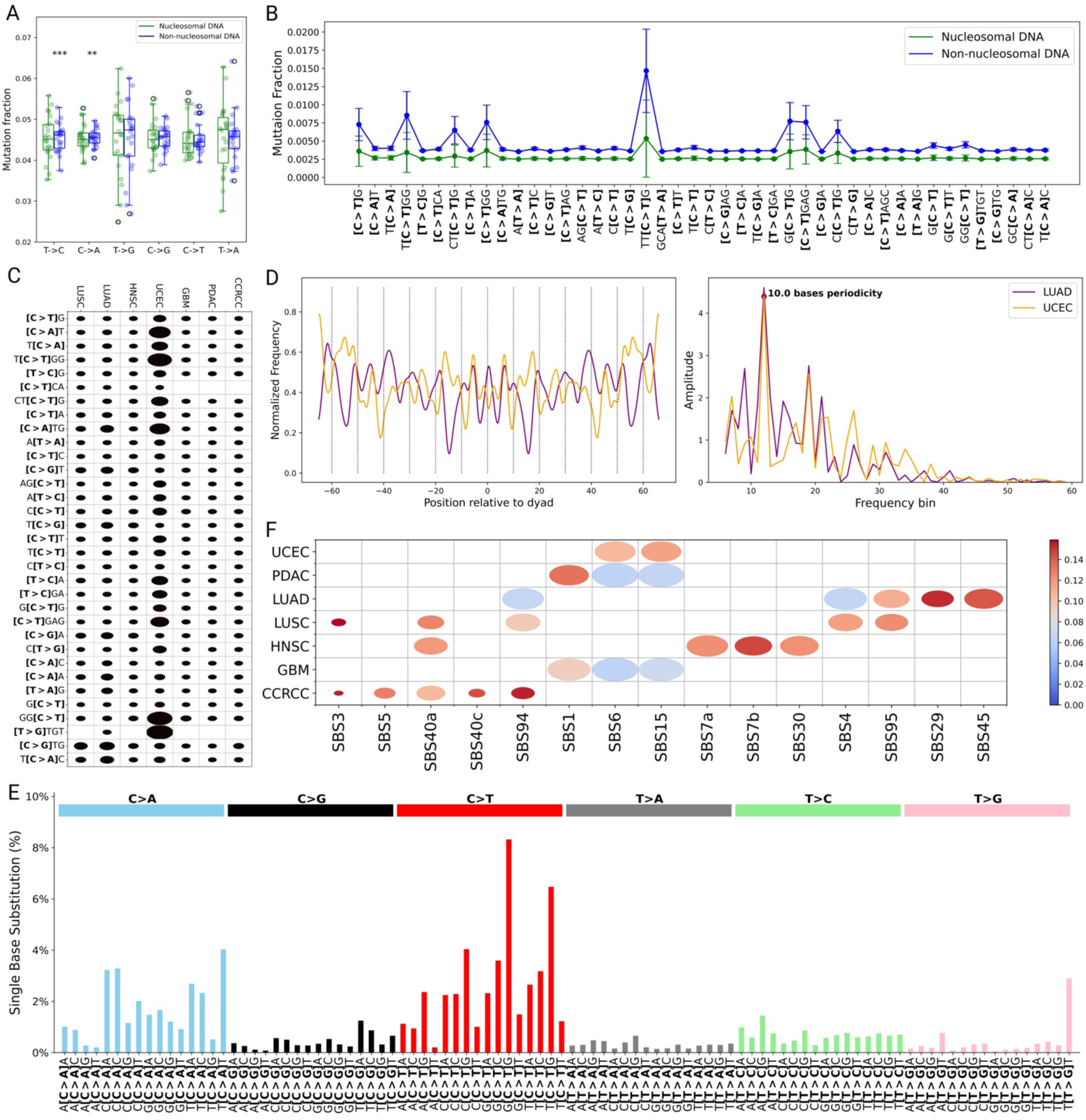
Associations between well-positioned nucleosomes and cancer mutational processes. (**A**) Distribution of single-nucleotide mutations in nucleosomal (green) and inter-nucleosomal regions. Symbols *** and ** indicate significant associations with p < 0.01 and p < 0.05, respectively. The mutation fraction is the number of a specific single-nucleotide mutation divided by the total number of mutations. Each point in the boxplots represents an individual chromosome. (**B**) Mutational motifs identified as significantly enriched or depleted in well-positioned nucleosomes. Error bars represent variability in mutation rates across chromosomes. For clarity, this panel displays mutation fractions for chromosomes with the largest differences. Error bars indicate variability in mutation fractions across all chromosomes. (**C**) Mutational motifs significantly associated with different cancer types. Black indicates depletion of motifs in the respective cancer types. Circle size represents the normalized odds ratio from Fisher’s exact test. (**D**) Frequency of mutations in UCEC and LUAD. Dashed lines indicate minor groove sites facing outward from histone proteins. (**E**) Mutational profiles were generated from the identified motifs, following the order of the 96 trinucleotide substitution patterns in the COSMIC platform. (**F**) Similarity between the generated mutational profiles for each cancer type and the COSMIC mutational signatures. Circle size and color indicate the Pearson correlation coefficient and cosine similarity, respectively. This analysis included various cancer cohorts such as clear cell renal cell carcinoma (CCRCC), glioblastoma (GBM), head and neck squamous cell carcinoma (HNSC), lung squamous cell carcinoma (LUSC), lung adenocarcinoma (LUAD), pancreatic ductal adenocarcinoma (PDAC), and uterine corpus endometrial carcinoma (UCEC).

Next, we investigated whether somatic mutations are preferentially enriched or depleted within informative sequence motifs. Across nucleosomal regions (well-positioned and non-well-positioned), we observed that significant mutational enrichment was more prevalent than depletion within mutational motifs occupying nucleosomal footprint DNA (**Figures 5B and S16–S17; Tables S12–S17**). However, when restricting our analyses to well-positioned nucleosomes, we observed 44 mutational motifs that were significantly depleted within nucleosomal footprint regions (**Figure 5B**). These distinct observations suggest that nucleosomal positional stability influences motif-specific mutational patterns in the cancer genome. This nucleosomal context dependence in mutational patterns is particularly evident in [C > T]G mutations, which were significantly enriched across all nucleosomal regions (**Tables S12 and S13**) but were significantly depleted when restricted to well-positioned nucleosome regions (**Table S14**). In flanking linker DNA regions, mutational motifs were largely enriched in nucleosomal DNA compared to inter-nucleosomal linker regions (**Tables S15–S17),** further highlighting the influence of these flanking regions on mutational accumulation in cancer genomes.

Furthermore, we examined the relationship between the identified mutational motifs and various cancer types. We observed statistically significant associations, with certain mutational motifs depleted relative to their overall mutation frequencies across multiple cancer types (**Figure S18 and Tables S18–S20**; p < 0.01). This consistent depletion pattern may suggest a relationship between the identified mutational motifs and underlying mutational processes. Across all nucleosomal region categories (well- and non-well-positioned) and cancer types, [C > T]G mutations emerged as the most prominent mutational motif, indicating a potentially central role in cancer-associated mutagenesis (**Figure 5C and Figure S18**). Notably, when we restricted this analysis to well-positioned nucleosomes, the number of significant associations between mutational motifs and cancer types increased substantially (**Figure 5C and Figure S17).** Specifically, 33 of 44 mutational motifs exhibited significant cancer-type-specific associations. In contrast, we didn’t find any significant associations between mutational motifs in flanking linker DNA regions and cancer types, indicating that these relationships are largely confined to nucleosome-associated DNA.

To further examine the structural basis of these effects, we assessed whether mutations in specific nucleosomal regions are associated with cancer types. We observed modest but significant cancer-type–specific enrichment across minor-groove orientations (**Table S21 and Figure 5D**). Notably, mutations in uterine corpus endometrial carcinoma (UCEC) showed no enrichment at minor-groove sites facing away from histone proteins (OR = 0.97; Fisher’s exact test, *p* = 6.7 × 10⁻³⁷), whereas mutations in several other cancer types exhibited significant enrichment at outward-facing minor-groove sites (**Table S21**). Consistent with these findings, fast Fourier transform (FFT) analysis revealed a significant ∼10-bp periodicity of mutations at minor-groove sites in UCEC and lung adenocarcinoma (LUAD) (OR = 1.05, p ∼ 0.0), reflecting canonical rotational positioning of DNA around the nucleosome (**Figure 5D**). In contrast, most other cancer types exhibited weaker signals with an approximately 5-bp periodicity and lower FFT amplitudes (**Figure S19**). Together, these results suggest that nucleosome structural organization constrains the sequence contexts and spatial distribution of mutations across cancer types, whereas no such associations are evident in inter-nucleosomal or linker DNA regions.

Finally, we characterized mutational spectra of cancer-type–specific profiles derived from the identified mutational motifs. We restricted these analyses to nucleosome-associated sequence space, where we constructed mutational profiles for each cancer type by including only somatic mutations overlapping the genomic coordinates of at least one mutational motif identified for that cancer type. Therefore, these profiles describe the mutational spectrum within predicted nucleosomal motifs rather than the genome-wide spectrum of each cohort. Across cancer types, the motif-restricted profiles were primarily dominated by trinucleotide contexts involving C > T and C > A substitutions, with G[C > T]G the most frequent motif (**Figures 5E** and **S20**). Comparison against reference COSMIC signatures showed that several motif-restricted profiles resembled signatures previously reported in the corresponding cohorts (Figure 5F). For example, the motif-restricted mutational profiles for clear cell renal cell carcinoma (CCRCC) and lung squamous cell carcinoma (LUSC) were highly similar to established mutational signatures within these cohorts, respectively (Pearson correlation coefficients and cosine similarity values exceeding 0.8) (**Figure 5F**). Recovery of SBS4, the tobacco-smoking signature characteristic of LUSC, and SBS94 in CCRCC within nucleosome-associated contexts aligns with expected cohort-level mutational signature patterns. Similarly, the UCEC profile corresponded to the mismatch-repair-deficiency signatures SBS6 and SBS15, in addition to the clock-like signature SBS1, consistent with the frequent mismatch-repair deficiency reported in this cohort. Notably, the LUAD profile showed the strongest overall similarity to reference signatures, though this similarity was distributed across several C>A–dominated signatures (SBS4, SBS29, SBS45, SBS95) that could not be reliably distinguished from one another within motif-restricted sequence space. Overall, the similarity of these motif-restricted mutational profiles to existing COSMIC signatures indicates consistency with cohort-wide mutational processes rather than evidence that these signatures act preferentially within nucleosomal DNA.

### Association of methylation motifs with different cancer types

Beyond understanding the relationship between nucleosome positioning and the mutational profile, we also investigated whether nucleosome positioning is associated with CpG methylation patterns in tumor genomes. Toward this goal, we employed the same hypothesis-testing framework used for the mutational motif analyses and evaluated associations between nucleosome organization and the identified CpG methylation motifs. Our analysis identified five CpG methylation motifs within nucleosomal footprint DNA and one within nucleosomal flanking linker DNA regions that were statistically significant (**Figure 6A**). Specifically, we observed significant depletion of the CG, CGG, and CGAA motifs within nucleosomal footprint DNA, whereas the GCG and CGTT motifs were significantly enriched (*p* < 0.01). Notably, the GCG motif was enriched in both the nucleosomal footprint and the flanking linker DNA regions relative to inter-nucleosomal DNA. However, the average methylation rate of the GCG motif was higher within nucleosomal footprint DNA than within nucleosomal flanking linker regions, with a ratio of 1.13. In contrast, the CGTT motif was significantly enriched in nucleosomal DNA overall, whereas the AACG motif was significantly depleted (**Table S22**).

**Figure 6.**
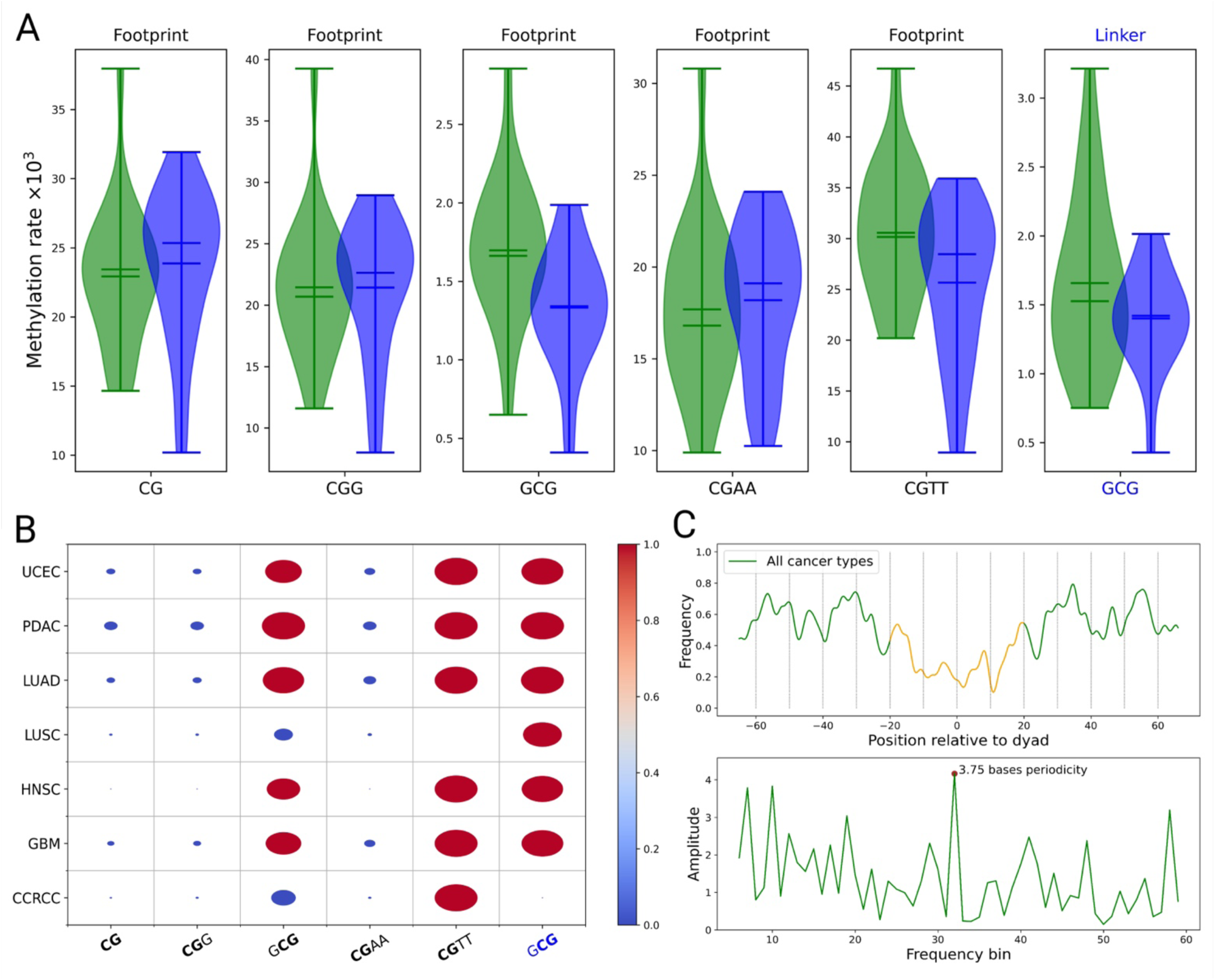
Associations between well-positioned nucleosomes and CpG methylation. (**A**) Distribution of CpG methylation motifs across chromosomes in well-positioned nucleosomes (green) and inter-nucleosomal regions. Methylation motifs associated with flanking linker DNA are shown in blue. (**B**) Enrichment and depletion of identified CpG methylation motifs across cancer types, including clear cell renal cell carcinoma (CCRCC), glioblastoma (GBM), head and neck squamous cell carcinoma (HNSC), lung squamous cell carcinoma (LUSC), lung adenocarcinoma (LUAD), pancreatic ductal adenocarcinoma (PDAC), and uterine corpus endometrial carcinoma (UCEC). Circle size represents the normalized odds ratio, with enrichment and depletion indicated by red and blue circles, respectively. (**C**) Frequency of CpG methylation as a function of distance from the nucleosome dyad across all cancer types, together with periodicity analysis of CpG methylation in nucleosomal DNA. The orange-highlighted region indicates CpG sites within ±20 bp of the dyad position.

Moreover, cancer-type-specific analyses showed that CpG methylation motifs were significantly depleted in nucleosomal footprint regions across multiple cancer types (p< 0.01; **Table S22**). However, we observed cancer-type-specific differences in the GCG methylation motif, which was depleted in lung squamous cell carcinoma (LUSC) and CCRCC but was particularly enriched in LUAD (**Figure 6B**). In flanking linker DNA regions, we found that the GCG methylation motif showed enrichment and depletion patterns consistent with those observed in nucleosomal footprint DNA across cancer types, except in LSC, where GCG was enriched in footprint DNA but depleted in linker regions. Together, these results indicate that CpG methylation is generally depleted within nucleosomal footprint DNA across cancer types, while sequence contexts flanking CpG sites can give rise to distinct methylation motifs that may be selectively enriched within nucleosomal DNA.

Finally, we examined the spatial distribution of CpG methylation sites in the cancer genome relative to nucleosome dyad positions. Across all cancer types, CpG methylation frequencies showed a noisy distribution with pronounced fluctuations (**Figure 6C**). FFT analysis revealed a dominant periodicity of 3.75 bp with the highest amplitude, followed by a secondary periodicity of 10-12 bp at the pan-cancer level. For glioblastoma (GBM) and pancreatic ductal adenocarcinoma (PDAC), the frequency profiles exhibited a dominant periodicity of 12 bases (**Figure S20**), although the signals remained noisy. Despite variability in CpG frequencies, we observed a statistically significant association between CpG methylation and minor-groove orientation across cancer types (**Table S23**). In particular, CpG methylation events associated with CCRCC and PDAC were significantly depleted at minor-groove sites facing outward from histone proteins (odds ratio < 1), whereas other cancer types showed significant enrichment at these sites (**Figure S21**). Additionally, CpG methylation frequency within 20 bases of the dyad position was consistently lower than in surrounding regions. We applied Fisher’s exact test to confirm a significant association between CpG methylation rate and distance from the dyad (**Table S24**), yielding an odds ratio of approximately 0.89 (p ≈ 0). Collectively, these findings suggest that both nucleosome positioning and local DNA structural context modulate CpG methylation patterns in a cancer-type–specific manner.

## Discussion

A longstanding question in understanding the role of nucleosome organization in cancer is how DNA sequence influences nucleosome positioning and, consequently, genomic stability and epigenetic regulation. Although prior studies have characterized sequence preferences associated with nucleosome positioning, they have primarily focused on short dinucleotide patterns or average base composition. Consequently, these approaches provide limited insight into how higher-order sequence context contributes to nucleosome organization and, in turn, relates to somatic mutation accumulation and CpG methylation patterns in cancer genomes. Moreover, despite the critical role of nucleosome organization in modulating DNA repair accessibility and DNA methyltransferase activity, the specific sequence motifs linking nucleosomal regions to somatic mutations and DNA methylation in cancer remain poorly characterized. In this study, we addressed these gaps by developing *DeepND*, an interpretable deep learning framework that identifies informative nucleosome-associated sequence motifs and enables their relationships with somatic mutations and CpG methylation patterns to be investigated across multiple cancer types. Compared with prior studies, our method identified longer informative sequence motifs spanning 2–16 bp, with 4–6 bp motifs representing the most prevalent motif lengths among the identified sequence features. This observation further highlights the limitations of dinucleotide-based models in fully capturing nucleosome sequence grammar, as dinucleotide bendability, groove geometry, and histone–DNA affinity depend strongly on local and higher-order sequence context, which are not simply additive across base-pair steps^24,38^. These sequence features were subsequently associated with cancer-type-specific mutational and methylation patterns, suggesting that higher-order nucleosome-associated sequence context may contribute to differences in the genomic landscapes of cancer.

A comprehensive analysis of convolutional kernels within our *DeepND* model identified 146 motifs associated with nucleosomal footprint DNA and 64 motifs associated with flanking linkers, revealing distinct sequence grammars in nucleosomal and inter-nucleosomal regions. Many of these motifs showed strong positional preferences within nucleosomes, localizing to minor-groove sites facing either inward or outward relative to the histone octamer. Consistent with prior observations, we found that A/T-rich motifs (such as AAAA/TTTT) were generally depleted in nucleosomal footprint DNA. However, when present within footprint regions, these motifs preferentially localized to inward-facing minor-groove sites, where groove narrowing and anisotropic bending can stabilize histone–DNA interactions^39,40^. These positional preferences are consistent with the characteristic periodic organization of nucleosomal DNA. Notably, the AGTACAGGCACAG motif and its complementary sequence were significantly enriched in nucleosomal footprint DNA. This enrichment pattern is consistent with the presence of highly bendable dinucleotide steps, such as TA and CA, together with the absence of extended GC-rich segments, which are known to increase DNA rigidity^41,42^. Analysis of longer convolutional kernels further revealed an intrinsic ∼10–11 bp periodicity, manifested as repeated sequence subregions enriched at minor-groove sites. These longer periodic motifs extend beyond conventional dinucleotide-based descriptions and highlight sequence periodicity as an important feature distinguishing nucleosomal from inter-nucleosomal DNA. Moreover, our analyses revealed significant associations between nucleosome-associated sequence motifs and somatic mutation patterns. At the SNV level, nucleosomal DNA exhibited clear mutation-type-specific biases, most notably an enrichment of C>T transitions across nucleosome classes. C>T mutations frequently arise from spontaneous cytosine deamination, a chemically driven process whose persistence and conversion into somatic mutations depend on DNA accessibility and repair efficiency. DNA wrapped around histones, particularly at inward-facing minor-groove sites, has reduced accessibility to repair machinery, potentially allowing deamination-associated lesions to persist^13^.

Importantly, we did not observe uniform mutation biases, as mutational patterns were strongly dependent on local DNA sequence context^43^. Consistent with this observation, [C>T]G mutations were significantly enriched within nucleosomal footprint DNA and showed significant associations with multiple cancer types. These findings align with previous studies demonstrating that mutation density is strongly influenced by chromatin organization and epigenetic factors, including histone modifications such as H3K9me3, which have been associated with substantial variation in genome-wide mutation rates^44^. Moreover, CpG methylation provides a plausible mechanistic link between nucleosome structure and C>T enrichment in cancer genomes. In particular, methylated cytosines at CpG dinucleotides undergo spontaneous hydrolytic deamination at an elevated rate, converting them to thymine and thereby contributing to C>T mutations. Reduced accessibility of DNA repair machinery within nucleosomal DNA may therefore enhance the persistence of CpG-associated mutations, contributing to the observed enrichment of C>T transitions.

Our findings also highlight that nucleosome-associated mutational biases are cancer-type-specific rather than universal. Many of the mutational motifs we identified here were significantly depleted across cancer types, potentially reflecting differences in mutational signature composition, DNA damage mechanisms, CpG density, epigenetic state, or nucleosome organization^43,45–47^. Consistent with the role of nucleosome organization in cancer mutagenesis, we observed a pronounced ∼10-bp periodicity in the mutation-frequency profiles of LUAD and UCEC, corresponding to the rotational positioning of DNA around the histone octamer. The counterphase oscillation between LUAD and UCEC suggests that distinct mutational processes differ in their sensitivity to nucleosomal rotational geometry. Specifically, mutations in LUAD were preferentially enriched at outward-facing minor-groove sites, whereas those in UCEC were enriched at inward-facing minor-groove sites. In contrast, most other cancer types showed weaker or absent periodic signals, suggesting that the influence of nucleosome organization on mutational patterns may be particularly pronounced in these tumor contexts.

Integrating nucleosome motif-specific mutational profiles with COSMIC mutational signatures further supported these observations and provided a sequence-level view of how established mutational processes are represented within nucleosome-associated contexts. For instance, motif-restricted mutational profiles for LUSC and CCRCC cohorts closely matched SBS4 and SBS94, respectively, consistent with the mutational processes previously reported in these cohorts^48^. Similarly, the motif-restricted mutational profile for the UCEC cohort corresponded to the mismatch-repair-deficiency signatures SBS6 and SBS15. The co-occurrence of these repair-associated signatures with inward-facing minor-groove mutational periodicity is consistent with mismatch repair being sensitive to accessibility constraints imposed by nucleosome organization, although our analyses establish association rather than mechanism. In the LUAD cohort, motif-restricted profiles showed strong overall similarity to the reference signatures. However, this similarity was distributed across several C>A–dominated signatures that could not be resolved from one another within motif-restricted sequence space. Overall, since motif-restricted profiles are drawn from compositionally selected sequence contexts, these correspondences likely indicate consistency with cohort-wide mutational processes rather than evidence that particular signatures act preferentially within nucleosomal DNA.

Finally, we identified five informative methylation motifs within nucleosomal footprint DNA and one within flanking linker DNA, providing sequence-level insights into how nucleosome organization relates to the methylation landscape in cancer. While previous studies have reported conflicting observations regarding CpG methylation enrichment or depletion in nucleosomal DNA^49–51^, our results revealed consistent depletion of CpG methylation within well-positioned nucleosomes. This observation is consistent with reduced accessibility of these regions to DNMT enzymes, as DNMT-mediated methylation requires access to the target cytosine and base flipping, which may be hindered when DNA is tightly wrapped around the histone octamer. Importantly, our findings indicate that CpG methylation patterns are strongly associated with local DNA sequence context, as nucleotides flanking CpG sites can give rise to distinct methylation motifs within nucleosomal DNA^52^. For instance, we observed an enrichment of GCG motifs in well-positioned nucleosomes. Consistent with prior reports, we found significant depletion and enrichment of CpG methylation within the CGAA and CGTT motifs in nucleosomal DNA, respectively. Despite being reverse complements, AA and TT dinucleotide methylation motifs can exhibit distinct structural and mechanical properties that influence DNA deformability and rotational positioning on the nucleosome surface^53^. These findings are consistent with prior observations that DNMT enzymes may preferentially methylate CpG sites in outward-facing minor grooves of the histone octamer, where CGTT motifs are enriched and DNA accessibility is higher. In contrast, CGAA motifs preferentially occupy inward-facing minor-groove orientations, which may reduce accessibility of these CpG sites to DNMT enzymes and contribute to the observed methylation depletion within well-positioned nucleosomes.

We note that our results do not imply that the identified sequence motifs are the sole determinants of nucleosome positioning or that they exclusively drive cancer-associated mutation and methylation patterns. Rather, these motifs likely capture statistical patterns that reflect recurring nucleosomal structural and biochemical constraints underlying chromatin organization. Importantly, our findings support a model in which nucleosome positioning, somatic mutational processes, and DNA methylation interact dynamically and may exert reciprocal influences. For instance, DNA methylation can promote the accumulation of somatic mutations, while mutational events may, in turn, alter local methylation landscapes. Both processes are likely constrained by nucleosome organization. These interdependent relationships warrant further investigation to clarify how sequence context, nucleosome positioning, and mutational signatures collectively contribute to cancer-type-specific genomic patterns. Despite these complexities, our machine learning approach provides a unified framework for investigating this interplay at the sequence level. Overall, our findings support the existence of a nucleosome-associated sequence grammar that may contribute to nucleosomal organization and its associations with genomic stability and epigenetic regulation, thereby providing a potential sequence-level basis for cancer-type-specific mutational and methylation patterns.

## Materials and methods

The *DeepND* framework comprises three main steps: (i) identification of nucleosome dyads from MNase-seq data, (ii) training a classifier to distinguish nucleosomal from inter-nucleosomal DNA, and (iii) interpretation of the learned convolutional kernels (Figure 1).

### Identification of representative dyad positions from MNase-seq data

A variety of experimental methods have been developed to generate genome-wide nucleosome positioning profiles^54–56^. In the present study, we used high-coverage paired-end MNase-seq data (GSE36979) from seven human lymphoblastoid cell lines, as these data provide a more precise and reliable means of identifying nucleosome positions than other experimental approaches^57^. In the MNase-seq method, the micrococcal nuclease (MNase) enzyme preferentially cuts DNA at linker regions, digesting chromatin into nucleosome-sized fragments^58,59^. DNA fragments of different lengths are then released from histone proteins. Among these fragments, 147 bp fragments are the most abundant and are generally considered to represent nucleosomal DNA, while shorter or longer fragments tend to result from over- or under-digestion, potentially compromising the accuracy of nucleosome positioning^60^. Therefore, we mapped the 147 bp fragments to the human reference genome and generated a nucleosome occupancy profile. In this profile, the x-axis represents genomic coordinates, and the y-axis represents the number of fragment midpoints at that coordinate (dyad count). To identify representative dyad positions, we applied a previously developed approach comprising three main steps^61^. First, the nucleosome occupancy profile is smoothed (Eq. 1):

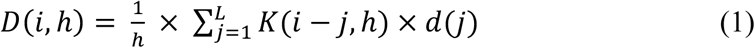

Here, *d(j)* denotes the dyad count at position *j*, and *L* represents the chromosome length. The bandwidth parameter *h* was set to 15, consistent with previous studies reporting that smaller bandwidth values can improve positional accuracy. *K* denotes the tri-weight kernel function (Eq. 2):

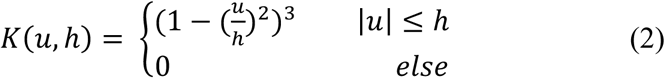

Second, the local maxima of the smoothed nucleosome occupancy profile were identified using the bwa-tool software library, with the minimum distance between two adjacent local maxima set to 150 bases^62^. Finally, 60-bp windows centered on each local maximum were examined, and the genomic coordinate with the highest dyad count within each window was selected as the representative dyad position. When multiple coordinates had the maximum dyad count, the coordinate closest to the window center was selected as the representative dyad position. Overall, we identified the genomic coordinates of more than 12 million nucleosomes across the human genome.

In this study, we separately identified representative dyad positions for both the hg19 and hg38 reference genomes for two main reasons. First, different data resources provide methylation and mutation data based on either hg19 or hg38. Therefore, we needed to map nucleosome positioning to both reference genomes and examine the occurrence of mutations and methylation events within them. Second, to systematically evaluate and compare our machine learning models with existing methods, we used the same conditions as in previous studies that utilized an older reference genome build (hg19). However, we note that the reference genome version does not affect the training and testing processes, as our training data (described in the next section) consists of DNA sequences that are identical across the two reference genomes, even though their genomic coordinates differ.

### Generation of nucleosomal and inter-nucleosomal DNA classification model

Briefly, we partitioned the reference genome into non-overlapping bins and classified them as nucleosomal or inter-nucleosomal regions. These regions were grouped according to the dyad identification procedure described above. We retained only bins within non-repetitive genomic regions to avoid inflated classification performance. We also excluded overlapping sequence bins between the two groups to ensure the identification of discriminative sequence motifs. Each data instance comprised 201 bp, including a 147-bp nucleosomal DNA footprint flanked by 27-bp linker regions on both sides. For the positive set (nucleosomal DNA), we centered sequences at the dyad position and extended them 100 bp upstream and downstream. In contrast, negative set (inter-nucleosomal DNA) sequences were centered on the midpoint between two consecutive nucleosomes separated by at least 400 bp, with each sequence extending 100 bp in each direction. In addition, the reverse complementary strand (5′–3′ orientation) was included for each sequence. In total, each group contained 115,640 instances.

We encoded each data instance using a one-hot representation to train a deep residual network (ResNet) to classify input sequence segments as nucleosomal or inter-nucleosomal DNA. In this architecture, all convolutional kernels shared the same dimensions, and identity skip connections added the input of the (*i*−1)^th^ residual block to the output of the *i*^th^ residual block. Each residual block comprised five convolutional layers. Accordingly, the output of the *c^th^* and *u^th^* cell of the kernel *f* in the *i*^th^ convolutional layer, denoted as 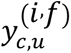, is defined as shown in Eq. (3):

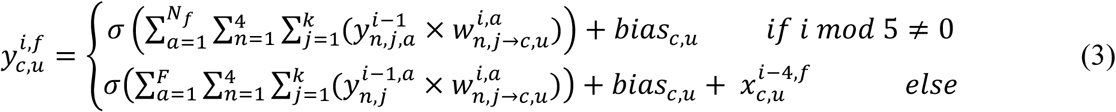

where *σ* denotes the sigmoid activation function, and *N_f_* represents the total number of kernels (filters) in a convolutional layer. Also, 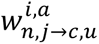 denotes the weight connecting the element (*n*, *j*) of filter *a* in the (*i* − 1)*^th^* layer to the element (*c*, *u*) of filter *f* in the *i^th^* layer. Similarly, 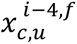 denotes the corresponding input element from the (*i* − 4)*^th^* layer that is added through the skip connection.

During each training phase, a snapshot of the model parameters was saved. When the loss on the training set decreased but the loss on the test set either increased or remained unchanged, the backpropagation update was halted. Instead, the most recent non-overfitted parameter snapshot was restored, and up to 2% of the trainable parameters were randomly selected and updated using Eq. 4.

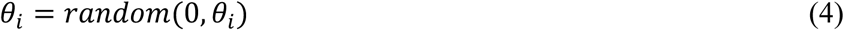

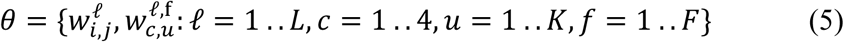

Here, *θ* denotes the set of all trainable parameters of the model, including both kernel weights and fully connected layer weights. The function *random* generates a floating-point value uniformly sampled between 0 and a specified upper bound. The terms 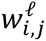 and 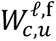 represent the weight connecting the *i^th^* neuron of layer *ℓ* to the *j^th^* neuron of layer *ℓ* + 1, and the kernel weight corresponding to nucleotide channel *c*, kernel position *u*, and filter *f*, respectively. In addition, *K* denotes the kernel length, *F* denotes the number of filters in a convolution layer, and *L* refers to the total number of layers in the model.

### Identification of nucleosomal-DNA related sequence motifs

We executed our framework (Figure 2B) 29 times, using convolutional kernel lengths from 2 to 30 bp to identify sequence motifs within this range. After each training run, we extracted the kernels from the first convolutional layer to detect sequence motifs. We primarily focused on the first layer because it captures the primary patterns within DNA sequences, whereas deeper layers learn higher-order combinations of these motifs^63^. In the run *R*, we set the length of all kernels to 4 × (*R* + 1), where the factor of four corresponds to the four nucleotide channels (A, T, C, and G). To construct sequence motifs from a given kernel, we examined each column matrix of the kernel matrix and selected the nucleotide at the position *u* whose channel exhibited the **maximum weight** at that position (Eq. 6). This allowed us to convert each kernel into a discrete DNA sequence motif representing the most informative pattern learned by the model.

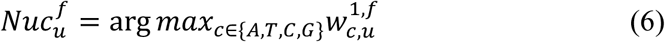

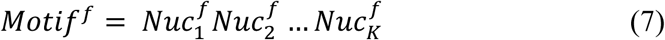

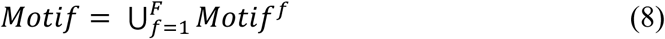

Here, Motif^*(f)*^consisting of *K* nucleotides (Nuc), denotes the consensus sequence learned by the filter *f*, where *K* corresponds to the kernel length. Accordingly, Motif represents the set of all sequence motifs learned by *DeepND*.

We applied Fisher’s exact test to evaluate associations between candidate sequence motifs and nucleosomal and inter-nucleosomal DNA groups^64^. We defined informative sequence motifs as those that showed statistically significant enrichment or depletion after multiple-testing correction. Specifically, we retained motifs with *p* < 0 and classified those with odds ratios > 1 as enriched and those with odds ratios < 1 as depleted. Furthermore, we prioritized the most informative sequence motifs by applying our previously developed wrapper-based feature selection algorithm to a dataset in which each DNA sequence was represented as a feature vector^65^. Each feature vector quantified the total occurrences of each informative motif within nucleosomal and inter-nucleosomal DNA sequences. Our wrapper feature selection method combines the *Trader* metaheuristic algorithm with a support vector machine (SVM) classifier to discriminate between these sequence classes. Briefly, we start with an initial population of randomly generated candidate feature subsets. Next, we calculate the score of each candidate based on the classification accuracy of the SVM model using the selected features. The algorithm then iteratively refines candidate solutions using the three operators of the *Trader* algorithm, which enables effective exploration of the feature space while avoiding fitness-decreasing modifications. Compared with other metaheuristic feature selection methods, our approach facilitates escaping local optima and improves the model’s overall performance. We previously demonstrated that this method outperforms widely used feature selection algorithms across standard evaluation metrics^66^.

### Identification of mutational and methylation sequence motifs

We obtained somatic mutation and CpG methylation coordinates mapped to GRCh38 (at the patient level) from the Clinical Proteomic Tumor Analysis Consortium (CPTAC) study. Cancer types analyzed in this study included clear cell renal cell carcinoma (CCRCC), glioblastoma multiforme (GBM), head and neck squamous cell carcinoma (HNSCC), lung squamous cell carcinoma (LUSC), lung adenocarcinoma (LUAD), pancreatic ductal adenocarcinoma (PDAC), and uterine corpus endometrial carcinoma (UCEC). We included CPTAC cohorts in our analyses due to the availability of whole-genome and methylation data for each patient. We downloaded each dataset using the Genomic Data Commons (GDC) API.

For mutational enrichment analyses, we included only Mutect2-based mutation calls that passed quality-control filters according to established statistical criteria. We then transformed the identified informative sequence motifs into a comprehensive set of putative nucleosomal-DNA mutational motifs (Eq. 9). For this, we mutated each nucleotide position within an informative sequence motif to each of the three alternative nucleotides, thereby enumerating all possible single-nucleotide substitution variants associated with each motif (Eq. 10).

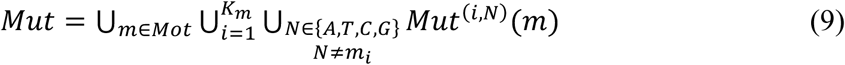

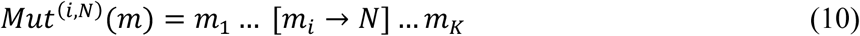

where *m* denotes a sequence motif and *K_m_* represents its length. The function *Mut*^*(i,N)*^(*m*) substitutes the *i^th^* nucleotide of *m* with nucleotide *N*. Also, *Mot* denotes the set of all informative sequence motifs.

We next analyzed the resulting list of potential mutational motifs (*Mut*) using Fisher’s exact test to assess their association with nucleosomal and inter-nucleosomal DNA. We retained mutational motifs with *p* < 0.01 and defined them as informative, while discarding the others. We classified motifs with an odds ratio greater than 1 as enriched and those with an odds ratio less than 1 as depleted in nucleosomal DNA.

For DNA methylation analysis, CpG sites were classified by beta value, with values greater than 0.6 indicating high methylation and lower values indicating reduced methylation. This threshold-based stratification was used to focus on strongly methylated loci. To identify methylation-associated sequence motifs, we applied an analogous procedure. Specifically, we filtered the set of informative motifs to retain only those containing a CpG dinucleotide, defined as motifs in which a guanine (G) immediately follows a cytosine (C) (Eq. 11):

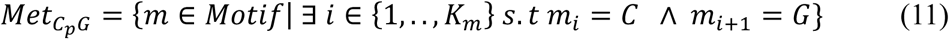

where *Met_CpG_* refers to a set of potential methylation motifs and *K_m_* represents the length of a sequence motif (*m*).

Similar to mutational motifs, we also analyzed the resulting list of potential methylation motifs (Met*_CpG_*) by applying Fisher’s exact test to quantify their association with nucleosomal versus inter-nucleosomal DNA. We retained motifs with p < 0.01 and defined them as informative methylation motifs, discarding those that did not meet this significance threshold. We classified motifs with odds ratios greater than 1 as enriched and those with odds ratios less than 1 as depleted in nucleosomal DNA, using the same criteria as in mutational motif analysis.

### Determination of the association between mutational motifs and mutational signatures

We investigated whether mutational signatures observed across cancer types are significantly associated with the identified mutational sequence motifs. To this end, we generated mutational signatures for each cancer type through a multi-step procedure. First, Fisher’s exact test was applied to identify mutational motifs significantly enriched in each cancer type (p < 0.01). Second, for each cancer type *m*, we constructed a mutational profile based on the 96 trinucleotide substitution classes defined by the COSMIC framework (Eq. 12), restricted to the identified enriched mutational motifs for that cancer type.

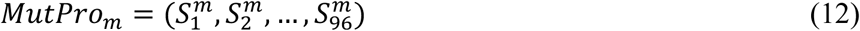

where 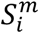 represents the contribution of the *i*^th^ trinucleotide substitution pattern in cancer type *m*, following the standard COSMIC ordering. Each component 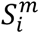 is defined as:

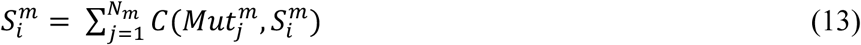

where *N_m_* denotes the total number of mutational sequence motifs significantly enriched in cancer type *m*, and 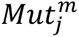 denotes the *j*^th^ enriched mutational motif for that cancer type. The function *C*(⋅) is defined as an indicator function: 1, *if S^m^ is contained wit*ℎ*in Mut^m^*

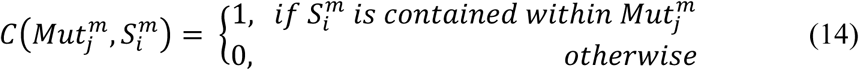

Finally, we quantified the similarity between cancer-specific mutational profiles and reference profiles from the COSMIC database using the Pearson correlation coefficient (PCC) and cosine similarity. These metrics assessed concordance between reconstructed mutational signatures and established COSMIC signatures across cancer types. In downstream analysis, we further interpreted these mutational signatures in the context of epigenetic features, including DNA methylation patterns and nucleosome positioning landscapes, to investigate their potential association with cancer-specific mutational processes.

## Source code and data availability

The authors have implemented their algorithm in Python and made it available on GitHub at https://github.com/MasoudiYosef/DeepND. The high-coverage paired-end MNase-seq data from seven human lymphoblastoid cell lines used for nucleosome positioning analysis were obtained from the NCBI Gene Expression Omnibus (GEO) under accession number GSE36979 (https://www.ncbi.nlm.nih.gov/geo/query/acc.cgi?acc=GSE36979). We accessed somatic mutation and DNA methylation data from the Clinical Proteomic Tumor Analysis Consortium (CPTAC) through the NCI Genomic Data Commons (GDC). These are controlled-access data; access was granted under dbGaP accession (phs001287). The COSMIC mutational signatures were obtained from the COSMIC database.

## Acknowledgments

We acknowledge support from the Digital Research Alliance of Canada for providing necessary computing resources to perform this study.

## References

1. Zhou, K., Gaullier, G. & Luger, K. Nucleosome structure and dynamics are coming of age. Nat Struct Mol Biol 26, 3–13 (2019).

2. Zhou, B.-R. et al. Distinct Structures and Dynamics of Chromatosomes with Different Human Linker Histone Isoforms. Mol Cell 81, 166–182.e6 (2021).

3. Li, S., Peng, Y. & Panchenko, A. R. DNA methylation: Precise modulation of chromatin structure and dynamics. Curr Opin Struct Biol 75, 102430 (2022).

4. Segal, E. et al. A genomic code for nucleosome positioning. Nature 442, 772–778 (2006).

5. Shtumpf, M., Piroeva, K. V., Agrawal, S. P., Jacob, D. R. & Teif, V. B. NucPosDB: a database of nucleosome positioning in vivo and nucleosomics of cell-free DNA. Chromosoma 131, 19–28 (2022).

6. Di Gangi, M., Lo Bosco, G. & Rizzo, R. Deep learning architectures for prediction of nucleosome positioning from sequences data. BMC Bioinformatics 19, 418 (2018).

7. Masoudi-Sobhanzadeh, Y., et al. Building Nucleosome Positioning Maps: Discovering Hidden Gems. WIREs Comput Mol Sci 15, e70029 (2025).

8. Li, C. & Luscombe, N. M. Nucleosome positioning stability is a modulator of germline mutation rate variation across the human genome. Nat Commun 11, 1363 (2020).

9. Espiritu, D., Gribkova, A. K., Gupta, S., Shaytan, A. K. & Panchenko, A. R. Molecular Mechanisms of Oncogenesis through the Lens of Nucleosomes and Histones. J. Phys. Chem. B 125, 3963–3976 (2021).

10. Guintini, L., Charton, R., Peyresaubes, F., Thoma, F. & Conconi, A. Nucleosome positioning, nucleotide excision repair and photoreactivation in Saccharomyces cerevisiae. DNA Repair 36, 98–104 (2015).

11. Ostroverkhova, D. et al. DNA polymerase ε and δ variants drive mutagenesis in polypurine tracts in human tumors. Cell Reports 43, 113655 (2024).

12. Mao, P., Smerdon, M. J., Roberts, S. A. & Wyrick, J. J. Asymmetric repair of UV damage in nucleosomes imposes a DNA strand polarity on somatic mutations in skin cancer. Genome Res. 30, 12–21 (2020).

13. Pich, O. et al. Somatic and Germline Mutation Periodicity Follow the Orientation of the DNA Minor Groove around Nucleosomes. Cell 175, 1074–1087.e18 (2018).

14. Portella, G., Battistini, F. & Orozco, M. Understanding the Connection between Epigenetic DNA Methylation and Nucleosome Positioning from Computer Simulations. PLoS Comput Biol 9, e1003354 (2013).

15. Heyn, H. & Esteller, M. DNA methylation profiling in the clinic: applications and challenges. Nat Rev Genet 13, 679–692 (2012).

16. Chodavarapu, R. K. et al. Relationship between nucleosome positioning and DNA methylation. Nature 466, 388–392 (2010).

17. George, K., Neininger, K., Schmitz, A. E., Walter, J. & Helms, V. Structural DNMT-nucleosome contacts are related to DNA methylation patterns. Epigenetics & Chromatin 18, 59 (2025).

18. Chen, W. et al. Improved nucleosome-positioning algorithm iNPS for accurate nucleosome positioning from sequencing data. Nat Commun 5, 4909 (2014).

19. Brogaard, K., Xi, L., Wang, J.-P. & Widom, J. A map of nucleosome positions in yeast at base-pair resolution. Nature 486, 496–501 (2012).

20. Snyder, M. W., Kircher, M., Hill, A. J., Daza, R. M. & Shendure, J. Cell-free DNA Comprises an In Vivo Nucleosome Footprint that Informs Its Tissues-Of-Origin. Cell 164, 57–68 (2016).

21. De Sarkar, N. et al. Nucleosome Patterns in Circulating Tumor DNA Reveal Transcriptional Regulation of Advanced Prostate Cancer Phenotypes. Cancer Discovery 13, 632–653 (2023).

22. Piroeva, K. V. et al. Nucleosome repositioning in chronic lymphocytic leukemia. Genome Res. 33, 1649–1661 (2023).

23. Underhill, H. R. et al. Fragment Length of Circulating Tumor DNA. PLoS Genet 12, e1006162 (2016).

24. Masoudi-Sobhanzadeh, Y., Li, S., Peng, Y. & Panchenko, A. R. Interpretable deep residual network uncovers nucleosome positioning and associated features. Nucleic Acids Research 52, 8734–8745 (2024).

25. Travers, A. The location of the linker histone on the nucleosome. Trends in Biochemical Sciences 24, 4–7 (1999).

26. Schones, D. E. et al. Dynamic Regulation of Nucleosome Positioning in the Human Genome. Cell 132, 887–898 (2008).

27. Valouev, A. et al. A high-resolution, nucleosome position map of *C. elegans* reveals a lack of universal sequence-dictated positioning. Genome Res. 18, 1051–1063 (2008).

28. Mavrich, T. N. et al. A barrier nucleosome model for statistical positioning of nucleosomes throughout the yeast genome. Genome Res. 18, 1073–1083 (2008).

29. Zhang, J., Peng, W. & Wang, L. LeNup: learning nucleosome positioning from DNA sequences with improved convolutional neural networks. Bioinformatics 34, 1705–1712 (2018).

30. Guo, S.-H. et al. iNuc-PseKNC: a sequence-based predictor for predicting nucleosome positioning in genomes with pseudo k-tuple nucleotide composition. Bioinformatics 30, 1522–1529 (2014).

31. Tahir, M. & Hayat, M. iNuc-STNC: a sequence-based predictor for identification of nucleosome positioning in genomes by extending the concept of SAAC and Chou’s PseAAC. Mol. BioSyst. 12, 2587–2593 (2016).

32. Tahir, M., Hayat, M. & Khan, S. A. iNuc-ext-PseTNC: an efficient ensemble model for identification of nucleosome positioning by extending the concept of Chou’s PseAAC to pseudo-tri-nucleotide composition. Mol Genet Genomics 294, 199–210 (2019).

33. Chen, L. et al. Nucleosome spacing can fine-tune higher-order chromatin assembly. Nat Commun 16, 6315 (2025).

34. Rohs, R. et al. The role of DNA shape in protein–DNA recognition. Nature 461, 1248–1253 (2009).

35. Jiang, C. & Pugh, B. F. Nucleosome positioning and gene regulation: advances through genomics. Nat Rev Genet 10, 161–172 (2009).

36. Bernstein, N. et al. Analysis of somatic mutations in whole blood from 200,618 individuals identifies pervasive positive selection and novel drivers of clonal hematopoiesis. Nat Genet 56, 1147–1155 (2024).

37. López-Cade, I. et al. Comparative Evaluation of Mutect2, Strelka2, and FreeBayes for Somatic SNV Detection in Synthetic and Clinical Whole-Exome Sequencing Data. Biomolecules 15, 1532 (2025).

38. Brukner, I., Sánchez, R., Suck, D. & Pongor, S. Sequence-dependent bending propensity of DNA as revealed by DNase I: parameters for trinucleotides. EMBO J 14, 1812–1818 (1995).

39. Field, Y. et al. Distinct modes of regulation by chromatin encoded through nucleosome positioning signals. PLoS Comput Biol 4, e1000216 (2008).

40. Awazu, A. Prediction of nucleosome positioning by the incorporation of frequencies and distributions of three different nucleotide segment lengths into a general pseudo k-tuple nucleotide composition. Bioinformatics 33, 42–48 (2017).

41. Widom, J. Role of DNA sequence in nucleosome stability and dynamics. Quart. Rev. Biophys. 34, 269–324 (2001).

42. Freeman, G. S., Lequieu, J. P., Hinckley, D. M., Whitmer, J. K. & De Pablo, J. J. DNA Shape Dominates Sequence Affinity in Nucleosome Formation. Phys. Rev. Lett. 113, 168101 (2014).

43. Aggarwala, V. & Voight, B. F. An expanded sequence context model broadly explains variability in polymorphism levels across the human genome. Nat Genet 48, 349–355 (2016).

44. Chen, X. et al. Nucleosomes Suppress Spontaneous Mutations Base-Specifically in Eukaryotes. Science 335, 1235–1238 (2012).

45. Sabarinathan, R., Mularoni, L., Deu-Pons, J., Gonzalez-Perez, A. & López-Bigas, N. Nucleotide excision repair is impaired by binding of transcription factors to DNA. Nature 532, 264–267 (2016).ss

46. Polak, P. et al. Cell-of-origin chromatin organization shapes the mutational landscape of cancer. Nature 518, 360–364 (2015).

47. Schuster-Böckler, B. & Lehner, B. Chromatin organization is a major influence on regional mutation rates in human cancer cells. Nature 488, 504–507 (2012).

48. Alexandrov, L. B. et al. The repertoire of mutational signatures in human cancer. Nature 578, 94–101 (2020).

49. Felle, M. et al. Nucleosomes protect DNA from DNA methylation in vivo and in vitro. Nucleic Acids Res 39, 6956–6969 (2011).

50. Collings, C. K. & Anderson, J. N. Links between DNA methylation and nucleosome occupancy in the human genome. Epigenetics Chromatin 10, 18 (2017).

51. Portela, A. et al. DNA methylation determines nucleosome occupancy in the 5’-CpG islands of tumor suppressor genes. Oncogene 32, 5421–5428 (2013).

52. Mallona, I. et al. Flanking sequence preference modulates de novo DNA methylation in the mouse genome. Nucleic Acids Res 49, 145–157 (2021).

53. Fujii, S., Kono, H., Takenaka, S., Go, N. & Sarai, A. Sequence-dependent DNA deformability studied using molecular dynamics simulations. Nucleic Acids Research 35, 6063–6074 (2007).

54. Chereji, R. V., Bryson, T. D. & Henikoff, S. Quantitative MNase-seq accurately maps nucleosome occupancy levels. Genome Biol 20, 198 (2019).

55. Zhang, Z. & Pugh, B. F. High-Resolution Genome-wide Mapping of the Primary Structure of Chromatin. Cell 144, 175–186 (2011).

56. Dingwall, C., Lomonossoff, G. P. & Laskey, R. A. High sequence specificity of micrococcal nuclease. Nucl Acids Res 9, 2659–2674 (1981).

57. Gaffney, D. J. et al. Controls of Nucleosome Positioning in the Human Genome. PLoS Genet 8, e1003036 (2012).

58. Lim, H.-W. & Iwafuchi, M. Profiling Accessible Chromatin and Nucleosomes in the Mammalian Genome. in DNA-Protein Interactions (ed. Simoes-Costa, M.) vol. 2599 59–68 (Springer US, New York, NY, 2023).

59. Lai, B. et al. Principles of nucleosome organization revealed by single-cell micrococcal nuclease sequencing. Nature 562, 281–285 (2018).

60. Peng, Y. et al. Detection of new pioneer transcription factors as cell-type-specific nucleosome binders. eLife 12, RP88936 (2024).

61. Valouev, A. et al. Determinants of nucleosome organization in primary human cells. Nature 474, 516–520 (2011).

62. Pohl, A. & Beato, M. bwtool: a tool for bigWig files. Bioinformatics 30, 1618–1619 (2014).

63. Angermueller, C., Lee, H. J., Reik, W. & Stegle, O. DeepCpG: accurate prediction of single-cell DNA methylation states using deep learning. Genome Biol 18, 67 (2017).

64. Fisher, R. A. On the Interpretation of χ 2 from Contingency Tables, and the Calculation of P. Journal of the Royal Statistical Society 85, 87 (1922).

65. Masoudi-Sobhanzadeh, Y., Omidi, Y., Amanlou, M. & Masoudi-Nejad, A. Trader as a new optimization algorithm predicts drug-target interactions efficiently. Sci Rep 9, 9348 (2019).

66. Daneshvar, N. H.-N., Masoudi-Sobhanzadeh, Y. & Omidi, Y. A voting-based machine learning approach for classifying biological and clinical datasets. BMC Bioinformatics 24, 140 (2023).

