## supplementary information for "Distinct sequence grammars of nucleosomal and linker DNA shape mutational and methylation landscapes in cancer"

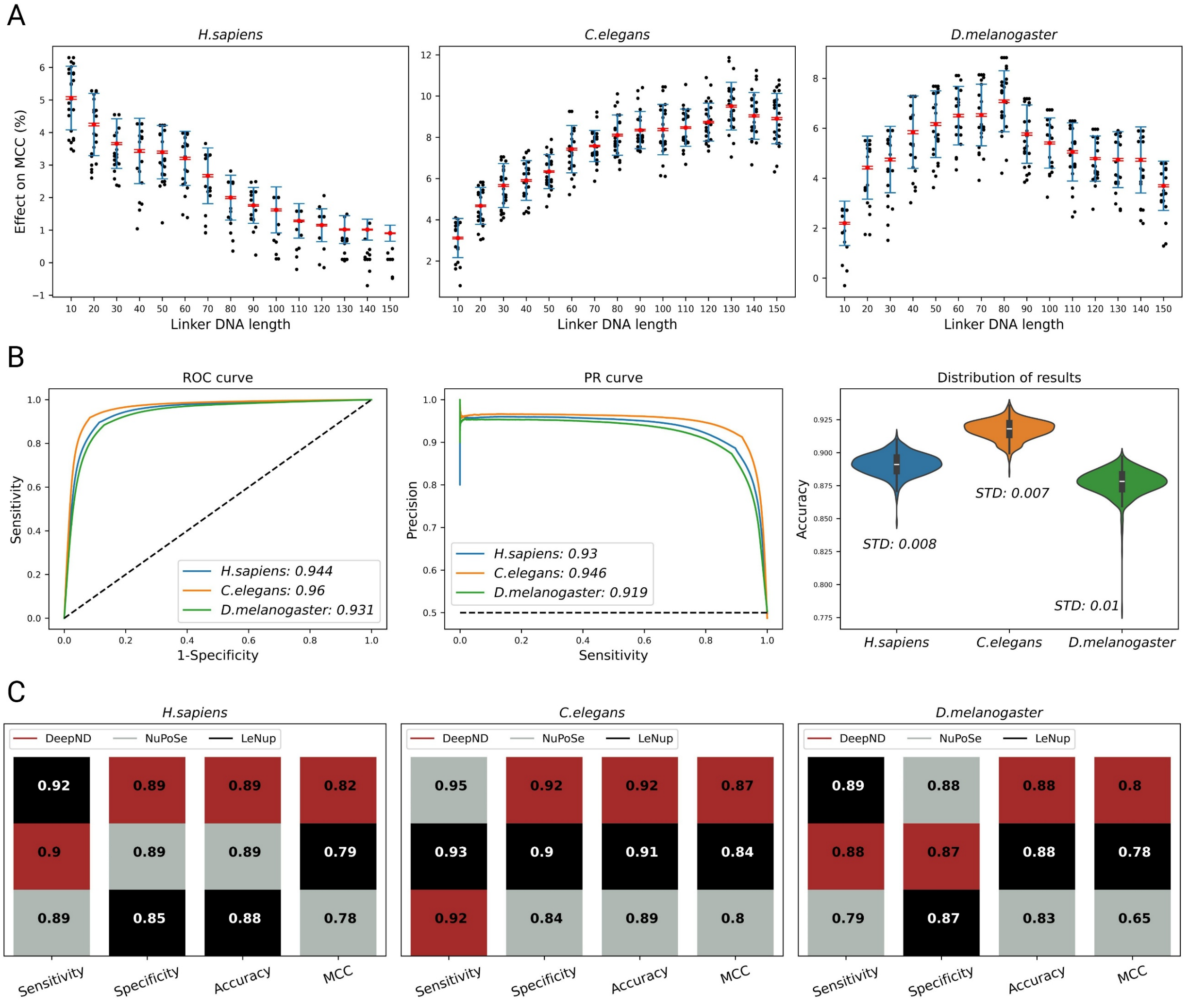


**Figure S1.** Effect of flanking linker DNA length on *DeepND* performance and comparison with other deep learning methods on gold-standard datasets. (A) Inclusion of flanking linker DNA consistently improves model performance. For *H. sapiens*, *C. elegans*, and *D. melanogaster*, performance gains are maximized when the linker DNA length is approximately 10–20 bp, 90–130 bp, and 60–90 bp, respectively. Each point corresponds to a distinct run of *DeepND* on the respective dataset. (B) Evaluation of *DeepND* on three gold-standard datasets using ROC curves, precision–recall (PR) curves, and performance distributions across multiple runs. (C) Comparison of *DeepND* with two recently introduced and high-performing deep learning methods.

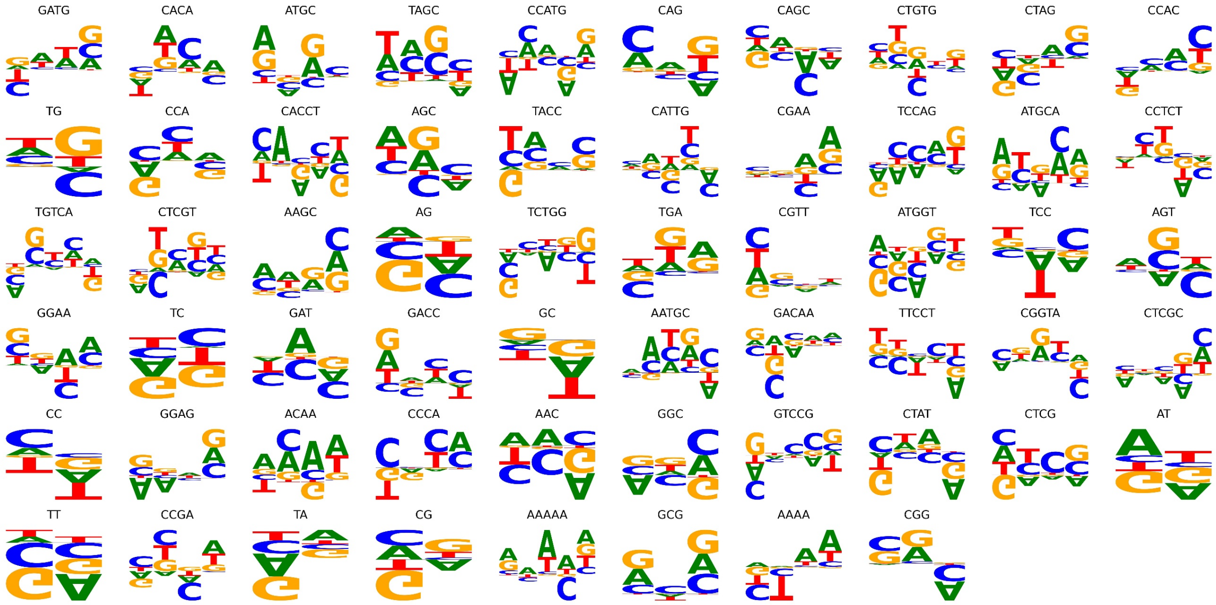


**Figure S2.** Identified sequence features significantly enriched or depleted in nucleosomal footprint DNA. Sequence features with lengths ranging from 2 to 5 bp are shown and are ranked by the odds ratio derived from Fisher's exact test (Table S3).

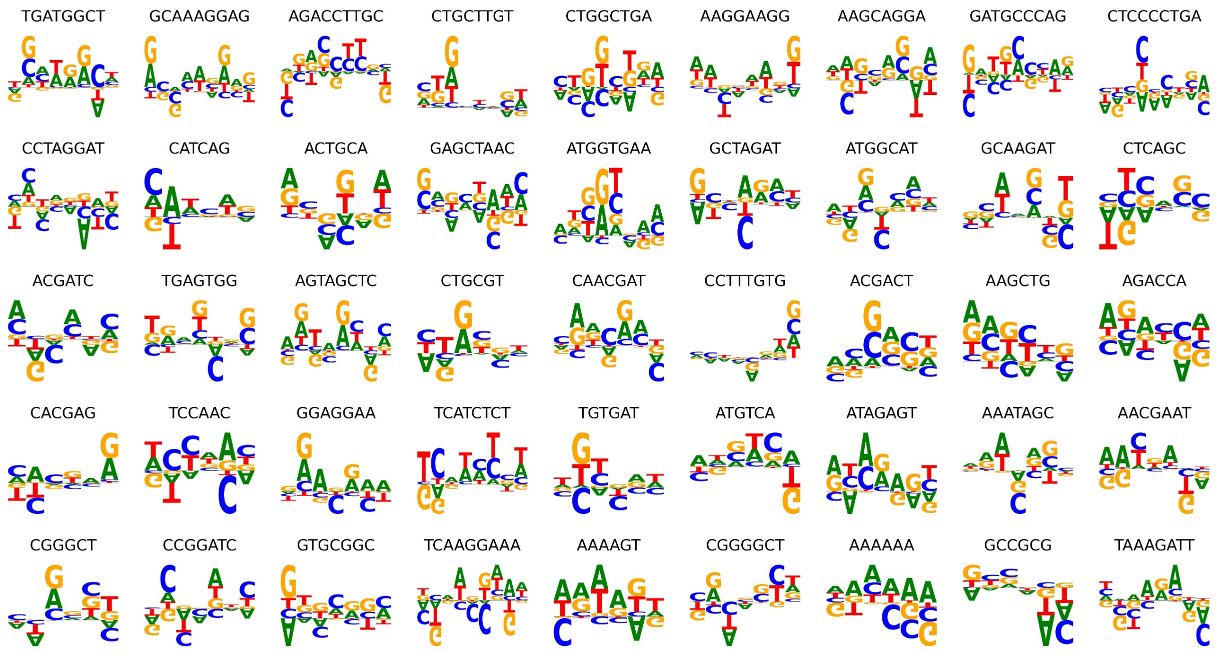


**Figure S3.** Identified sequence features significantly enriched or depleted in nucleosomal footprint DNA. Sequence features with lengths ranging from 6 to 9 bp are shown and are ranked by the odds ratio derived from Fisher's exact test (Table S3).

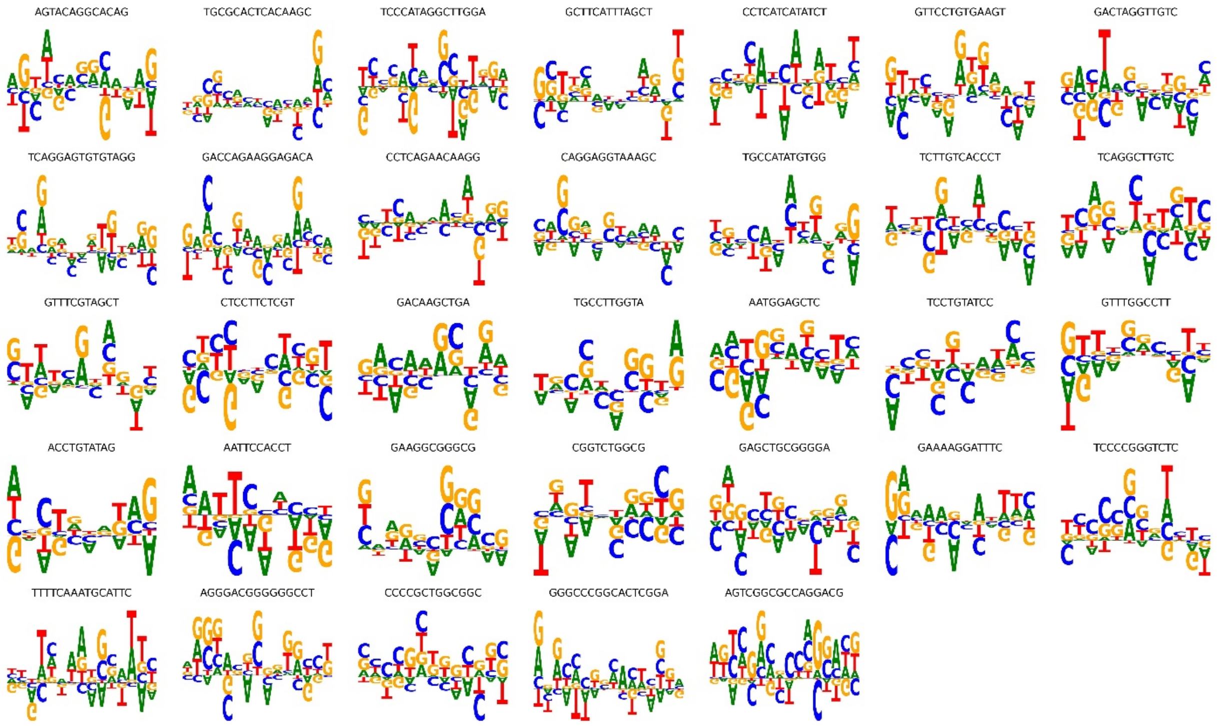


**Figure S4.** Identified sequence features significantly enriched or depleted in nucleosomal footprint DNA. Sequence features with lengths ranging from 10 to 16 bp are shown and are ranked by the odds ratio derived from Fisher's exact test (Table S3).

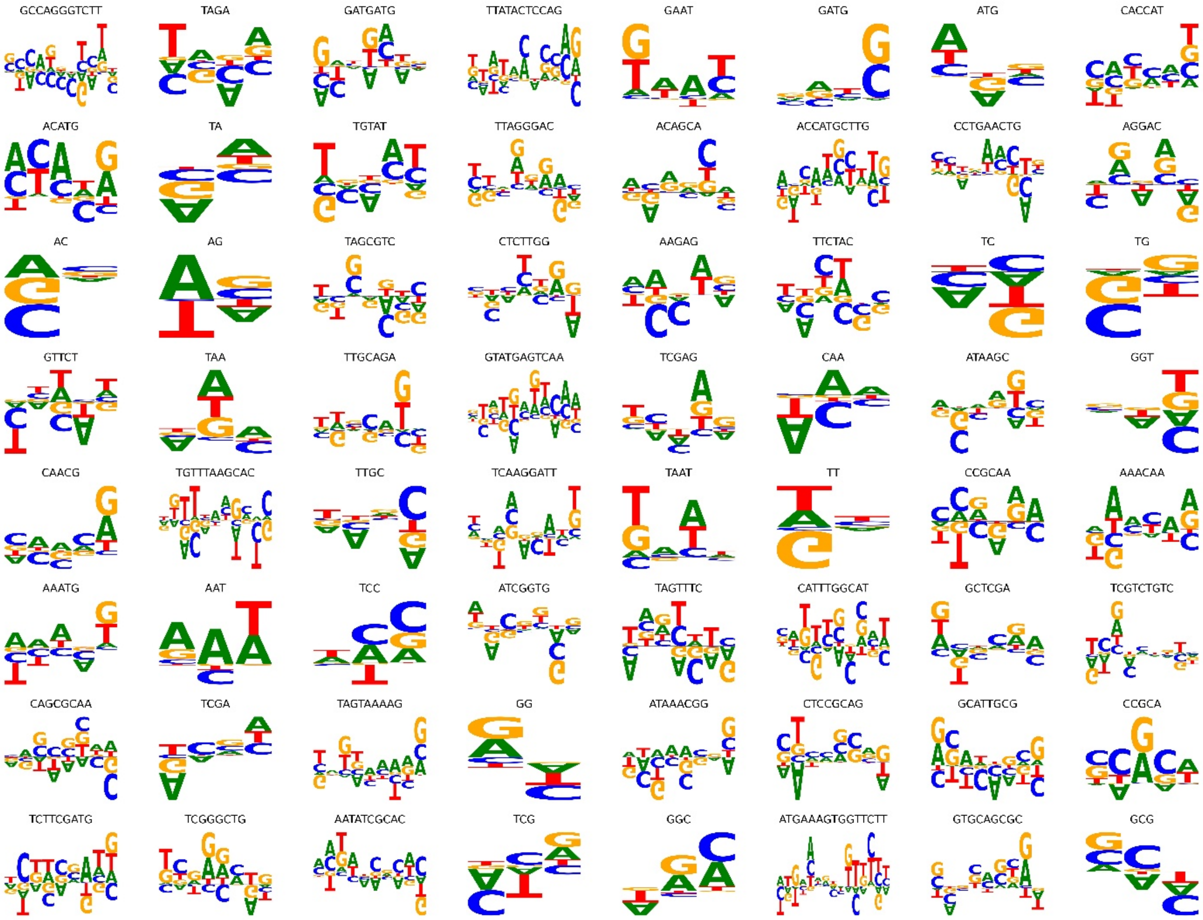


**Figure S5.** Identified sequence features significantly enriched or depleted in nucleosomal flanking linker DNA. Sequence features are ranked by odds ratios derived from Fisher's exact test (Table S4).

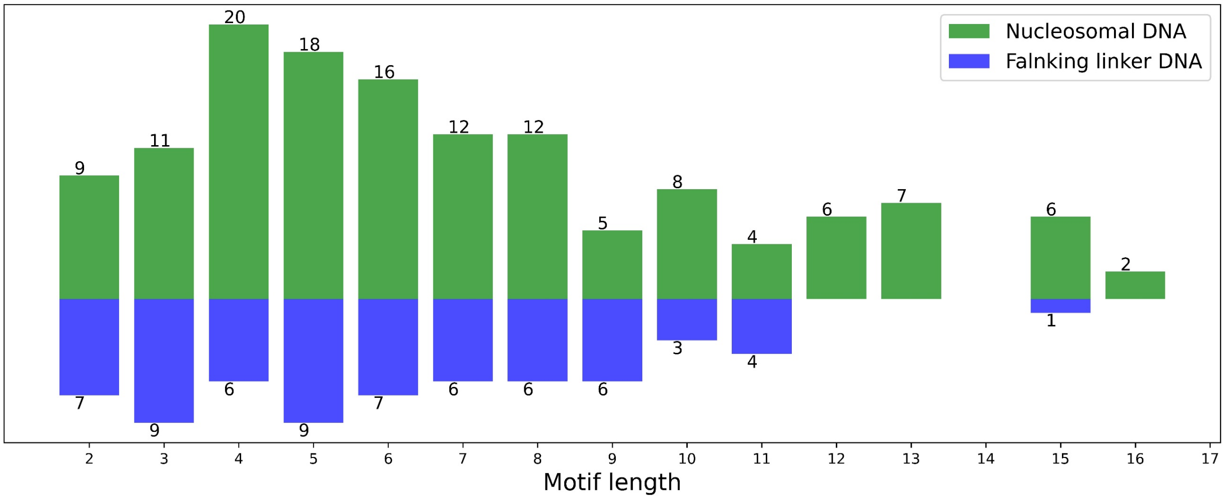


**Figure S6.** Frequency of identified sequence features in nucleosomal footprint (green) and flanking linker DNA (blue) regions. The x-axis represents motif length, and numbers on the bars indicate motif frequencies.

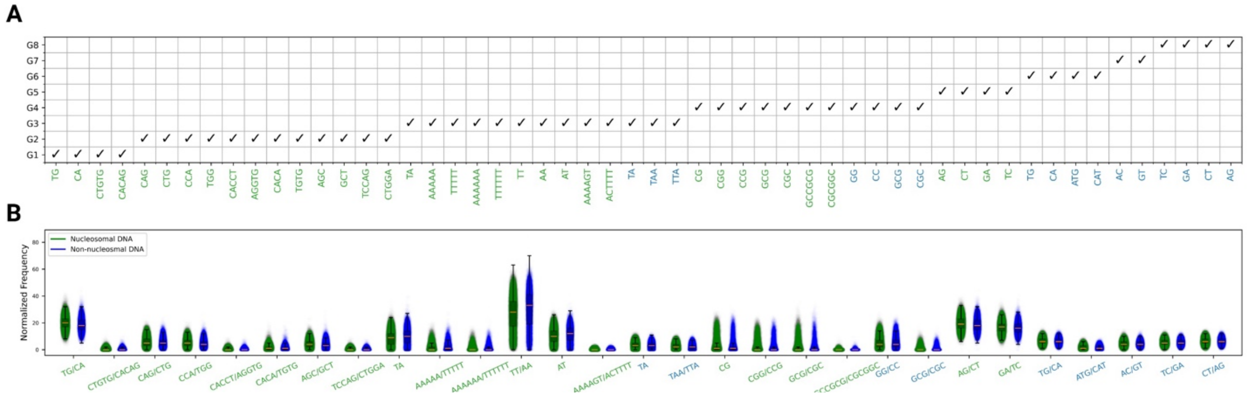


**Figure S7.** An extended set of informative sequence motifs ranked by their impact on the classifier. (A) The extended set of informative sequence motifs grouped by Pearson correlation coefficients (>0.5). (B) Frequencies of the extended group of selected informative sequence motifs in nucleosomal and inter-nucleosomal DNA regions.

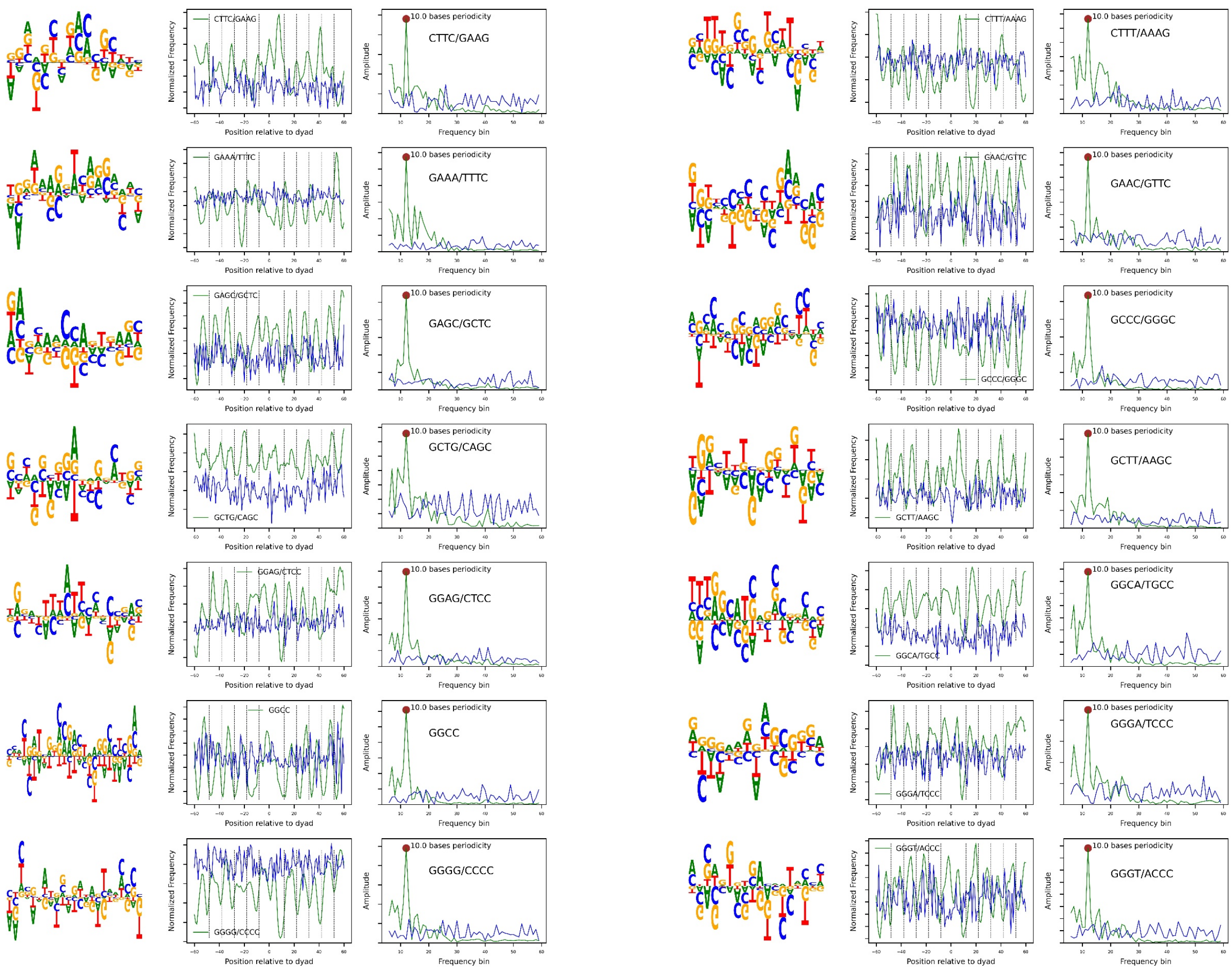


**Figure S8.** Kernels consisting of identical subregions spaced 10–11 bp apart. These subregions are referred to as *periodicity-related sequence motifs*. Long kernels (>15 bp) are visualized using sequence logos, and for each periodicity-related feature, both the frequency in nucleosomal (green) and inter-nucleosomal (blue) DNA sites and the results of fast Fourier transform (FFT) analysis of the frequency are shown. This figure presents some periodicity-related sequence features of length 4 bp.

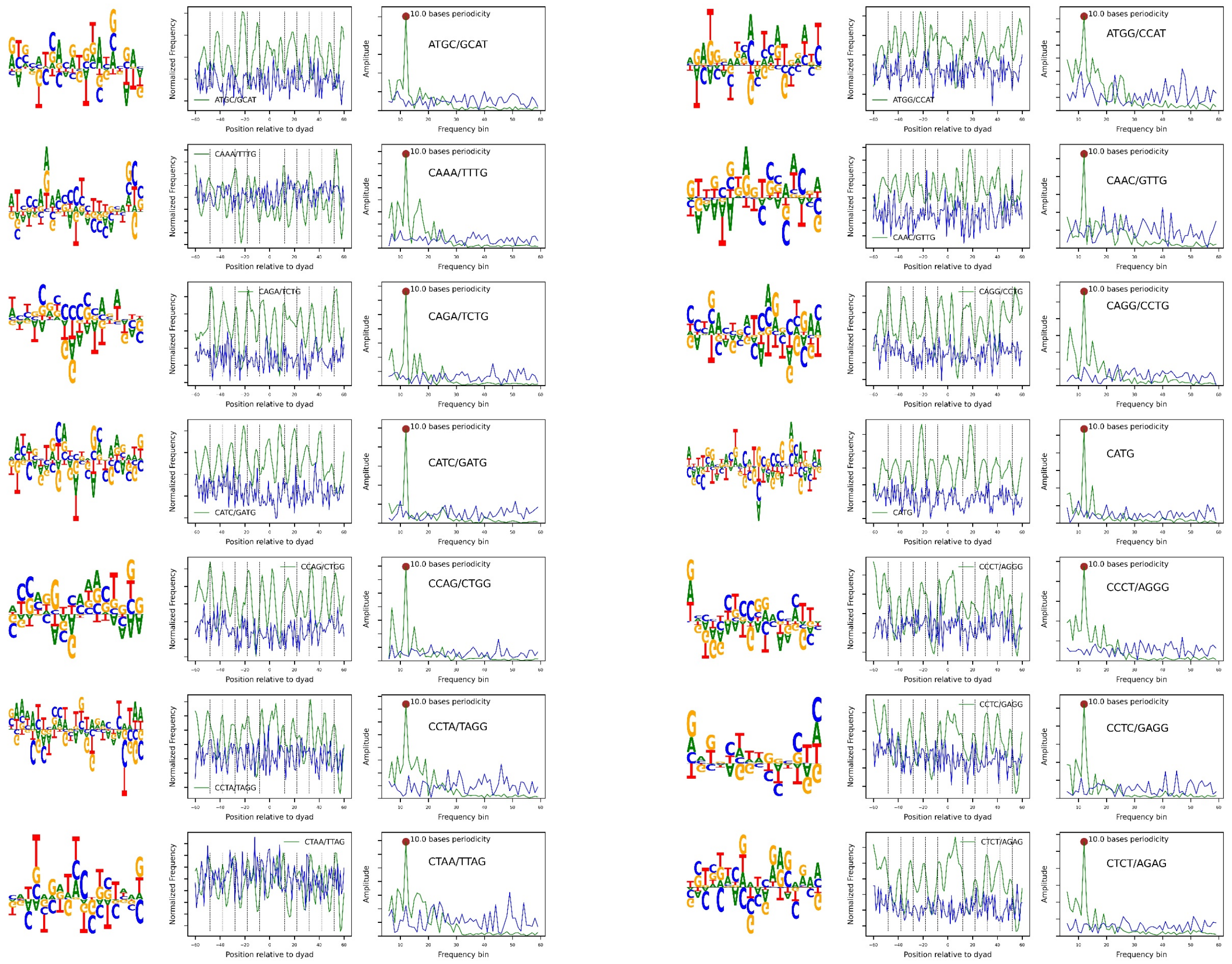


**Figure S9.** Kernels consisting of identical subregions spaced 10–11 bp apart. These subregions are referred to as *periodicity-related sequence motifs*. Long kernels (>15 bp) are visualized using sequence logos, and for each periodicity-related feature, both the frequency in nucleosomal (green) and inter-nucleosomal (blue) DNA sites and the results of fast Fourier transform (FFT) analysis of the frequency are shown. This figure presents some periodicity-related sequence features of length 4 bp.

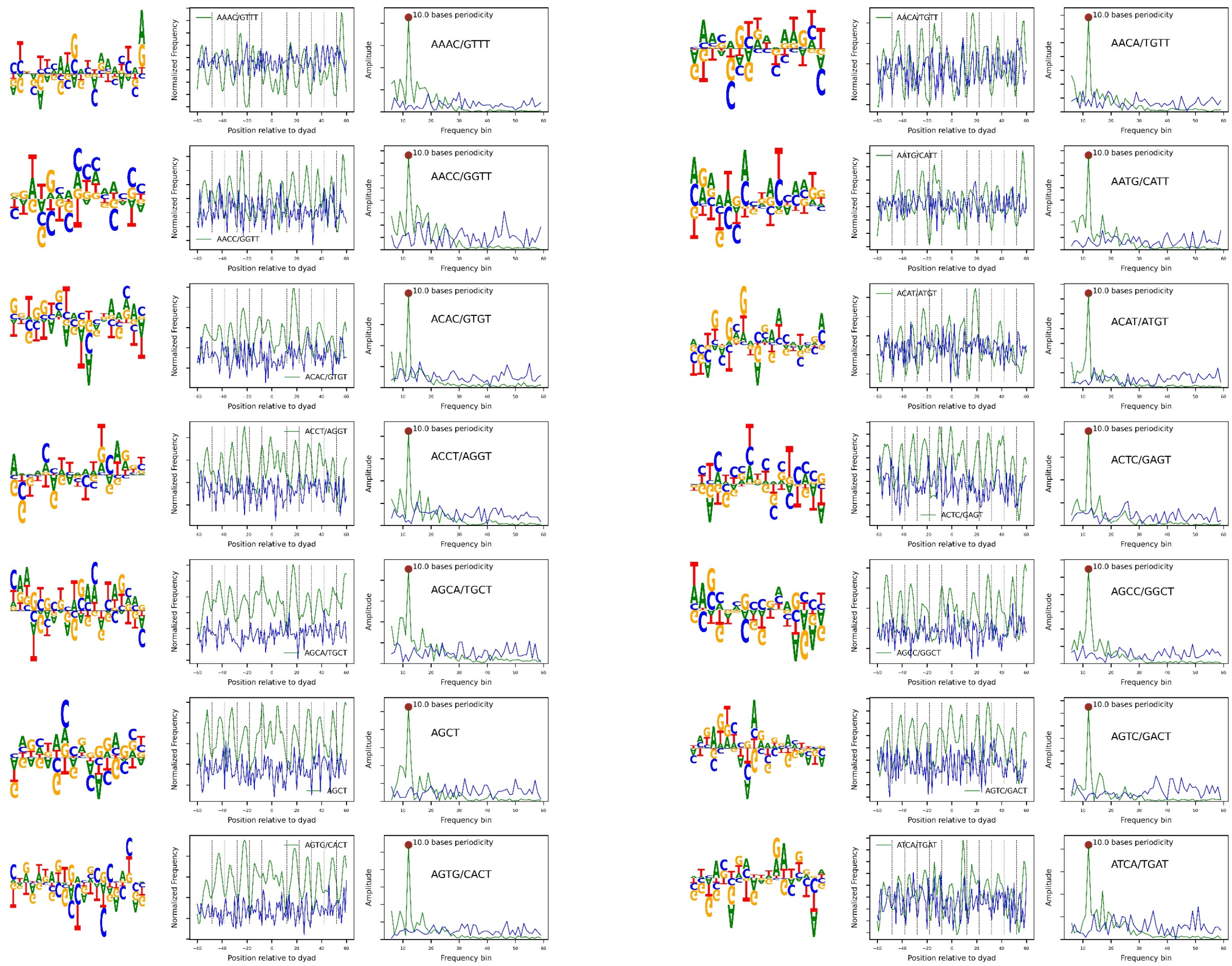


**Figure S10.** Kernels consisting of identical subregions spaced 10–11 bp apart. These subregions are referred to as *periodicity-related sequence motifs*. Long kernels (>15 bp) are visualized using sequence logos, and for each periodicity-related feature, both the frequency in nucleosomal (green) and inter-nucleosomal (blue) DNA sites and the results of fast Fourier transform (FFT) analysis of the frequency are shown. This figure presents some periodicity-related sequence features of length 4 bp.

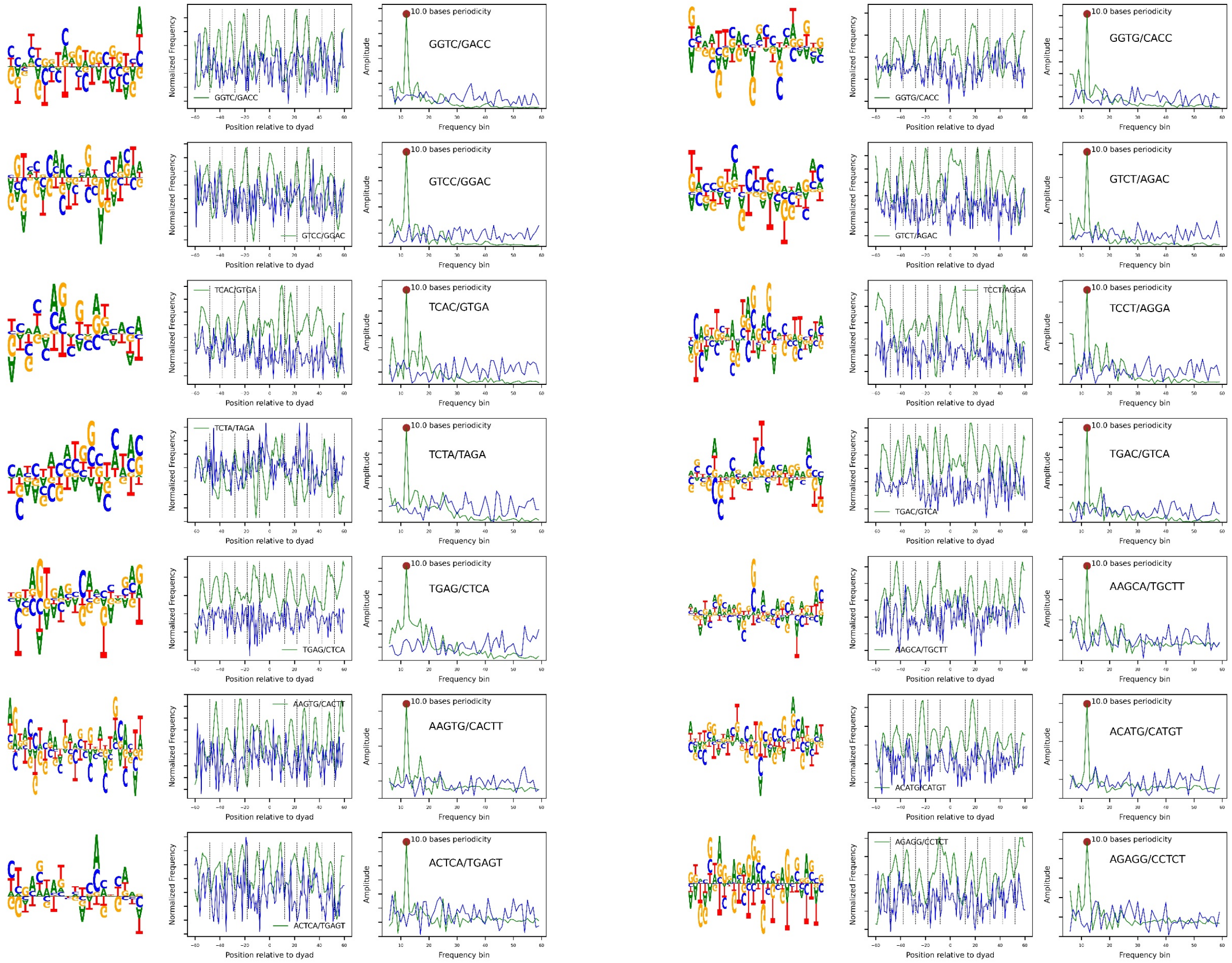


**Figure S11.** Kernels consisting of identical subregions spaced 10–11 bp apart. These subregions are referred to as *periodicity-related sequence motifs*. Long kernels (>15 bp) are visualized using sequence logos, and for each periodicity-related feature, both the frequency in nucleosomal (green) and inter-nucleosomal (blue) DNA sites and the results of fast Fourier transform (FFT) analysis of the frequency are shown. This figure presents some periodicity-related sequence features of length 4-5 bp.

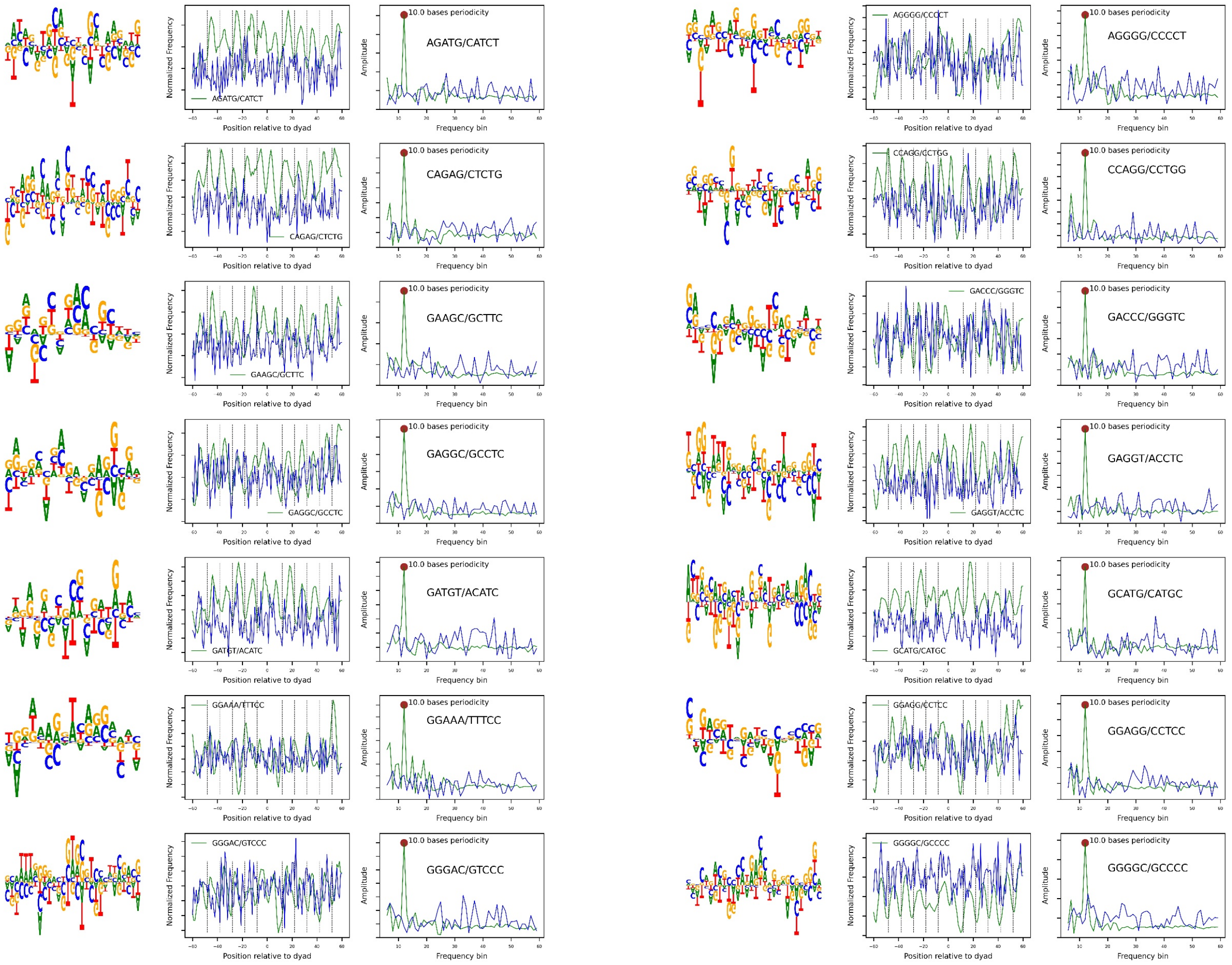


**Figure S12.** Kernels consisting of identical subregions spaced 10–11 bp apart. These subregions are referred to as *periodicity-related sequence features*. Long kernels (>15 bp) are visualized using sequence logos, and for each periodicity-related feature, both the frequency in nucleosomal (green) and inter-nucleosomal (blue) DNA sites and the results of fast Fourier transform (FFT) analysis of the frequency are shown. This figure presents some periodicity-related sequence features of length 5 bp.

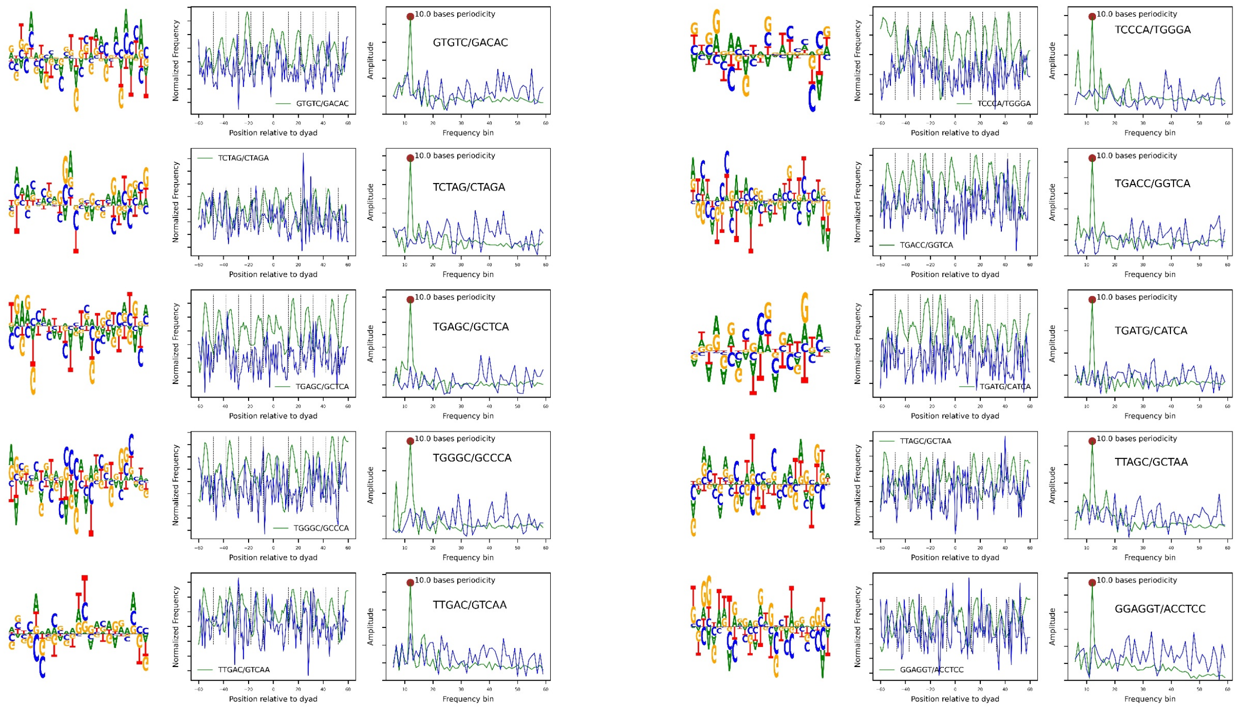


**Figure S13.** Kernels consisting of identical subregions spaced 10–11 bp apart. These subregions are referred to as *periodicity-related sequence motifs*. Long kernels (>15 bp) are visualized using sequence logos, and for each periodicity-related feature, both the frequency in nucleosomal (green) and inter-nucleosomal (blue) DNA sites and the results of fast Fourier transform (FFT) analysis of the frequency are shown. This figure presents some periodicity-related sequence features of length 5-6 bp.

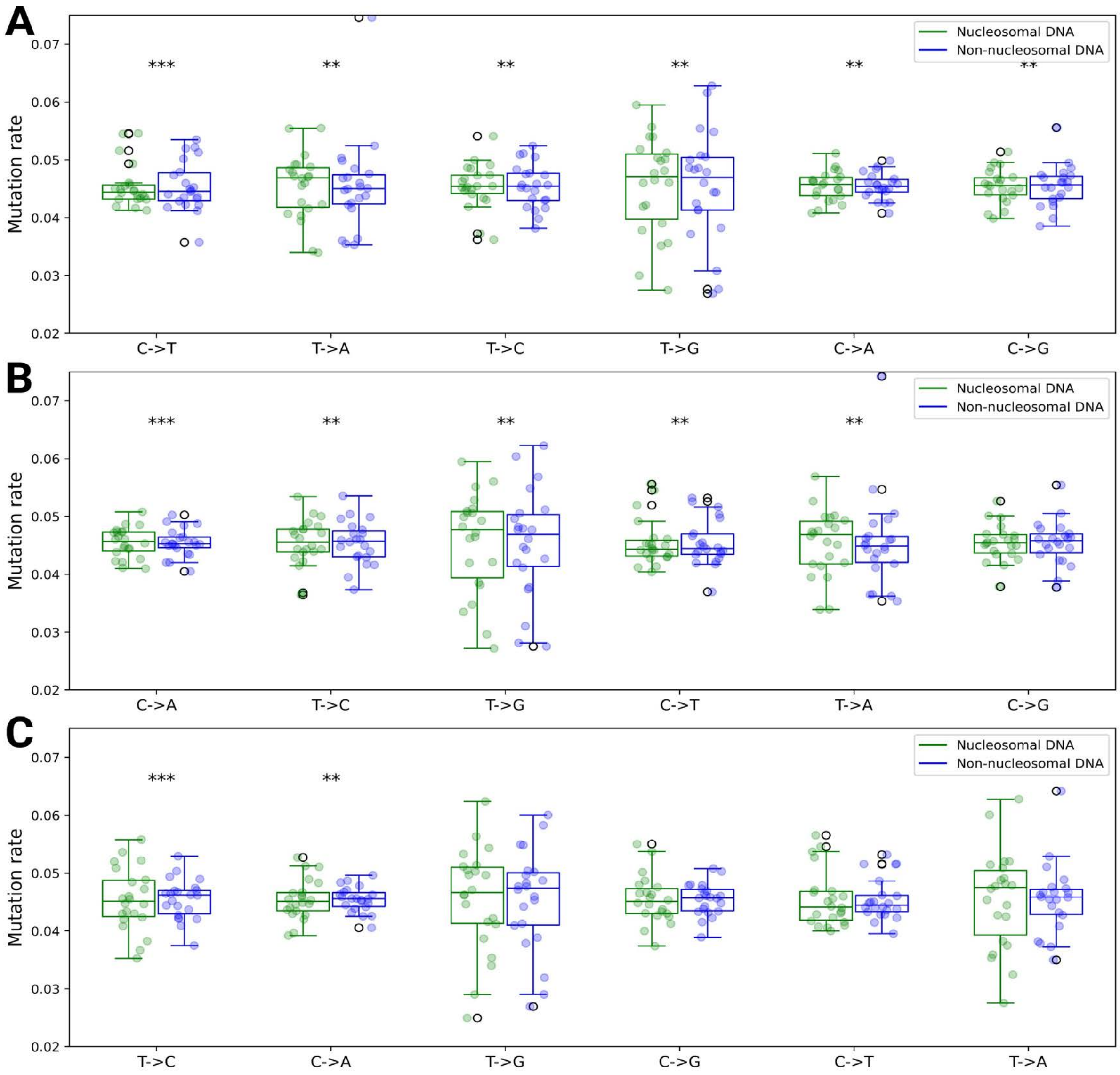


**Figure S14.** Distribution of single-nucleotide mutations in nucleosomal (green) and inter-nucleosomal regions. Symbols *** and ** indicate significant associations with p-values <0.01 and <0.05, respectively. The mutation rate is defined as the number of a specific single-nucleotide mutation divided by the total number of mutations. Each point within the boxplots represents an individual chromosome. (A) Distribution of single-nucleotide mutations in all nucleosomes (both well-positioned and non–well-positioned) using mutations that passed quality control. (B) Distribution of single-nucleotide mutations in all nucleosomes (both well-positioned and non–well-positioned) using Mutect2 quality-passed single-nucleotide mutations. (C) Distribution of single-nucleotide mutations in well-positioned nucleosomes using Mutect2 quality-passed single-nucleotide mutations.

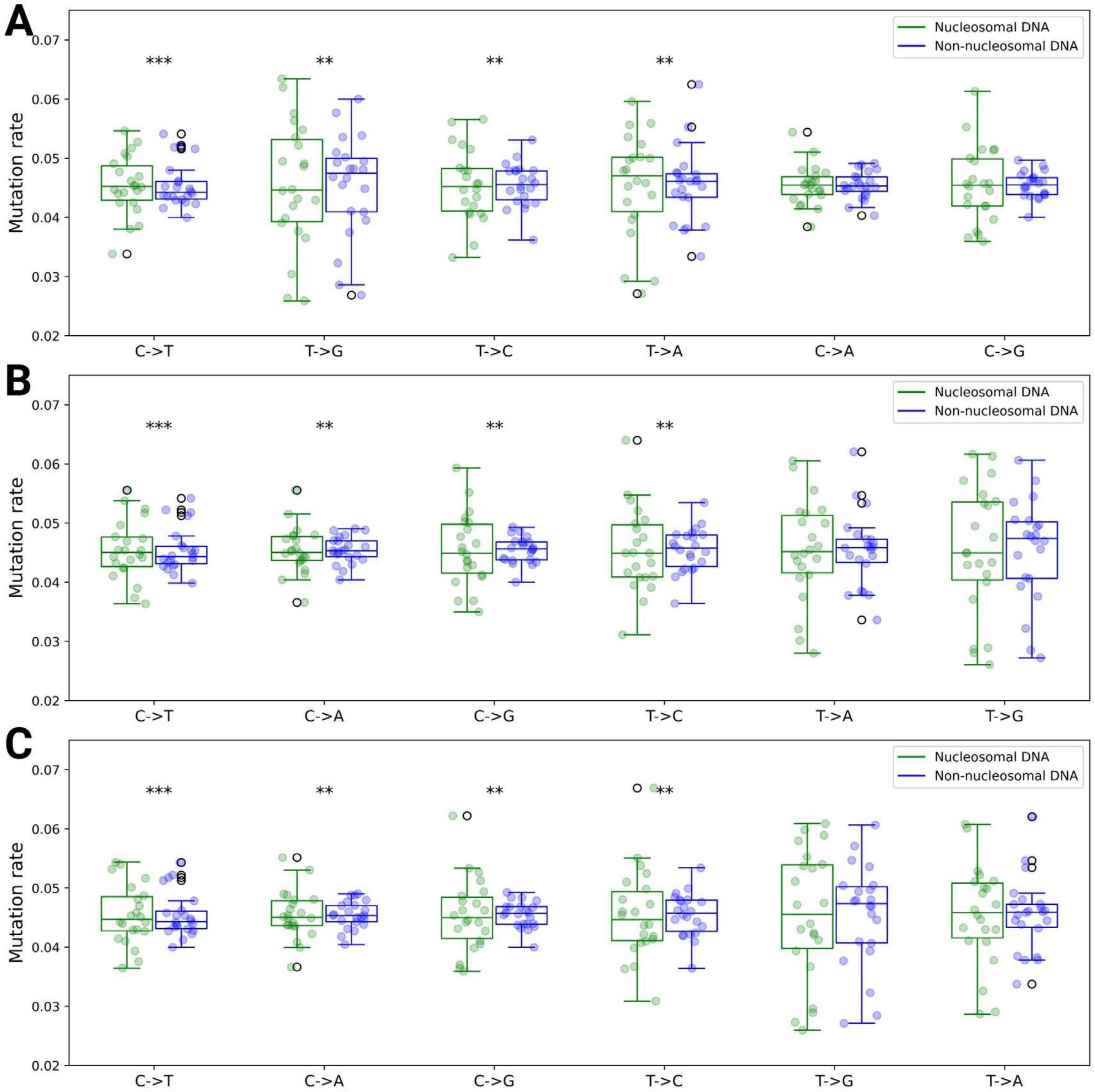


**Figure S15.** Distribution of single-nucleotide mutations in flanking linker DNA regions of nucleosomal (green) and inter-nucleosomal sites. Symbols *** and ** indicate significant associations with p-values <0.01 and <0.05, respectively. The mutation rate is defined as the number of a specific single-nucleotide mutation divided by the total number of mutations. Each point within the boxplots represents an individual chromosome. (A) Distribution of single-nucleotide mutations in flanking linker DNA regions of all nucleosomes (both well-positioned and non–well-positioned) using mutations that passed quality control. (B) Distribution of single-nucleotide mutations in flanking linker DNA regions of all nucleosomes (both well-positioned and non–well-positioned) using Mutect2 quality-passed single-nucleotide mutations. (C) Distribution of single-nucleotide mutations in flanking linker DNA regions of well-positioned nucleosomes using Mutect2 quality-passed single-nucleotide mutations.

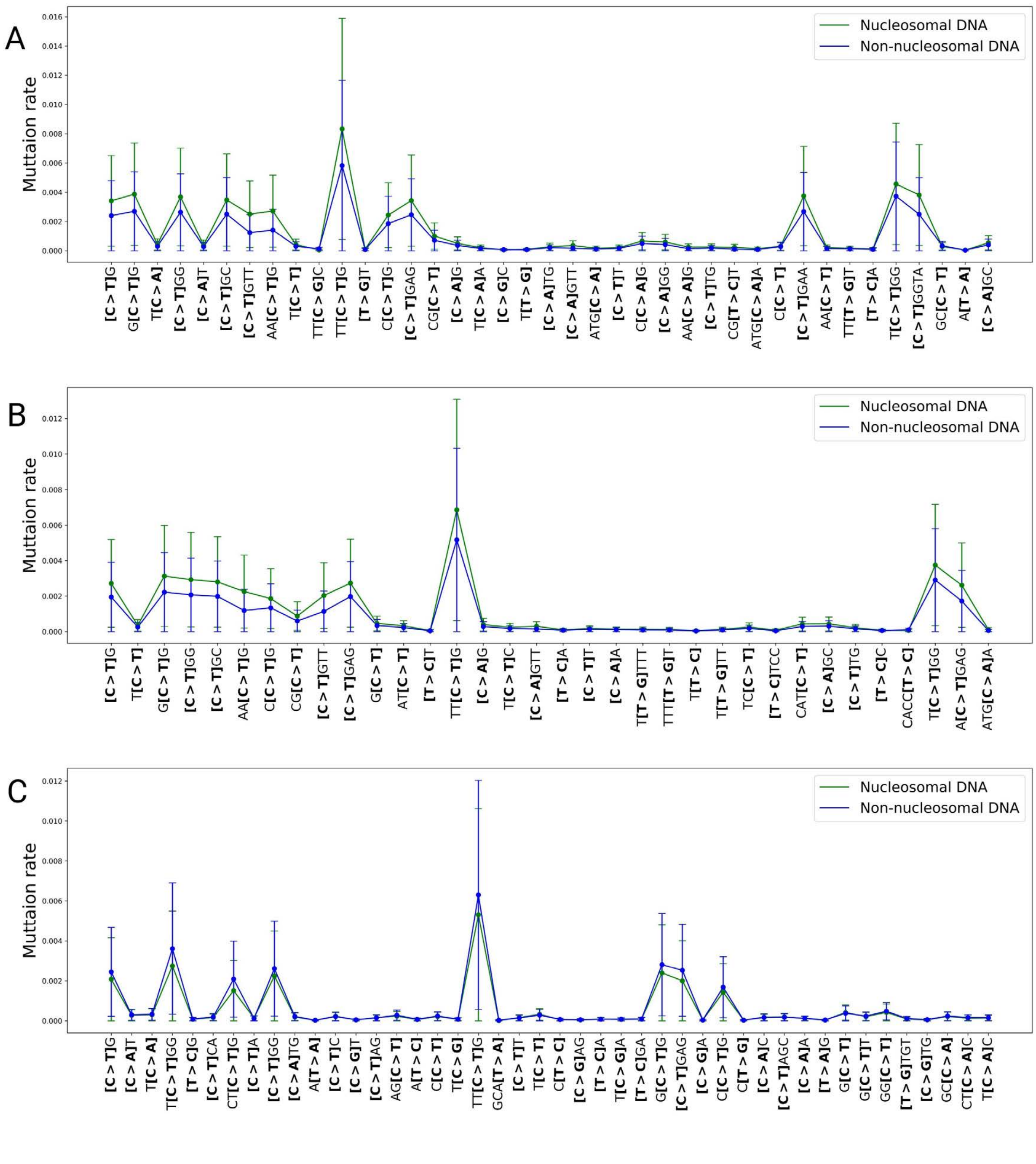


**Figure S16.** Mutational motifs significantly enriched or depleted in nucleosomes under different analysis scenarios. (A) Both well-positioned and non–well-positioned nucleosomes using mutations that passed quality control. (B) Both well-positioned and non–well-positioned nucleosomes using Mutect2 quality-passed single-nucleotide mutations. (C) Well-positioned nucleosomes using Mutect2 quality-passed single-nucleotide mutations.

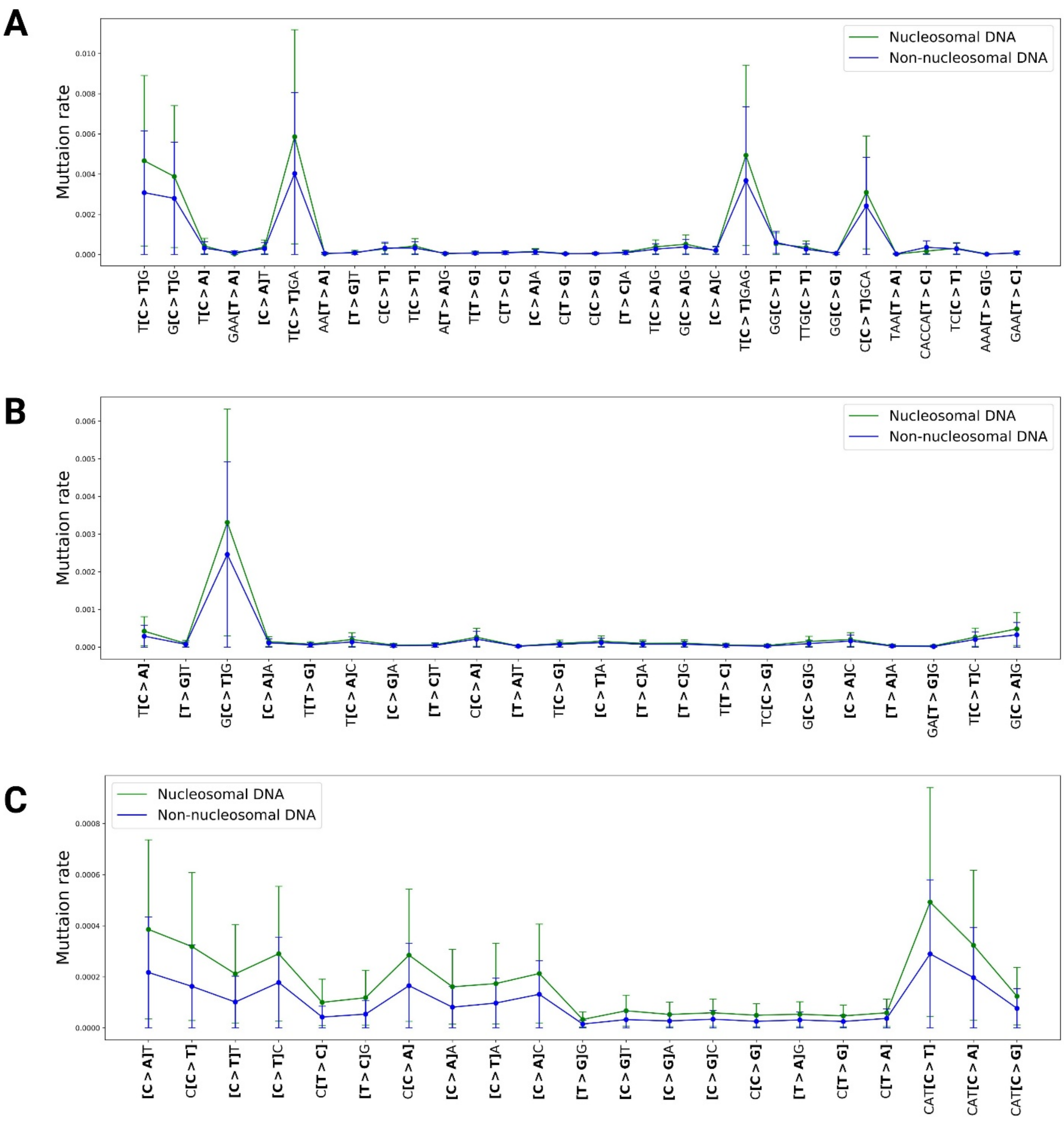


**Figure S17.** Mutational motifs significantly enriched or depleted in flanking regions of nucleosomes under different analysis scenarios. (A) Flanking linker DNA of both well-positioned and non–well-positioned nucleosomes using mutations that passed quality control. (B) Flanking linker DNA of both well-positioned and non–well-positioned nucleosomes using Mutect2 quality-passed single-nucleotide mutations. (C) Flanking linker DNA of well-positioned nucleosomes using Mutect2 quality-passed single-nucleotide mutations.

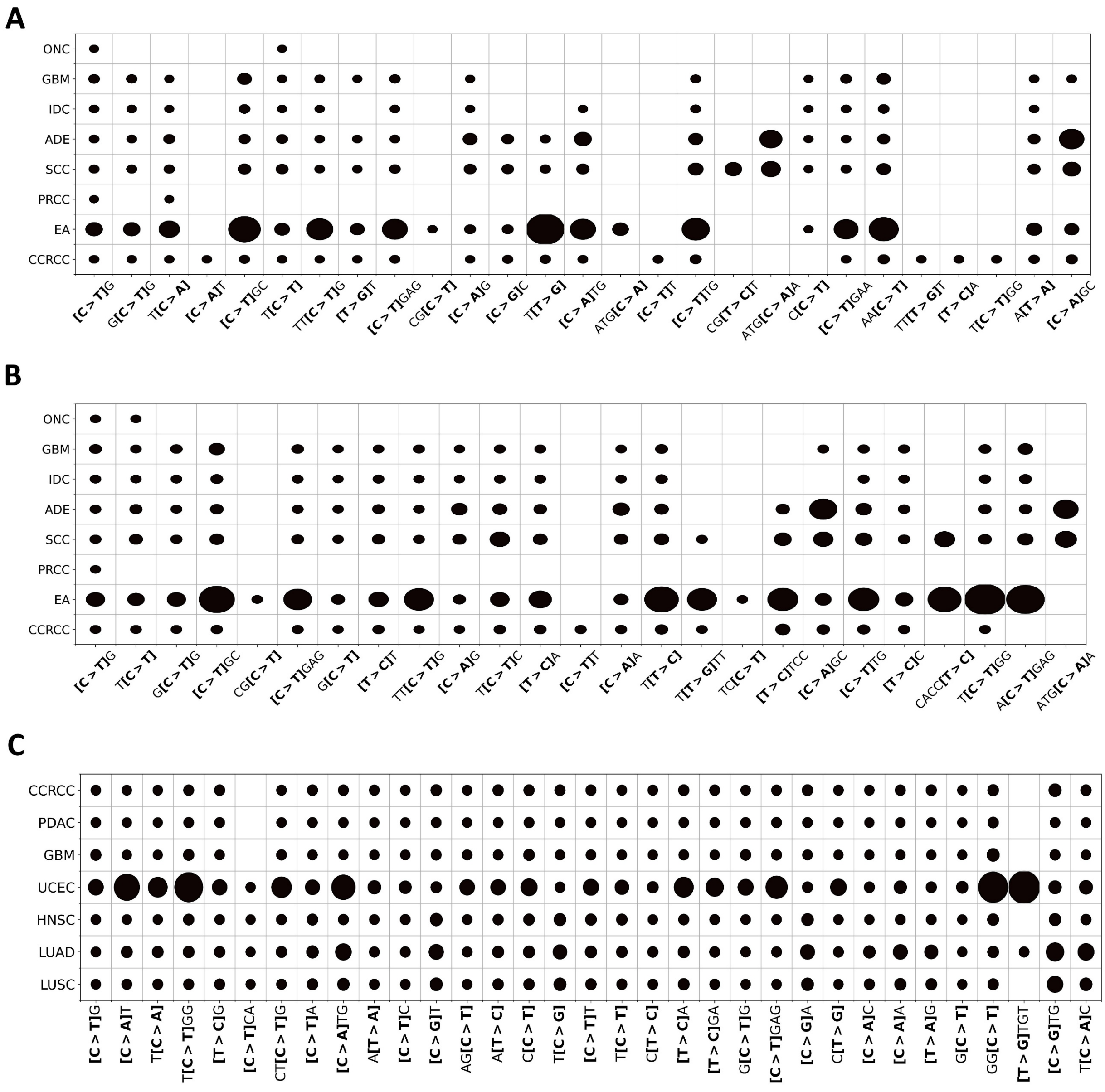


**Figure S18.** The association between mutational motifs and nucleosomes under different analysis scenarios. (A) Both well-positioned and non–well-positioned nucleosomes using mutations that passed quality control. (B) Both well-positioned and non–well-positioned nucleosomes using Mutect2 quality-passed single-nucleotide mutations. (C) Well-positioned nucleosomes using Mutect2 quality-passed single-nucleotide mutations. CCRCC: Renal cell carcinoma; HNSC: head and neck squamous cell carcinoma; UCEC: Urothelial carcinoma; EA: Endometrioid adenocarcinoma; PRCC: Papillary renal cell carcinoma; SCC: Squamous cell carcinoma; ADE: Adenocarcinoma; IDC: Infiltrating duct carcinoma; GBM: GBMoblastoma; ONC: Oncocytoma.

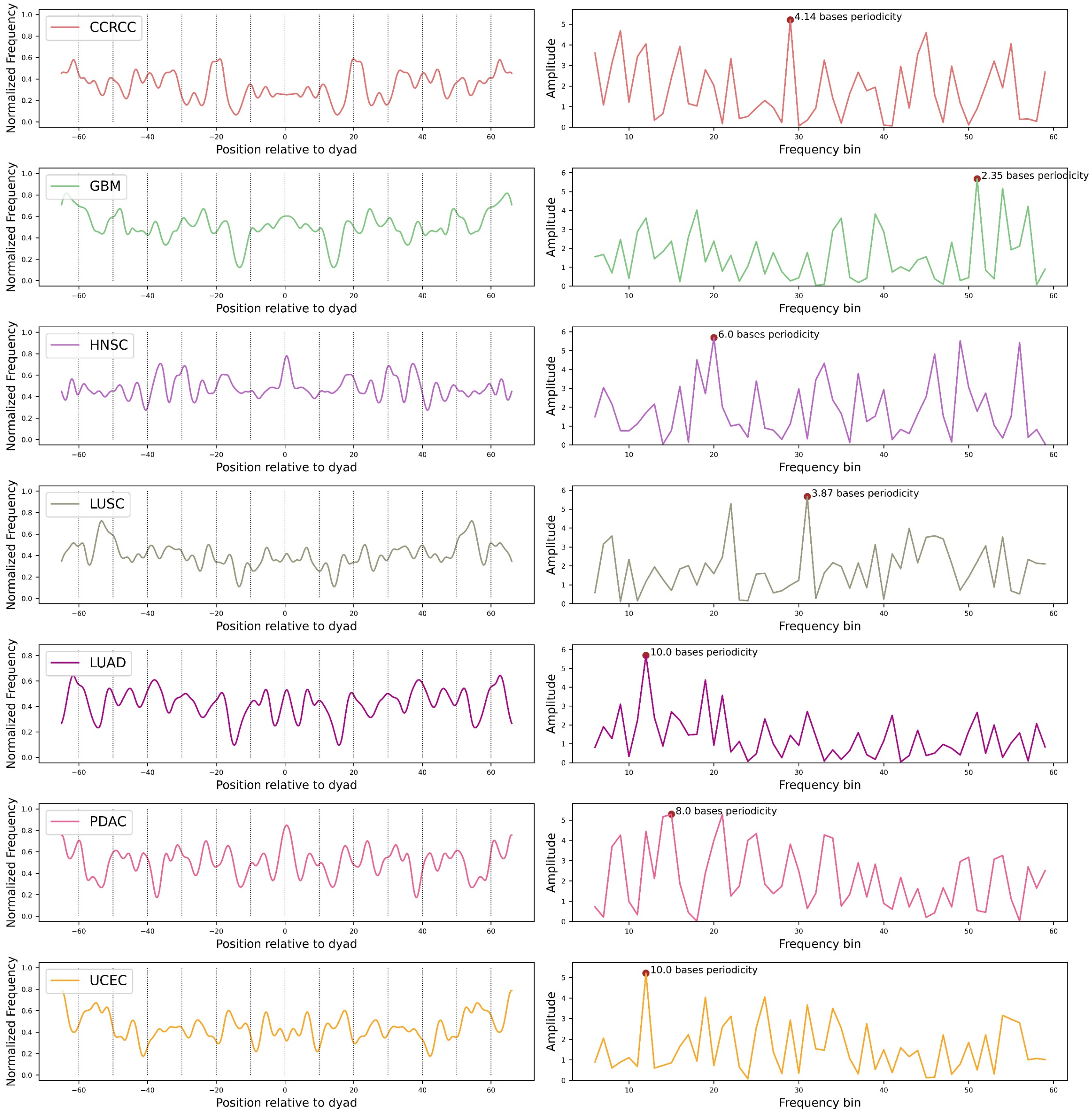


**Figure S19.** Distribution of mutations across nucleosomal footprint DNA relative to dyad positions across cancer types, including clear cell renal cell carcinoma (CCRCC), glioblastoma (GBM), head and neck squamous cell carcinoma (HNSC), lung squamous cell carcinoma (LUSC), lung adenocarcinoma (LUAD), pancreatic ductal adenocarcinoma (PDAC), and uterine corpus endometrial carcinoma (UCEC).

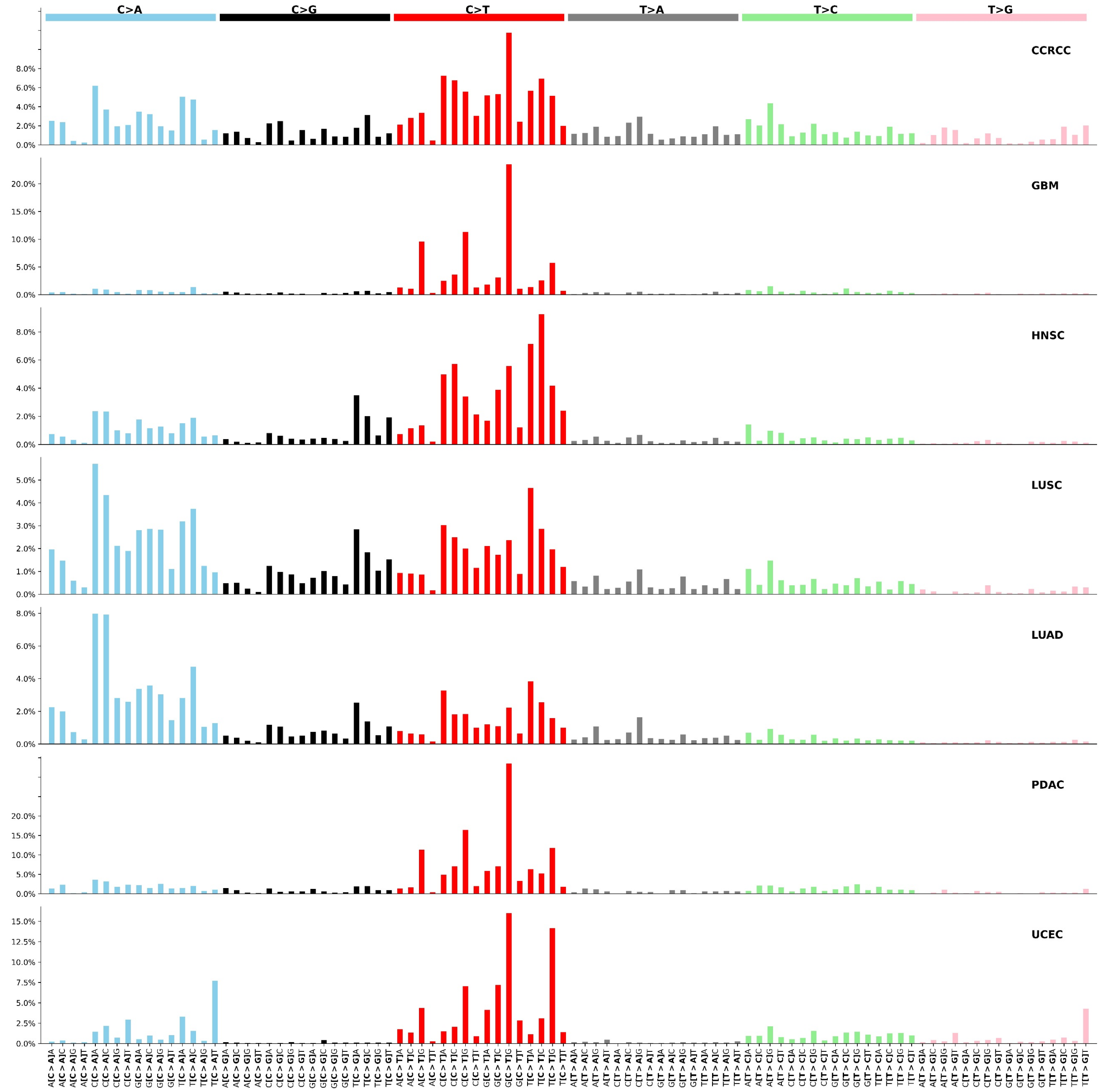


**Figure S20.** Cancer-type–specific mutational profiles derived from statistically significant mutational motifs, including clear cell renal cell carcinoma (CCRCC), glioblastoma (GBM), head and neck squamous cell carcinoma (HNSC), lung squamous cell carcinoma (LUSC), lung adenocarcinoma (LUAD), pancreatic ductal adenocarcinoma (PDAC), and uterine corpus endometrial carcinoma (UCEC).

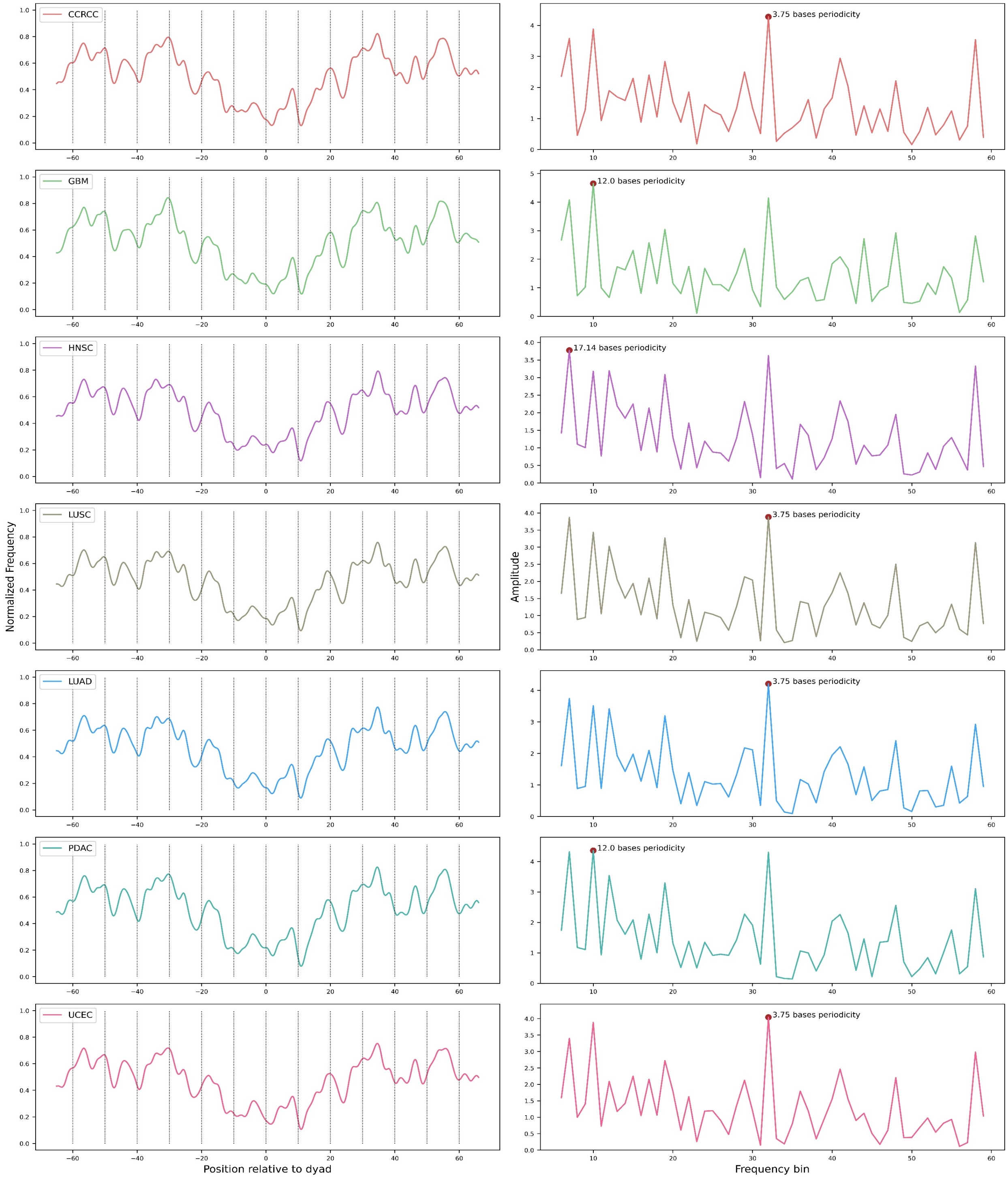


**Figure S21.** Distribution of CpG methylation across nucleosomal footprint DNA relative to dyad positions in different cancer types, including clear cell renal cell carcinoma (CCRCC), glioblastoma (GBM), head and neck squamous cell carcinoma (HNSC), lung squamous cell carcinoma (LUSC), lung adenocarcinoma (LUAD), pancreatic ductal adenocarcinoma (PDAC), and uterine corpus endometrial carcinoma (UCEC).

**Table S1.** Performance improvements from incorporating short flanking linker DNA across machine learning methods and data representations. Bold values in each column indicate the method with the highest performance for the corresponding metric.

| Data representation | Presence of linker DNA | Method | Accuracy | Specificity | Sensitivity | Precision | F1 score | MCC | C Kappa |
|---|---|---|---|---|---|---|---|---|---|
| One-hot | No linker | SVM* | 0.62 | 0.61 | 0.63 | 0.62 | 0.63 | 0.25 | 0.28 |
| | | CNN* | 0.63 | 0.51 | 0.77 | 0.61 | 0.68 | 0.29 | 0.30 |
| | | ResNet* | 0.65 | 0.61 | 0.69 | 0.64 | 0.66 | 0.30 | 0.33 |
| | | DeepND* | 0.81 | 0.77 | 0.85 | 0.79 | 0.82 | 0.63 | 0.67 |
| | With linker | SVM | 0.75 | 0.74 | 0.76 | 0.75 | 0.75 | 0.50 | 0.55 |
| | | CNN | 0.78 | 0.78 | 0.78 | 0.78 | 0.78 | 0.56 | 0.61 |
| | | ResNet | 0.80 | 0.85 | 0.76 | 0.84 | 0.79 | 0.61 | 0.67 |
| | | DeepND | **0.93** | 0.91 | **0.94** | 0.92 | **0.93** | **0.86** | **0.88** |
| Sequence features | No linker | SVM* | 0.59 | 0.51 | 0.67 | 0.58 | 0.62 | 0.18 | 0.20 |
| | | CNN* | 0.66 | 0.59 | 0.74 | 0.64 | 0.69 | 0.33 | 0.36 |
| | | ResNet* | 0.68 | 0.69 | 0.67 | 0.68 | 0.68 | 0.36 | 0.40 |
| | | $SVM^{Trader*}$ | 0.74 | 0.69 | 0.80 | 0.72 | 0.76 | 0.49 | 0.53 |
| | | $NuPoSe^{WS*}$ | 0.79 | 0.76 | 0.81 | 0.77 | 0.79 | 0.57 | 0.62 |
| | | NuPoSe* | 0.80 | 0.84 | 0.76 | 0.82 | 0.79 | 0.60 | 0.66 |
| | With linker | SVM | 0.60 | 0.56 | 0.63 | 0.59 | 0.61 | 0.20 | 0.22 |
| | | CNN | 0.70 | 0.68 | 0.72 | 0.69 | 0.71 | 0.40 | 0.45 |
| | | ResNet | 0.71 | 0.60 | 0.83 | 0.67 | 0.74 | 0.44 | 0.46 |
| | | $SVM^{Trader}$ | 0.85 | **0.97** | 0.74 | 0.96 | 0.84 | 0.73 | 0.78 |
| | | $NuPoSe^{WS}$ | 0.88 | **0.97** | 0.79 | **0.97** | 0.87 | 0.78 | 0.84 |
| | | NuPoSe | 0.89 | **0.97** | 0.80 | **0.97** | 0.88 | 0.79 | 0.84 |

**Table S2.** Comparison of different machine learning methods on gold-standard datasets. The size of positive and negative samples for each dataset is indicated in parentheses. Bold values in each column indicate the method with the highest performance for the corresponding metric.

| Train/test dataset | K-fold | Method | Sensitivity | Specificity | Accuracy | MCC | AUC |
|---|---|---|---|---|---|---|---|
| *H.sapiens (2273, 2300)* | K= 5 | DeepND | **0.90** | **0.89** | **0.89** | **0.82** | 0.95 |
| | | NuPoSe* | 0.89 | **0.89** | **0.89** | 0.78 | **0.96** |
| | | LeNup | **0.90** | 0.84 | 0.87 | 0.77 | --- |
| | K= 20 | DeepND | 0.92 | 0.91 | 0.92 | 0.82 | 0.97 |
| | | NuPoSe* | **0.94** | **0.94** | **0.94** | **0.85** | **0.98** |
| | | LeNup | 0.92 | 0.85 | 0.88 | 0.79 | 0.94 |
| | K= M | DeepND | 0.96 | **0.95** | 0.94 | 0.84 | 0.98 |
| | | NuPoSe* | **0.98** | **0.95** | **0.97** | **0.88** | **0.99** |
| | | 3LS | 0.91 | 0.88 | 0.90 | 0.80 | 0.95 |
| | | iNuc-PseKNC | 0.87 | 0.84 | 0.86 | 0.73 | 0.90 |
| | | iNuc-STNC | 0.89 | 0.85 | 0.87 | 0.75 | --- |
| *C.elegans (2567, 2608)* | K= 5 | DeepND | 0.92 | **0.92** | **0.92** | **0.87** | **0.96** |
| | | NuPoSe* | **0.95** | 0.84 | 0.89 | 0.80 | 0.96 |
| | K= 20 | DeepND | 0.93 | 0.90 | **0.92** | 0.84 | 0.96 |
| | | NuPoSe* | **0.95** | **0.91** | **0.92** | **0.86** | **0.98** |
| | | LeNup | 0.93 | 0.90 | 0.91 | 0.84 | 0.96 |
| | K= M | DeepND | 0.96 | 0.97 | 0.97 | 0.87 | 0.98 |
| | | NuPoSe* | **0.98** | **0.98** | **0.98** | **0.89** | **0.99** |
| | | 3LS | 0.86 | 0.89 | 0.87 | 0.75 | 0.95 |
| | | iNuc-PseKNC | 0.90 | 0.83 | 0.86 | 0.74 | 0.93 |
| | | iNuc-STNC | 0.91 | 0.86 | 0.88 | 0.77 | --- |
| *D.melanogaster (2900, 2850)* | K= 5 | DeepND | **0.88** | 0.87 | **0.88** | **0.80** | **0.93** |
| | | NuPoSe* | 0.79 | **0.88** | 0.83 | 0.65 | 0.90 |
| | K= 20 | DeepND | **0.90** | 0.89 | **0.90** | **0.82** | **0.95** |
| | | NuPoSe* | 0.86 | **0.91** | 0.87 | 0.76 | 0.93 |
| | | LeNup | 0.89 | 0.87 | 0.88 | 0.78 | 0.94 |
| | K= M | DeepND | **0.92** | **0.93** | **0.91** | **0.84** | **0.96** |
| | | NuPoSe* | 0.90 | **0.93** | **0.91** | 0.80 | 0.95 |
| | | 3LS | 0.84 | 0.82 | 0.83 | 0.66 | 0.91 |
| | | iNuc-PseKNC | 0.78 | 0.81 | 0.79 | 0.60 | 0.87 |
| | | iNuc-STNC | 0.79 | 0.83 | 0.81 | 0.63 | --- |

**Table S3.** (a) Identified sequence motifs significantly enriched or depleted in nucleosomal DNA

| Row No. | Sequence Motif | P-value | Adjusted P-value | Odds Ratio |
|---|---|---|---|---|
| 1 | AGTACAGGCACAG/CTGTGCCTGTACT | 0.000172831 | 0.035664336 | 1.891414135 |
| 2 | TGCGCACTCACAAGC/GCTTGTGAGTGCGCA | 0.006268046 | 0.048601399 | 1.890331122 |
| 3 | TCCCATAGGCTTGGA/TCCAAGCCTATGGGA | 0.001798448 | 0.041258741 | 1.782357742 |
| 4 | GCTTCATTTAGCT/AGCTAAATGAAGC | 0.000657703 | 0.038461538 | 1.68298796 |
| 5 | CCTCATCATATCT/AGATATGATGAGG | 0.001503721 | 0.04020979 | 1.649896073 |
| 6 | GTTCCTGTGAAGT/ACTTCACAGGAAC | 0.001859077 | 0.041608392 | 1.640676634 |
| 7 | GACTAGGTTGTC/GACAACCTAGTC | 0.001565135 | 0.040559441 | 1.605850994 |
| 8 | TCAGGAGTGTGTAGG/CCTACACACTCCTGA | 0.005450403 | 0.047552448 | 1.528400288 |
| 9 | GACCAGAAGGAGACA/TGTCTCCTTCTGGTC | 0.002611425 | 0.043706294 | 1.517903829 |
| 10 | CCTCAGAACAAGG/CCTTGTTCTGAGG | 0.005341453 | 0.047202797 | 1.510358171 |
| 11 | CAGGAGGTAAAGC/GCTTTACCTCCTG | 0.009106476 | 0.04965035 | 1.509444635 |
| 12 | TGCCATATGTGG/CCACATATGGCA | 0.000982789 | 0.039160839 | 1.375795308 |
| 13 | TCTTGTCACCCT/AGGGTGACAAGA | 0.003284924 | 0.045804196 | 1.278965902 |
| 14 | TGATGGCT/AGCCATCA | 6.80548E-15 | 0.02027972 | 1.25129228 |
| 15 | TCAGGCTTGTC/GACAAGCCTGA | 3.72312E-05 | 0.032517483 | 1.246653085 |
| 16 | GCAAAGGAG/CTCCTTTGC | 5.70626E-06 | 0.029370629 | 1.226525303 |
| 17 | AGACCTTGC/GCAAGGTCT | 0.001251075 | 0.03986014 | 1.223185934 |
| 18 | GTTTCGTAGCT/AGCTACGAAAC | 0.006517919 | 0.048951049 | 1.21759655 |
| 19 | CTGCTTGT/ACAAGCAG | 1.76095E-13 | 0.022377622 | 1.215004483 |
| 20 | CTCCTTCTCGT/ACGAGAAGGAG | 0.000163207 | 0.035314685 | 1.198638802 |
| 21 | GACAAGCTGA/TCAGCTTGTC | 6.70746E-11 | 0.025524476 | 1.196851934 |
| 22 | CTGGCTGA/TCAGCCAG | 1.79147E-10 | 0.026223776 | 1.183327107 |
| 23 | AAGGAAGG/CCTTCCTT | 4.42062E-14 | 0.021678322 | 1.179444555 |
| 24 | TGCCTTGGTA/TACCAAGGCA | 3.99488E-08 | 0.027622378 | 1.168524542 |
| 25 | AAGCAGGA/TCCTGCTT | 3.97045E-11 | 0.025174825 | 1.164453247 |
| 26 | GATGCCCAG/CTGGGCATC | 0.002715339 | 0.044405594 | 1.16107914 |
| 27 | GATG/CATC | 7.86294E-52 | 0.01013986 | 1.15824616 |
| 28 | CACA/TGTG | 2.66257E-69 | 0.007342657 | 1.155781624 |
| 29 | ATGC/GCAT | 3.97709E-47 | 0.011538462 | 1.15517757 |
| 30 | CTCCCCTGA/TCAGGGGAG | 0.004120279 | 0.046503497 | 1.138529826 |
| 31 | CCTAGGAT/ATCCTAGG | 0.00014035 | 0.034965035 | 1.136861851 |
| 32 | CATCAG/CTGATG | 1.91755E-54 | 0.00979021 | 1.136036495 |
| 33 | ACTGCA/TGCAGT | 7.2139E-51 | 0.01048951 | 1.128308416 |
| 34 | GAGCTAAC/GTTAGCTC | 0.002166517 | 0.042657343 | 1.12773197 |
| 35 | TAGC/GCTA | 3.46566E-23 | 0.016783217 | 1.126277392 |
| 36 | ATGGTGAA/TTCACCAT | 3.07198E-06 | 0.028671329 | 1.123401765 |
| 37 | GCTAGAT/ATCTAGC | 1.61804E-09 | 0.026923077 | 1.122304209 |
| 38 | CCATG/CATGG | 1.5073E-125 | 0.004895105 | 1.121679978 |
| 39 | ATGGCAT/ATGCCAT | 4.04928E-14 | 0.021328671 | 1.119207924 |
| 40 | GCAAGAT/ATCTTGC | 1.41107E-12 | 0.023426573 | 1.118670653 |
| 41 | CTCAGC/GCTGAG | 2.44232E-43 | 0.012937063 | 1.11713785 |
| 42 | AATGGAGCTC/GAGCTCCATT | 0.000189952 | 0.036013986 | 1.117074724 |
| 43 | ACGATC/GATCGT | 2.25621E-14 | 0.020629371 | 1.117007565 |
| 44 | TGAGTGG/CCACTCA | 5.37423E-13 | 0.023076923 | 1.116585181 |
| 45 | CAG/CTG | 5.7521E-153 | 0.004195804 | 1.113230875 |
| 46 | TCCTGTATCC/GGATACAGGA | 8.36474E-05 | 0.033566434 | 1.109472828 |
| 47 | GTTTGGCCTT/AAGGCCAAAC | 0.000119888 | 0.034615385 | 1.109350785 |
| 48 | CAGC/GCTG | 5.50966E-35 | 0.013986014 | 1.109169622 |
| 49 | AGTAGCTC/GAGCTACT | 0.002419834 | 0.043006993 | 1.108996703 |
| 50 | CTGTG/CACAG | 4.3434E-104 | 0.005594406 | 1.103962102 |
| 51 | CTAG | 6.93262E-18 | 0.018881119 | 1.103068121 |
| 52 | CTGCGT/ACGCAG | 2.1574E-21 | 0.017482517 | 1.102299636 |
| 53 | CAACGAT/ATCGTTG | 7.63063E-05 | 0.033216783 | 1.099447558 |
| 54 | CCTTTGTG/CACAAAGG | 0.000216893 | 0.036363636 | 1.096650431 |
| 55 | ACGACT/AGTCGT | 4.92085E-13 | 0.022727273 | 1.096112338 |
| 56 | CCAC/GTGG | 4.25537E-24 | 0.016433566 | 1.096098929 |
| 57 | TG/CA | 0 | 0.00034965 | 1.092679509 |
| 58 | AAGCTG/CAGCTT | 3.31514E-30 | 0.014335664 | 1.092348463 |
| 59 | CCA/TGG | 7.64173E-88 | 0.006293706 | 1.090498882 |
| 60 | CACCT/AGGTG | 1.33923E-70 | 0.006993007 | 1.088795604 |
| 61 | ACCTGTATAG/CTATACAGGT | 0.007092091 | 0.049300699 | 1.087997848 |
| 62 | AGACCA/TGGTCT | 2.74725E-22 | 0.017132867 | 1.086082146 |
| 63 | AGC/GCT | 1.03992E-64 | 0.008391608 | 1.085293659 |

| | | | | |
|---|---|---|---|---|
| 64 | TACC/GGTA | 2.02838E-11 | 0.024475524 | 1.082687103 |
| 65 | CATTG/CAATG | 6.74996E-69 | 0.007692308 | 1.08170003 |
| 66 | CGAA/TTCG | 0.000755626 | 0.038811189 | 1.076629255 |
| 67 | TCCAG/CTGGA | 2.45403E-59 | 0.008741259 | 1.076028568 |
| 68 | CACGAG/CTCGTG | 1.17696E-10 | 0.025874126 | 1.075365375 |
| 69 | ATGCA/TGCAT | 7.13358E-57 | 0.009090909 | 1.073866961 |
| 70 | TCCAAC/GTTGGA | 1.5843E-15 | 0.01958042 | 1.072678917 |
| 71 | GGAGGAA/TTCCTCC | 2.47102E-08 | 0.027272727 | 1.072598536 |
| 72 | CCTCT/AGAGG | 1.25487E-48 | 0.011188811 | 1.070236351 |
| 73 | AATTCCACCT/AGGTGGAATT | 0.005610548 | 0.047902098 | 1.068471864 |
| 74 | TGTCA/TGACA | 1.66521E-50 | 0.010839161 | 1.067827457 |
| 75 | TCATCTCT/AGAGATGA | 0.002670149 | 0.044055944 | 1.066481677 |
| 76 | CTCGT/ACGAG | 5.80354E-25 | 0.016083916 | 1.065616861 |
| 77 | AAGC/GCTT | 5.23607E-12 | 0.024125874 | 1.065062282 |
| 78 | AG/CT | 4.1156E-155 | 0.003496503 | 1.062512647 |
| 79 | TCTGG/CCAGA | 6.03597E-41 | 0.013286713 | 1.062038645 |
| 80 | TGA/TCA | 9.1792E-44 | 0.012587413 | 1.059815577 |
| 81 | TGTGAT/ATCACA | 3.94579E-14 | 0.020979021 | 1.059434021 |
| 82 | CGTT/AACG | 0.009664805 | 0.05 | 1.054757885 |
| 83 | ATGGT/ACCAT | 2.9632E-28 | 0.015034965 | 1.054596473 |
| 84 | ATGTCA/TGACAT | 3.3877E-12 | 0.023776224 | 1.053232127 |
| 85 | TCC/GGA | 8.98543E-29 | 0.014685315 | 1.051827364 |
| 86 | AGT/ACT | 5.03252E-28 | 0.015384615 | 1.051759849 |
| 87 | GGAA/TTCC | 4.08292E-10 | 0.026573427 | 1.049635952 |
| 88 | TC/GA | 6.0718E-76 | 0.006643357 | 1.045903664 |
| 89 | GAT/ATC | 1.8461E-18 | 0.018531469 | 1.045671387 |
| 90 | GACC/GGTC | 0.000253425 | 0.037412587 | 1.043478395 |
| 91 | GC | 6.03747E-47 | 0.011888112 | 1.039880615 |
| 92 | AATGC/GCATT | 1.36445E-16 | 0.019230769 | 1.038446625 |
| 93 | GACAA/TTGTC | 1.5363E-13 | 0.022027972 | 1.033932685 |
| 94 | TTCCT/AGGAA | 2.16694E-11 | 0.024825175 | 1.028485285 |
| 95 | CGGTA/TACCG | 0.000239678 | 0.036713287 | 1.026892128 |
| 96 | CTCGC/GCGAG | 2.99058E-05 | 0.031468531 | 1.026885849 |
| 97 | CC/GG | 8.57747E-20 | 0.017832168 | 1.025522157 |
| 98 | GGAG/CTCC | 0.002425149 | 0.043356643 | 1.025093466 |
| 99 | ACAA/TTGT | 0.001913318 | 0.042307692 | 1.024986209 |
| 100 | CCCA/TGGG | 0.003244636 | 0.045454545 | 1.023680279 |
| 101 | AAC/GTT | 2.25728E-06 | 0.028321678 | 1.022803775 |
| 102 | GGC/GCC | 8.52633E-05 | 0.033916084 | 0.980731845 |
| 103 | GTCCG/CGGAC | 2.43498E-05 | 0.030769231 | 0.971680937 |
| 104 | CTAT/ATAG | 0.000611246 | 0.038111888 | 0.967630236 |
| 105 | CTCG/CGAG | 0.003414988 | 0.046153846 | 0.950816512 |
| 106 | ATAGAGT/ACTCTAT | 0.000372519 | 0.037762238 | 0.948741842 |
| 107 | AAATAGC/GCTATTT | 2.73249E-05 | 0.031118881 | 0.947937722 |
| 108 | AT | 3.2684E-195 | 0.002797203 | 0.937567424 |
| 109 | AACGAAT/ATTCGTT | 7.25143E-05 | 0.032867133 | 0.934413091 |
| 110 | TT/AA | 2.4687E-210 | 0.002447552 | 0.933489571 |
| 111 | CGGGCT/AGCCCG | 4.79985E-15 | 0.01993007 | 0.922096676 |
| 112 | CCGA/TCGG | 4.71472E-06 | 0.029020979 | 0.918494754 |
| 113 | TA | 0 | 0.001048951 | 0.90672284 |
| 114 | CCGGATC/GATCCGG | 3.63086E-05 | 0.032167832 | 0.903033254 |
| 115 | GTGCGGC/GCCGCAC | 3.78747E-07 | 0.027972028 | 0.900627096 |
| 116 | CG | 1.3412E-155 | 0.003146853 | 0.889550057 |
| 117 | TCAAGGAAA/TTTCCTTGA | 0.000252541 | 0.037062937 | 0.886090306 |
| 118 | AAAAA/TTTTT | 3.0445E-253 | 0.001748252 | 0.869275853 |
| 119 | AAAAGT/ACTTTT | 2.6377E-137 | 0.004545455 | 0.86498168 |
| 120 | GAAGGCGGGCG/CGCCCGCCTTC | 0.001881005 | 0.041958042 | 0.84638395 |
| 121 | CGGGGCT/AGCCCCG | 2.39534E-25 | 0.015734266 | 0.841367424 |
| 122 | CGGTCTGGCG/CGCCAGACCG | 1.51106E-05 | 0.02972028 | 0.834315938 |
| 123 | GCG/CGC | 4.8322E-108 | 0.005244755 | 0.831000352 |
| 124 | AAAA/TTTT | 3.9558E-259 | 0.001398601 | 0.821438426 |
| 125 | AAAAAA/TTTTTT | 2.7634E-243 | 0.002097902 | 0.821053312 |
| 126 | GAGCTGCGGGGA/TCCCCGCAGCTC | 0.004548847 | 0.046853147 | 0.819676196 |
| 127 | CGG/CCG | 1.7822E-153 | 0.003846154 | 0.818365687 |
| 128 | GAAAAGGATTTC/GAAATCCTTTTC | 9.43138E-05 | 0.034265734 | 0.798859148 |
| 129 | GCCGCG/CGCGGC | 6.24065E-88 | 0.005944056 | 0.789021361 |

| | | | | |
|---|---|---|---|---|
| 130 | TCCCCGGGTCTC/GAGACCCGGGGA | 0.00296153 | 0.044755245 | 0.785954811 |
| 131 | TAAAGATT/AATCTTTA | 8.02627E-57 | 0.009440559 | 0.75667606 |
| 132 | TTTTCAAATGCATTC/GAATGCATTTGAAAA | 1.90196E-05 | 0.03006993 | 0.751115309 |
| 133 | AGGGACGGGGGGCCT/AGGCCCCCCGTCCCT | 0.001115799 | 0.03951049 | 0.661438478 |
| 134 | CCCCGCTGGCGGC/GCCGCCAGCGGGG | 0.001732002 | 0.040909091 | 0.654108981 |
| 135 | GGGCCCGGCACTCGGA/TCCGAGTGCCGGGCCC | 0.005984159 | 0.048251748 | 0.51148339 |
| 136 | AGTCGGCGCCAGGACG/CGTCCTGGCGCCGACT | 2.31975E-05 | 0.03041958 | 0.230796863 |

(b) Contingency table for Fisher's exact test to assess the association between the presence/absence of a specific sequence motif and nucleosomal or inter-nucleosomal DNA. Odds ratios greater than 1 indicate sequence motifs enriched in nucleosomal DNA.

Odds Ratio = (a*d)/(c*b)

| | a given sequence motif | other sequence motifs |
|---|---|---|
| Nucleosomal DNA | a | b |
| Inter-nucleosomal DNA | c | d |

p-values have been adjusted using the Benjamini–Hochberg method.

**Table S4.** (a) Identified sequence motifs significantly enriched or depleted in flanking linker regions of nucleosomal DNA

| Row No. | Sequence Motif | P-value | Adjusted P-value | Odds Ratio |
|---|---|---|---|---|
| 1 | GCCAGGGTCTT/AAGACCCTGGC | 0.001659946 | 0.049253731 | 0.811397454 |
| 2 | TAGA/TCTA | 1.8452E-157 | 0.026865672 | 0.796510654 |
| 3 | GATGATG/CATCATC | 9.13482E-38 | 0.032835821 | 0.795332527 |
| 4 | TTATACTCCAG/CTGGAGTATAA | 3.21506E-05 | 0.046268657 | 0.785311583 |
| 5 | GAAT/ATTC | 1.9862E-202 | 0.025373134 | 0.780967839 |
| 6 | GATG/CATC | 4.6506E-137 | 0.028358209 | 0.779195909 |
| 7 | ATG/CAT | 0 | 0.008208955 | 0.777487398 |
| 8 | CACCAT/ATGGTG | 7.0839E-157 | 0.02761194 | 0.776210992 |
| 9 | ACATG/CATGT | 0 | 0.019402985 | 0.773424329 |
| 10 | TA | 0 | 0.005970149 | 0.772134862 |
| 11 | TGTAT/ATACA | 0 | 0.018656716 | 0.772089234 |
| 12 | TTAGGGAC/GTCCCTAA | 1.28868E-10 | 0.041044776 | 0.771811829 |
| 13 | ACAGCA/TGCTGT | 2.2598E-212 | 0.024626866 | 0.771789579 |
| 14 | ACCATGCTTG/CAAGCATGGT | 1.52876E-12 | 0.040298507 | 0.769203303 |
| 15 | CCTGAACTG/CAGTTCAGG | 3.87208E-05 | 0.047014925 | 0.769100562 |
| 16 | AGGAC/GTCCT | 0 | 0.017164179 | 0.768966142 |
| 17 | AC/GT | 0 | 0.006716418 | 0.76806329 |
| 18 | AG/CT | 0 | 0.002238806 | 0.766799014 |
| 19 | TAGCGTC/GACGCTA | 1.72828E-15 | 0.036567164 | 0.766535959 |
| 20 | CTCTTGG/CCAAGAG | 5.74734E-57 | 0.032089552 | 0.765950264 |
| 21 | AAGAG/CTCTT | 0 | 0.014925373 | 0.765813989 |
| 22 | TTCTAC/GTAGAA | 4.3428E-255 | 0.02238806 | 0.761974934 |
| 23 | TC/GA | 0 | 0.005223881 | 0.760640164 |
| 24 | TG/CA | 0 | 0.001492537 | 0.75972113 |
| 25 | GTTCT/AGAAC | 0 | 0.020895522 | 0.75954026 |
| 26 | TAA/TTA | 0 | 0.008955224 | 0.757127789 |
| 27 | TTGCAGA/TCTGCAA | 2.37124E-83 | 0.031343284 | 0.757024925 |
| 28 | GTATGAGTCAA/TTGACTCATAC | 4.87245E-06 | 0.045522388 | 0.756198595 |
| 29 | TCGAG/CTCGA | 0 | 0.017910448 | 0.754745925 |
| 30 | CAA/TTG | 0 | 0.01119403 | 0.754146153 |
| 31 | ATAAGC/GCTTAT | 5.0953E-228 | 0.023134328 | 0.751954328 |
| 32 | GGT/ACC | 0 | 0.011940299 | 0.749611038 |
| 33 | CAACG/CGTTG | 0 | 0.015671642 | 0.749525643 |
| 34 | TGTTTAAGCAC/GTGCTTAAACA | 3.3141E-07 | 0.044776119 | 0.748913754 |
| 35 | TTGC/GCAA | 2.4006E-186 | 0.026119403 | 0.748620747 |
| 36 | TCAAGGATT/AATCCTTGA | 1.05293E-07 | 0.043283582 | 0.74816716 |
| 37 | TAAT/ATTA | 0 | 0.014179104 | 0.747503794 |
| 38 | TT/AA | 0 | 0.003731343 | 0.746268122 |
| 39 | CCGCAA/TTGCGG | 1.8336E-110 | 0.029850746 | 0.744071925 |
| 40 | AAACAA/TTGTTT | 0 | 0.021641791 | 0.741192194 |
| 41 | AAATG/CATTT | 0 | 0.01641791 | 0.741054295 |
| 42 | AAT/ATT | 0 | 0.012686567 | 0.740924057 |
| 43 | TCC/GGA | 0 | 0.013432836 | 0.740557362 |
| 44 | ATCGGTG/CACCGAT | 2.62817E-25 | 0.035074627 | 0.740266536 |
| 45 | TAGTTTC/GAAACTA | 3.5788E-112 | 0.029104478 | 0.738403762 |
| 46 | CATTTGGCAT/ATGCCAAATG | 1.39062E-27 | 0.034328358 | 0.73620126 |
| 47 | GCTCGA/TCGAGC | 8.27597E-97 | 0.030597015 | 0.734616622 |
| 48 | TCGTCTGTC/GACAGACGA | 0.001074206 | 0.048507463 | 0.733641471 |
| 49 | CAGCGCAA/TTGCGCTG | 8.06183E-10 | 0.041791045 | 0.732535519 |
| 50 | TCGA | 9.51963E-32 | 0.03358209 | 0.731927064 |
| 51 | TAGTAAAAG/CTTTTACTA | 5.58064E-14 | 0.039552239 | 0.730844106 |
| 52 | GG/CC | 0 | 0.000746269 | 0.730221498 |
| 53 | ATAAACGG/CCGTTTAT | 4.32278E-14 | 0.03880597 | 0.727070798 |
| 54 | CTCCGCAG/CTGCGGAG | 8.85925E-18 | 0.035820896 | 0.722190106 |
| 55 | GCATTGCG/CGCAATGC | 9.69985E-08 | 0.042537313 | 0.717573679 |
| 56 | CCGCA/TGCGG | 0 | 0.020149254 | 0.711360731 |
| 57 | TCTTCGATG/CATCGAAGA | 0.000139911 | 0.047761194 | 0.710870893 |
| 58 | TCGGGCTG/CAGCCCGA | 8.36297E-15 | 0.037313433 | 0.707252711 |
| 59 | AATATCGCAC/GTGCGATATT | 1.04805E-14 | 0.038059701 | 0.706333721 |
| 60 | TCG/CGA | 1.4419E-214 | 0.023880597 | 0.69876802 |
| 61 | GGC/GCC | 0 | 0.010447761 | 0.694873023 |

| | | | | |
|---|---|---|---|---|
| 62 | ATGAAAGTGGTTCTT/AAGAACCACTTTCAT | 0.002779112 | 0.05 | 0.671880625 |
| 63 | GTGCAGCGC/GCGCTGCAC | 2.39646E-07 | 0.044029851 | 0.646393626 |
| 64 | GCG/CGC | 0 | 0.009701493 | 0.644371853 |

(b) Contingency table for Fisher's exact test to assess the association between the presence/absence of a specific sequence motif and flanking linker regions of nucleosomal or inter-nucleosomal DNA. Odds ratios less than 1 indicate sequence motifs depleted in nucleosomal DNA.

Odds Ratio = (a*d)/(c*b)

| | a given sequence motif | other sequence motifs |
|---|---|---|
| Linker region of nucleosomal DNA | a | b |
| Linker region of inter-nucleosomal DNA | c | d |

p-values have been adjusted using the Benjamini–Hochberg method.

**Table S5.** (a) Informative features associated with the periodicity of sequence patterns and their statistical association with the nucleosomal structural features.

| Sequence Motif | Elements of contingency table | | | | P-value | Odds ratio |
|---|---|---|---|---|---|---|
| | a | b | c | d | | |
| AAAC/GTTT | 17722 | 20747 | 20041 | 21030 | 1.42102E-14 | 0.896349354 |
| AACA/TGTT | 18651 | 23186 | 19725 | 21390 | 1.13598E-22 | 0.872308417 |
| AACC/GGTT | 9669 | 11527 | 9602 | 9927 | 7.91846E-13 | 0.867204629 |
| AATG/CATT | 19163 | 24043 | 21085 | 21597 | 1.07062E-49 | 0.816384341 |
| ACAC/GTGT | 11858 | 14583 | 11396 | 11793 | 1.1104E-21 | 0.84146572 |
| ACAT/ATGT | 16661 | 20007 | 17527 | 18720 | 3.04191E-15 | 0.88944142 |
| ACCT/AGGT | 12825 | 13982 | 11427 | 11925 | 0.014847068 | 0.957225449 |
| ACTC/GAGT | 10895 | 12185 | 10066 | 10470 | 0.000161361 | 0.93001822 |
| AGCA/TGCT | 17091 | 20227 | 14964 | 15443 | 9.2537E-19 | 0.872007001 |
| AGCC/GGCT | 12661 | 14856 | 11803 | 12152 | 1.58973E-13 | 0.877448168 |
| AGCT | 14975 | 17410 | 13010 | 14158 | 6.18275E-05 | 0.936036257 |
| AGTC/GACT | 10043 | 11637 | 9427 | 9670 | 8.80073E-10 | 0.885269283 |
| AGTG/CACT | 13864 | 17036 | 12467 | 13416 | 4.19108E-15 | 0.875753756 |
| ATCA/TGAT | 14680 | 17339 | 14713 | 16025 | 4.25927E-07 | 0.922144143 |
| ATGC/GCAT | 11294 | 13486 | 9875 | 10765 | 1.45511E-06 | 0.912938575 |
| ATGG/CCAT | 13863 | 15772 | 12294 | 12875 | 1.41809E-06 | 0.920501464 |
| CAAA/TTTG | 25659 | 22787 | 23785 | 26272 | 1.55014E-65 | 1.243777093 |
| CAAC/GTTG | 9678 | 11000 | 8763 | 9000 | 7.598E-07 | 0.903613333 |
| CAGA/TCTG | 22257 | 20920 | 17875 | 18716 | 3.2152E-14 | 1.11396599 |
| CAGG/CCTG | 19920 | 20017 | 17195 | 16464 | 0.001101799 | 0.952847771 |
| CATC/GATG | 12828 | 13854 | 10230 | 11477 | 0.037789134 | 1.038810943 |
| CATG | 14447 | 16846 | 12283 | 12978 | 6.19566E-09 | 0.906116825 |
| CCAG/CTGG | 20516 | 18723 | 16361 | 16874 | 2.51193E-16 | 1.130122322 |
| CCCT/AGGG | 17088 | 15915 | 13978 | 15173 | 1.76965E-21 | 1.165496608 |
| CCTA/TAGG | 10568 | 9619 | 8821 | 9052 | 5.3789E-09 | 1.127430042 |
| CCTC/GAGG | 16752 | 16359 | 14609 | 15322 | 7.66224E-06 | 1.074001483 |
| CTAA/TTAG | 14244 | 13203 | 12874 | 15340 | 1.7809E-49 | 1.285497382 |
| CTCT/AGAG | 21432 | 19986 | 17451 | 18246 | 2.45133E-15 | 1.121202789 |
| CTTC/GAAG | 18480 | 18323 | 15620 | 16892 | 1.18724E-08 | 1.09070029 |
| CTTT/AAAG | 26972 | 25327 | 26745 | 28181 | 4.19531E-21 | 1.122130065 |
| GAAA/TTTC | 29352 | 26338 | 29971 | 30327 | 1.75138E-24 | 1.127672846 |
| GAAC/GTTC | 9329 | 11089 | 8744 | 9572 | 5.59947E-05 | 0.9209483 |
| GAGC/GCTC | 9878 | 12936 | 10058 | 10636 | 1.39435E-28 | 0.807487322 |
| GCCC/GGGC | 11665 | 11779 | 12062 | 13517 | 8.73759E-09 | 1.109781066 |
| GCTG/CAGC | 16749 | 17252 | 13412 | 14874 | 4.61691E-06 | 1.076672611 |
| GCTT/AAGC | 13611 | 14834 | 12199 | 12802 | 0.029467076 | 0.962909233 |
| GGAG/CTCC | 15467 | 18718 | 15590 | 15857 | 1.29298E-28 | 0.840468718 |
| GGCA/TGCC | 13156 | 15952 | 11241 | 11962 | 1.44202E-13 | 0.877622147 |
| GGCC | 10228 | 12759 | 11739 | 12478 | 4.73975E-18 | 0.852094889 |
| GGGA/TCCC | 14764 | 16997 | 14148 | 15318 | 0.000154824 | 0.940456638 |
| GGGG/CCCC | 12094 | 15084 | 15117 | 16422 | 9.09867E-17 | 0.870991417 |
| GGGT/ACCC | 9118 | 11798 | 9713 | 10249 | 1.01302E-24 | 0.81549124 |
| GGTC/GACC | 7284 | 8763 | 7275 | 7409 | 3.48489E-13 | 0.846532668 |
| GGTG/CACC | 11308 | 14048 | 11169 | 11590 | 8.41463E-23 | 0.835296086 |
| GTCC/GGAC | 9293 | 8652 | 8329 | 8468 | 4.29605E-05 | 1.092012007 |
| GTCT/AGAC | 12499 | 11850 | 10418 | 10846 | 6.22686E-07 | 1.098100691 |
| TCAC/GTGA | 13234 | 14388 | 12104 | 12025 | 3.18757E-07 | 0.913790989 |
| TCCT/AGGA | 21413 | 19705 | 17277 | 18302 | 2.70492E-22 | 1.151148351 |
| TCTA/TAGA | 14827 | 13934 | 14797 | 15121 | 3.96415E-07 | 1.087387462 |

| | | | | | | |
|---|---|---|---|---|---|---|
| TGAC/GTCA | 10890 | 13056 | 10156 | 10193 | 1.41141E-20 | 0.837138027 |
| TGAG/CTCA | 16700 | 19020 | 15218 | 15261 | 3.73876E-16 | 0.880504077 |
| AAGCA/TGCTT | 5191 | 6251 | 4764 | 5013 | 1.03529E-06 | 0.873831069 |
| AAGTG/CACTT | 3895 | 5156 | 4106 | 4029 | 1.68452E-22 | 0.741263943 |
| ACATG/CATGT | 3691 | 4951 | 3590 | 3560 | 5.39315E-21 | 0.739276104 |
| ACTCA/TGAGT | 3671 | 4376 | 3371 | 3694 | 0.010297988 | 0.919274493 |
| AGAGG/CCTCT | 5194 | 5962 | 4485 | 4801 | 0.013294566 | 0.932565258 |
| AGATG/CATCT | 4425 | 5606 | 4056 | 4251 | 1.91458E-10 | 0.827281553 |
| AGGGG/CCCCT | 3803 | 4329 | 3883 | 3988 | 0.001178734 | 0.902249186 |
| CAGAG/CTCTG | 6457 | 6515 | 4935 | 5487 | 0.000229152 | 1.101955786 |
| CCAGG/CCTGG | 6437 | 5570 | 5335 | 5341 | 4.46216E-08 | 1.156955002 |
| GAAGC/GCTTC | 3148 | 4215 | 2892 | 3355 | 3.51184E-05 | 0.866425809 |
| GACCC/GGGTC | 2015 | 2565 | 2288 | 2429 | 1.3725E-05 | 0.833986798 |
| GAGGC/GCCTC | 3142 | 4384 | 3351 | 3835 | 2.48012E-09 | 0.820212863 |
| GAGGT/ACCTC | 2761 | 3719 | 2948 | 2491 | 1.59993E-36 | 0.627316162 |
| GATGT/ACATC | 2945 | 3729 | 2744 | 2949 | 6.03906E-06 | 0.848757415 |
| GCATG/CATGC | 2734 | 3618 | 2409 | 2389 | 6.16607E-14 | 0.749392423 |
| GGAAA/TTTCC | 7756 | 6878 | 6755 | 7511 | 7.76583E-22 | 1.253857083 |
| GGAGG/CCTCC | 4944 | 5920 | 5204 | 5379 | 8.22105E-08 | 0.863219041 |
| GGGAC/GTCCC | 2267 | 3033 | 2511 | 2967 | 0.001376757 | 0.88318146 |
| GGGGC/GCCCC | 2793 | 4001 | 4125 | 4411 | 4.54016E-19 | 0.746475381 |
| GTGTC/GACAC | 2471 | 2895 | 2022 | 2567 | 0.047737118 | 1.083599746 |
| TCCCA/TGGGA | 5677 | 4839 | 4070 | 4542 | 2.28291E-20 | 1.309230134 |
| TCTAG/CTAGA | 3342 | 2912 | 2657 | 3093 | 2.50198E-15 | 1.335990717 |
| TGACC/GGTCA | 2463 | 3203 | 2428 | 2232 | 2.36889E-18 | 0.706891861 |
| TGAGC/GCTCA | 3055 | 3958 | 2910 | 2794 | 6.0408E-17 | 0.74108639 |
| TGATG/CATCA | 3832 | 4945 | 3361 | 3762 | 9.09087E-06 | 0.867380158 |
| TGGGC/GCCCA | 3365 | 4197 | 3132 | 3505 | 0.001339978 | 0.89724773 |
| TTAGC/GCTAA | 2424 | 2856 | 2451 | 2266 | 1.60093E-09 | 0.784677151 |
| TTGAC/GTCAA | 2458 | 3105 | 2581 | 2448 | 2.0426E-13 | 0.750833572 |
| GGAGGT/ACCTCC | 900 | 1053 | 870 | 859 | 0.010936991 | 0.843894292 |

(b) Contingency table used in Fisher's exact test to assess the association between the presence or absence of a specific sequence motif and minor groove sites. An odds ratio greater than 1 indicates that the sequence motif is enriched in minor groove facing out sites, whereas an odds ratio less than 1 indicates it is depleted.

Odds Ratio = (a*d)/(c*b)

| | Minor grooves facing out the histone octamer | Minor grooves facing the histone octamer |
|---|---|---|
| Nucleosomal DNA | a | b |
| Inter-nucleosomal DND | c | d |

**Table S6.** (a) Quality control-passed single nucleotide mutations in nucleosomal and non-nucleosomal regions.

| Chromosome # | Single nucleotide mutation | Elements of contingency table | | | | Odds ratio | P-value |
|---|---|---|---|---|---|---|---|
| | | a | b | c | d | | |
| 1 | T->A | 4750 | 177944 | 8710 | 308568 | 0.945677359 | 0.002231075 |
| 2 | | 6064 | 228804 | 4780 | 153324 | 0.850115284 | 1.64504E-16 |
| 3 | | 4591 | 169515 | 2970 | 96458 | 0.879591375 | 8.91164E-08 |
| 4 | | 3490 | 123352 | 2341 | 76983 | 0.930406315 | 0.008078103 |
| 5 | | 3983 | 151203 | 2765 | 91367 | 0.870450598 | 3.7805E-08 |
| 6 | | 1757 | 74669 | 4726 | 169476 | 0.843812422 | 1.27581E-09 |
| 7 | | 2028 | 76596 | 6329 | 220623 | 0.922948674 | 0.001880551 |
| 8 | | 3343 | 118903 | 2261 | 75105 | 0.93392468 | 0.013834995 |
| 9 | | 1311 | 56327 | 3053 | 132235 | 1.008104807 | 0.814854703 |
| 10 | | 1111 | 39921 | 4650 | 170382 | 1.019725797 | 0.562731421 |
| 11 | | 2618 | 107256 | 4760 | 169114 | 0.867202767 | 6.10534E-09 |
| 12 | | 4379 | 163999 | 2649 | 94353 | 0.951059089 | 0.044610736 |
| 13 | | 1421 | 58389 | 1253 | 40261 | 0.781981587 | 4.84353E-10 |
| 14 | | 2389 | 94381 | 1868 | 61502 | 0.833381667 | 7.23632E-09 |
| 15 | | 2078 | 84614 | 2361 | 85949 | 0.894021901 | 0.000247384 |
| 16 | | 779 | 29617 | 3906 | 165594 | 1.115086994 | 0.006572925 |
| 17 | | 2832 | 129756 | 2693 | 112795 | 0.914153855 | 0.001011694 |
| 18 | | 714 | 23056 | 659 | 19909 | 0.935574293 | 0.226596899 |
| 19 | | 1893 | 90605 | 1205 | 51685 | 0.896140124 | 0.003432464 |
| 20 | | 764 | 30412 | 1217 | 31915 | 0.658798569 | 2.23313E-19 |
| 21 | | 435 | 14465 | 690 | 21206 | 0.924230902 | 0.206204911 |
| 22 | | 652 | 30658 | 944 | 34280 | 0.7722761 | 4.87437E-07 |
| 1 | T->C | 10967 | 171727 | 20570 | 296708 | 0.921179416 | 1.50029E-11 |
| 2 | | 14288 | 220580 | 10646 | 147458 | 0.897195706 | 2.9526E-16 |
| 3 | | 10814 | 163292 | 6796 | 92632 | 0.902670247 | 1.93601E-10 |
| 4 | | 8370 | 118472 | 5552 | 73772 | 0.938754081 | 0.000435225 |
| 5 | | 9379 | 145807 | 6122 | 88010 | 0.924734067 | 4.34537E-06 |
| 6 | | 4690 | 71736 | 11873 | 162329 | 0.893863763 | 2.41733E-10 |
| 7 | | 4299 | 74325 | 13083 | 213869 | 0.945525018 | 0.001935191 |
| 8 | | 7161 | 115085 | 4754 | 72612 | 0.950395067 | 0.008597905 |
| 9 | | 3656 | 53982 | 7984 | 127304 | 1.079888056 | 0.000208601 |
| 10 | | 2438 | 38594 | 11233 | 163799 | 0.921147963 | 0.000340337 |
| 11 | | 6131 | 103743 | 10466 | 163408 | 0.922709724 | 1.16442E-06 |
| 12 | | 10684 | 157694 | 6424 | 90578 | 0.955291481 | 0.005264056 |
| 13 | | 3845 | 55965 | 2801 | 38713 | 0.949562499 | 0.044158272 |
| 14 | | 5821 | 90949 | 4233 | 59137 | 0.89415064 | 9.22124E-08 |
| 15 | | 5288 | 81404 | 5557 | 82753 | 0.967362062 | 0.095764202 |
| 16 | | 1594 | 28802 | 9423 | 160077 | 0.940167894 | 0.027029536 |
| 17 | | 7503 | 125085 | 7046 | 108442 | 0.923176187 | 3.03366E-06 |
| 18 | | 1414 | 22356 | 1287 | 19281 | 0.947559121 | 0.175885659 |
| 19 | | 4449 | 88049 | 3019 | 49871 | 0.834685639 | 1.23682E-13 |
| 20 | | 1782 | 29394 | 1942 | 31190 | 0.973677555 | 0.43718215 |
| 21 | | 838 | 14062 | 1347 | 20549 | 0.909117508 | 0.036644286 |
| 22 | | 1593 | 29717 | 2091 | 33133 | 0.849410332 | 1.80906E-06 |
| 1 | T->G | 5615 | 177079 | 11524 | 305754 | 0.841301373 | 6.29724E-26 |
| 2 | | 8697 | 226171 | 6138 | 151966 | 0.952033214 | 0.003918576 |
| 3 | | 6508 | 167598 | 4028 | 95400 | 0.91968187 | 4.50512E-05 |
| 4 | | 4993 | 121849 | 3607 | 75717 | 0.860175073 | 1.87538E-11 |

| | | | | | | | |
|---|---|---|---|---|---|---|---|
| 5 | | 5454 | 149732 | 3541 | 90591 | 0.931879236 | 0.001368147 |
| 6 | | 2369 | 74057 | 7696 | 166506 | 0.692091913 | 1.78495E-56 |
| 7 | | 2135 | 76489 | 7237 | 219715 | 0.847422578 | 1.93308E-11 |
| 8 | | 4054 | 118192 | 2910 | 74456 | 0.87761164 | 1.44876E-07 |
| 9 | | 2117 | 55521 | 4094 | 131194 | 1.221883256 | 3.44625E-13 |
| 10 | | 1484 | 39548 | 6347 | 168685 | 0.997280535 | 0.941512466 |
| 11 | | 3187 | 106687 | 5076 | 168798 | 0.993381902 | 0.774508555 |
| 12 | | 5810 | 162568 | 3547 | 93455 | 0.941634632 | 0.005695731 |
| 13 | | 2276 | 57534 | 1681 | 39833 | 0.937395775 | 0.049750667 |
| 14 | | 3176 | 93594 | 2314 | 61056 | 0.895359569 | 7.47535E-05 |
| 15 | | 2939 | 83753 | 2877 | 85433 | 1.042041485 | 0.125095667 |
| 16 | | 799 | 29597 | 4299 | 165201 | 1.03739539 | 0.343042957 |
| 17 | | 3313 | 129275 | 3302 | 112186 | 0.870699879 | 2.88399E-08 |
| 18 | | 783 | 22987 | 776 | 19792 | 0.868775224 | 0.006626863 |
| 19 | | 1910 | 90588 | 1189 | 51701 | 0.916810935 | 0.0202782 |
| 20 | | 879 | 30297 | 1020 | 32112 | 0.913390376 | 0.053084972 |
| 21 | | 558 | 14342 | 730 | 21166 | 1.128081332 | 0.037468014 |
| 22 | | 675 | 30635 | 887 | 34337 | 0.852952049 | 0.002079598 |
| 1 | | 25029 | 157665 | 41272 | 276006 | 1.061625182 | 4.14121E-12 |
| 2 | | 32245 | 202623 | 21967 | 136137 | 0.986231946 | 0.142407103 |
| 3 | | 23382 | 150724 | 13267 | 86161 | 1.007481886 | 0.524785464 |
| 4 | | 17392 | 109450 | 10717 | 68607 | 1.017252953 | 0.196101081 |
| 5 | | 21441 | 133745 | 12794 | 81338 | 1.019188758 | 0.114470141 |
| 6 | | 9959 | 66467 | 23719 | 150483 | 0.950606371 | 7.60349E-05 |
| 7 | | 11245 | 67379 | 32962 | 193990 | 0.982201712 | 0.129119731 |
| 8 | | 17287 | 104959 | 10711 | 66655 | 1.024949956 | 0.063064169 |
| 9 | | 7526 | 50112 | 17373 | 117915 | 1.019334477 | 0.196791634 |
| 10 | | 5872 | 35160 | 23691 | 151341 | 1.066867258 | 4.23941E-05 |
| 11 | C->A | 15674 | 94200 | 23861 | 150013 | 1.046090348 | 4.97841E-05 |
| 12 | | 22558 | 145820 | 13427 | 83575 | 0.962899353 | 0.001307605 |
| 13 | | 8123 | 51687 | 5863 | 35651 | 0.955623783 | 0.01413242 |
| 14 | | 12752 | 84018 | 8388 | 54982 | 0.994873989 | 0.734107841 |
| 15 | | 11592 | 75100 | 11500 | 76810 | 1.030951798 | 0.031212247 |
| 16 | | 4011 | 26385 | 21016 | 148484 | 1.074051639 | 0.000123092 |
| 17 | | 15958 | 116630 | 13550 | 101938 | 1.029354574 | 0.020069501 |
| 18 | | 3214 | 20556 | 2887 | 17681 | 0.957562883 | 0.118305974 |
| 19 | | 11184 | 81314 | 6503 | 46387 | 0.981102461 | 0.25330267 |
| 20 | | 4156 | 27020 | 4284 | 28848 | 1.035753576 | 0.13477868 |
| 21 | | 2195 | 12705 | 3061 | 18835 | 1.063070704 | 0.043563245 |
| 22 | | 3853 | 27457 | 4307 | 30917 | 1.007322187 | 0.758237096 |
| 1 | | 38164 | 144530 | 65112 | 252166 | 1.022636709 | 0.002023189 |
| 2 | | 48197 | 186671 | 29787 | 128317 | 1.112245314 | 1.61449E-38 |
| 3 | | 36024 | 138082 | 19358 | 80070 | 1.079106251 | 1.83199E-14 |
| 4 | | 24722 | 102120 | 14568 | 64756 | 1.07610061 | 2.30924E-10 |
| 5 | | 32340 | 122846 | 18532 | 75600 | 1.073936244 | 4.20337E-12 |
| 6 | | 16701 | 59725 | 33088 | 141114 | 1.192575555 | 1.77097E-60 |
| 7 | | 16517 | 62107 | 45343 | 181609 | 1.065167074 | 7.08648E-10 |
| 8 | | 24969 | 97277 | 15211 | 62155 | 1.048840105 | 3.35749E-05 |
| 9 | C->T | 12292 | 45346 | 30523 | 104765 | 0.930406142 | 2.08196E-09 |
| 10 | | 8045 | 32987 | 35450 | 139582 | 0.960276731 | 0.003271224 |
| 11 | | 23697 | 86177 | 36398 | 137476 | 1.038607284 | 5.8616E-05 |
| 12 | | 34877 | 133501 | 19096 | 77906 | 1.065818129 | 2.30772E-10 |
| 13 | | 12278 | 47532 | 7730 | 33784 | 1.128945859 | 5.51771E-14 |
| 14 | | 20853 | 75917 | 12517 | 50853 | 1.115952814 | 4.08961E-18 |
| 15 | | 18572 | 68120 | 18793 | 69517 | 1.008507062 | 0.469453852 |
| 16 | | 6969 | 23427 | 39884 | 129616 | 0.966748917 | 0.022226676 |

| | | | | | | | |
|---|---|---|---|---|---|---|---|
| 17 | | 31977 | 100611 | 27096 | 88392 | 1.036812032 | 0.000132102 |
| 18 | | 4961 | 18809 | 3933 | 16635 | 1.115584263 | 4.68457E-06 |
| 19 | | 23501 | 68997 | 12467 | 40423 | 1.104390639 | 5.50767E-15 |
| 20 | | 6779 | 24397 | 6111 | 27021 | 1.228622152 | 1.61363E-25 |
| 21 | | 2954 | 11946 | 4154 | 17742 | 1.056146255 | 0.04225715 |
| 22 | | 7834 | 23476 | 8000 | 27224 | 1.135589623 | 3.20525E-12 |
| 1 | C->G | 6822 | 175872 | 11451 | 305827 | 1.035970581 | 0.023721395 |
| 2 | | 7943 | 226925 | 5734 | 152370 | 0.930130739 | 4.28529E-05 |
| 3 | | 5734 | 168372 | 3295 | 96133 | 0.993584698 | 0.772404138 |
| 4 | | 4454 | 122388 | 2877 | 76447 | 0.967012215 | 0.171046818 |
| 5 | | 4996 | 150190 | 3312 | 90820 | 0.912163273 | 5.8893E-05 |
| 6 | | 2737 | 73689 | 5999 | 168203 | 1.041422628 | 0.084203859 |
| 7 | | 3088 | 75536 | 8522 | 218430 | 1.047837808 | 0.02959974 |
| 8 | | 4309 | 117937 | 2836 | 74530 | 0.960177031 | 0.100075758 |
| 9 | | 1917 | 55721 | 4617 | 130671 | 0.973694128 | 0.342781566 |
| 10 | | 1566 | 39466 | 6145 | 168887 | 1.090543466 | 0.002822885 |
| 11 | | 3630 | 106244 | 6376 | 167498 | 0.897560081 | 3.05203E-07 |
| 12 | | 5881 | 162497 | 3358 | 93644 | 1.00926473 | 0.684151054 |
| 13 | | 1962 | 57848 | 1429 | 40085 | 0.951393793 | 0.160636692 |
| 14 | | 3394 | 93376 | 2365 | 61005 | 0.937585449 | 0.018939387 |
| 15 | | 2877 | 83815 | 3067 | 85243 | 0.954032264 | 0.074825059 |
| 16 | | 1046 | 29350 | 6222 | 163278 | 0.935236061 | 0.049590505 |
| 17 | | 4711 | 127877 | 4057 | 111431 | 1.011862929 | 0.593303375 |
| 18 | | 799 | 22971 | 742 | 19826 | 0.929390169 | 0.160420563 |
| 19 | | 3312 | 89186 | 2062 | 50828 | 0.915393875 | 0.002087262 |
| 20 | | 1228 | 29948 | 1992 | 31140 | 0.641002638 | 9.77246E-34 |
| 21 | | 470 | 14430 | 966 | 20930 | 0.705705706 | 6.4077E-10 |
| 22 | | 1048 | 30262 | 1383 | 33841 | 0.847392593 | 7.02262E-05 |

(b) Contingency table used in Fisher's exact test to assess the association between mutations and genomic regions, including nucleosomal (well-positioned and non-well-positioned) and non-nucleosomal regions. An odds ratio greater than 1 indicates that a specific mutation is enriched in nucleosomal DNA, whereas an odds ratio less than 1 indicates depletion.

Odds Ratio = (a*d)/(c*b)

| | Specific single nucleotide mutation | Other mutations |
|---|---|---|
| All nucleosomal regions | a | b |
| Inter-ncleosomal regions | c | d |

**Table S7.** (a) Mutect2 quality-passed single nucleotide mutations in nucleosomal and non-nucleosomal regions.

| Chromosome # | Single nucleotide mutation | Elements of contingency table | | | | Odds ratio | P-value |
|---|---|---|---|---|---|---|---|
| | | a | b | c | d | | |
| 1 | T->A | 4301 | 154943 | 9159 | 331569 | 1.004901148 | 0.792890283 |
| 2 | | 5375 | 196985 | 5469 | 185143 | 0.923729227 | 4.64703E-05 |
| 3 | | 4111 | 146095 | 3450 | 119878 | 0.977760566 | 0.336590035 |
| 4 | | 2945 | 102919 | 2886 | 97416 | 0.965881188 | 0.192795244 |
| 5 | | 3587 | 130543 | 3161 | 112027 | 0.973813964 | 0.287125683 |
| 6 | | 1580 | 64350 | 4903 | 179795 | 0.90037671 | 0.000316278 |
| 7 | | 1810 | 65512 | 6547 | 231707 | 0.977810125 | 0.414357734 |
| 8 | | 2948 | 101392 | 2656 | 92616 | 1.013868754 | 0.61578033 |
| 9 | | 1192 | 48532 | 3172 | 140030 | 1.084266342 | 0.019001943 |
| 10 | | 957 | 33353 | 4804 | 176950 | 1.056877379 | 0.124862764 |
| 11 | | 2367 | 93241 | 5011 | 183129 | 0.927735246 | 0.002976425 |
| 12 | | 3912 | 141686 | 3116 | 116666 | 1.033757719 | 0.173687092 |
| 13 | | 1253 | 49609 | 1421 | 49041 | 0.871677503 | 0.000483732 |
| 14 | | 2148 | 81032 | 2109 | 74851 | 0.940803114 | 0.050132624 |
| 15 | | 1870 | 72328 | 2569 | 98235 | 0.988637991 | 0.723501907 |
| 16 | | 724 | 25980 | 3961 | 169231 | 1.19062363 | 2.98911E-05 |
| 17 | | 2487 | 112641 | 3038 | 129910 | 0.944135008 | 0.035668341 |
| 18 | | 626 | 19526 | 747 | 23439 | 1.005957251 | 0.912352918 |
| 19 | | 1680 | 78018 | 1418 | 64272 | 0.976023001 | 0.511298271 |
| 20 | | 642 | 26242 | 1339 | 36085 | 0.659301752 | 2.7019E-18 |
| 21 | | 371 | 12369 | 754 | 23302 | 0.92696038 | 0.25192754 |
| 22 | | 549 | 26485 | 1047 | 38453 | 0.761300147 | 2.4052E-07 |
| 1 | T->C | 9339 | 149905 | 22198 | 318530 | 0.893965484 | 7.92325E-19 |
| 2 | | 12043 | 190317 | 12891 | 177721 | 0.872387116 | 1.89002E-25 |
| 3 | | 9128 | 141078 | 8482 | 114846 | 0.876060184 | 2.39485E-17 |
| 4 | | 6823 | 99041 | 7099 | 93203 | 0.904467716 | 1.08698E-08 |
| 5 | | 7844 | 126286 | 7657 | 107531 | 0.872283013 | 1.88938E-16 |
| 6 | | 3912 | 62018 | 12651 | 172047 | 0.857834163 | 2.35567E-16 |
| 7 | | 3528 | 63794 | 13854 | 224400 | 0.89576979 | 1.06226E-08 |
| 8 | | 6028 | 98312 | 5887 | 89385 | 0.930973504 | 0.00015489 |
| 9 | | 3094 | 46630 | 8546 | 134656 | 1.04548478 | 0.040947206 |
| 10 | | 2048 | 32262 | 11623 | 170131 | 0.92918862 | 0.002937557 |
| 11 | | 5119 | 90489 | 11478 | 176662 | 0.870695426 | 7.71675E-16 |
| 12 | | 8972 | 136626 | 8136 | 111646 | 0.901131434 | 5.06037E-11 |
| 13 | | 3200 | 47662 | 3446 | 47016 | 0.916026675 | 0.000556711 |
| 14 | | 4808 | 78372 | 5246 | 71714 | 0.838646985 | 1.45841E-17 |
| 15 | | 4481 | 69717 | 6364 | 94440 | 0.953810416 | 0.018908377 |
| 16 | | 1368 | 25336 | 9649 | 163543 | 0.915161414 | 0.002730362 |
| 17 | | 6284 | 108844 | 8265 | 124683 | 0.870955705 | 9.76885E-16 |
| 18 | | 1170 | 18982 | 1531 | 22655 | 0.912079655 | 0.021849374 |
| 19 | | 3670 | 76028 | 3798 | 61892 | 0.786632806 | 6.46534E-24 |
| 20 | | 1400 | 25484 | 2324 | 35100 | 0.829719758 | 7.57445E-08 |
| 21 | | 695 | 12045 | 1490 | 22566 | 0.873868965 | 0.004345448 |
| 22 | | 1341 | 25693 | 2343 | 37157 | 0.827717838 | 6.51031E-08 |
| 1 | T->G | 4793 | 154451 | 12346 | 328382 | 0.825410094 | 3.00878E-29 |
| 2 | | 7247 | 195113 | 7588 | 183024 | 0.895886052 | 5.16564E-11 |
| 3 | | 5356 | 144850 | 5180 | 118148 | 0.843371039 | 1.12444E-17 |
| 4 | | 4126 | 101738 | 4474 | 95828 | 0.868645296 | 1.74212E-10 |

| | | | | | | | |
|---|---|---|---|---|---|---|---|
| 5 | | 4614 | 129516 | 4381 | 110807 | 0.901048381 | 1.31888E-06 |
| 6 | | 1993 | 63937 | 8072 | 176626 | 0.682069335 | 1.29307E-54 |
| 7 | | 1759 | 65563 | 7613 | 230641 | 0.812807504 | 3.67095E-15 |
| 8 | | 3402 | 100938 | 3562 | 91710 | 0.867765525 | 6.55048E-09 |
| 9 | | 1774 | 47950 | 4437 | 138765 | 1.157059028 | 4.54451E-07 |
| 10 | | 1196 | 33114 | 6635 | 175119 | 0.953261112 | 0.138783441 |
| 11 | | 2702 | 92906 | 5561 | 182579 | 0.954859584 | 0.052754217 |
| 12 | | 4903 | 140695 | 4454 | 115328 | 0.902334946 | 1.14166E-06 |
| 13 | | 1931 | 48931 | 2026 | 48436 | 0.943467646 | 0.07443789 |
| 14 | | 2665 | 80515 | 2825 | 74135 | 0.868610862 | 3.14351E-07 |
| 15 | | 2455 | 71743 | 3361 | 97443 | 0.99209691 | 0.776912919 |
| 16 | | 652 | 26052 | 4446 | 168746 | 0.949883962 | 0.234592945 |
| 17 | | 2826 | 112302 | 3789 | 129159 | 0.857797419 | 1.06062E-09 |
| 18 | | 672 | 19480 | 887 | 23299 | 0.906137246 | 0.058773605 |
| 19 | | 1589 | 78109 | 1510 | 64180 | 0.864660431 | 6.4339E-05 |
| 20 | | 716 | 26168 | 1183 | 36241 | 0.838219808 | 0.000228548 |
| 21 | | 445 | 12295 | 843 | 23213 | 0.996632797 | 0.976221528 |
| 22 | | 552 | 26482 | 1010 | 38490 | 0.794355366 | 1.48916E-05 |
| 1 | | 22270 | 136974 | 44031 | 296697 | 1.095561305 | 9.01706E-25 |
| 2 | | 28539 | 173821 | 25673 | 164939 | 1.054831872 | 8.31731E-09 |
| 3 | | 20781 | 129425 | 15868 | 107460 | 1.087358899 | 1.29701E-13 |
| 4 | | 14982 | 90882 | 13127 | 87175 | 1.094758656 | 1.95642E-12 |
| 5 | | 18964 | 115166 | 15271 | 99917 | 1.077401491 | 1.85274E-10 |
| 6 | | 8808 | 57122 | 24870 | 159828 | 0.990948263 | 0.497519104 |
| 7 | | 9967 | 57355 | 34240 | 204014 | 1.03542677 | 0.004851991 |
| 8 | | 15231 | 89109 | 12767 | 82505 | 1.104582754 | 1.44967E-14 |
| 9 | | 6616 | 43108 | 18283 | 124919 | 1.048621269 | 0.002111087 |
| 10 | | 5045 | 29265 | 24518 | 157236 | 1.105553054 | 2.63413E-09 |
| 11 | C->A | 14078 | 81530 | 25457 | 162683 | 1.103464761 | 5.46297E-18 |
| 12 | | 20028 | 125570 | 15957 | 103825 | 1.037773038 | 0.001164915 |
| 13 | | 7117 | 43745 | 6869 | 43593 | 1.032504103 | 0.080342724 |
| 14 | | 11298 | 71882 | 9842 | 67118 | 1.071857533 | 2.7125E-06 |
| 15 | | 10123 | 64075 | 12969 | 87835 | 1.069994973 | 2.08249E-06 |
| 16 | | 3671 | 23033 | 21356 | 151836 | 1.133153433 | 1.21189E-10 |
| 17 | | 14297 | 100831 | 15211 | 117737 | 1.097503824 | 7.0251E-14 |
| 18 | | 2792 | 17360 | 3309 | 20877 | 1.014698497 | 0.598875518 |
| 19 | | 9965 | 69733 | 7722 | 57968 | 1.072747406 | 1.3937E-05 |
| 20 | | 3770 | 23114 | 4670 | 32754 | 1.14396755 | 1.14857E-08 |
| 21 | | 1946 | 10794 | 3310 | 20746 | 1.12996971 | 8.47271E-05 |
| 22 | | 3401 | 23633 | 4759 | 34741 | 1.05054434 | 0.040811488 |
| 1 | | 32850 | 126394 | 70426 | 270302 | 0.997528132 | 0.744296051 |
| 2 | | 40993 | 161367 | 36991 | 153621 | 1.054992794 | 2.34508E-11 |
| 3 | | 30670 | 119536 | 24712 | 98616 | 1.023892931 | 0.013619108 |
| 4 | | 20341 | 85523 | 18949 | 81353 | 1.021119688 | 0.062553475 |
| 5 | | 27638 | 106492 | 23234 | 91954 | 1.027155675 | 0.007224075 |
| 6 | | 14259 | 51671 | 35530 | 149168 | 1.158571022 | 2.74184E-39 |
| 7 | | 14001 | 53321 | 47859 | 190395 | 1.044606416 | 5.44769E-05 |
| 8 | | 20851 | 83489 | 19329 | 75943 | 0.981241691 | 0.090431219 |
| 9 | C->T | 10566 | 39158 | 32249 | 110953 | 0.928352479 | 3.91468E-09 |
| 10 | | 6561 | 27749 | 36934 | 144820 | 0.927096409 | 3.50046E-07 |
| 11 | | 20299 | 75309 | 39796 | 148344 | 1.00475072 | 0.626937508 |
| 12 | | 29834 | 115764 | 24139 | 95643 | 1.021108464 | 0.031439437 |
| 13 | | 10216 | 40646 | 9792 | 40670 | 1.043916685 | 0.006477106 |
| 14 | | 17728 | 65452 | 15642 | 61318 | 1.061775062 | 1.14823E-06 |
| 15 | | 15742 | 58456 | 21623 | 79181 | 0.986133807 | 0.237845091 |
| 16 | | 6005 | 20699 | 40848 | 132344 | 0.939933452 | 7.81735E-05 |

| | | | | | | | |
|---|---|---|---|---|---|---|---|
| 17 | | 27598 | 87530 | 31475 | 101473 | 1.01649546 | 0.08368669 |
| 18 | | 4148 | 16004 | 4746 | 19440 | 1.061643565 | 0.011979828 |
| 19 | | 20042 | 59656 | 15926 | 49764 | 1.04977324 | 7.21098E-05 |
| 20 | | 5861 | 21023 | 7029 | 30395 | 1.205551133 | 5.46775E-21 |
| 21 | | 2512 | 10228 | 4596 | 19460 | 1.039900367 | 0.156955599 |
| 22 | | 6744 | 20290 | 9090 | 30410 | 1.111957149 | 9.61146E-09 |
| 1 | C->G | 6069 | 153175 | 12204 | 328524 | 1.066581758 | 6.01863E-05 |
| 2 | | 6983 | 195377 | 6694 | 183918 | 0.981990153 | 0.30011512 |
| 3 | | 5057 | 145149 | 3972 | 119356 | 1.046921024 | 0.034121949 |
| 4 | | 3715 | 102149 | 3616 | 96686 | 0.972433407 | 0.243650863 |
| 5 | | 4418 | 129712 | 3890 | 111298 | 0.974503301 | 0.249060945 |
| 6 | | 2413 | 63517 | 6323 | 178375 | 1.071712136 | 0.004623158 |
| 7 | | 2596 | 64726 | 9014 | 229240 | 1.019996697 | 0.385632224 |
| 8 | | 3710 | 100630 | 3435 | 91837 | 0.985683267 | 0.554553516 |
| 9 | | 1620 | 48104 | 4914 | 138288 | 0.947726812 | 0.065533431 |
| 10 | | 1348 | 32962 | 6363 | 175391 | 1.127253826 | 0.000107952 |
| 11 | | 3239 | 92369 | 6767 | 181373 | 0.939855671 | 0.004326701 |
| 12 | | 5150 | 140448 | 4089 | 115693 | 1.03748456 | 0.084766947 |
| 13 | | 1714 | 49148 | 1677 | 48785 | 1.014514397 | 0.687878495 |
| 14 | | 2943 | 80237 | 2816 | 74144 | 0.965737157 | 0.19726806 |
| 15 | | 2428 | 71770 | 3516 | 97288 | 0.936086852 | 0.014016118 |
| 16 | | 932 | 25772 | 6336 | 166856 | 0.952345314 | 0.176284513 |
| 17 | | 4072 | 111056 | 4696 | 128252 | 1.001386657 | 0.956531176 |
| 18 | | 668 | 19484 | 873 | 23313 | 0.91555041 | 0.095619112 |
| 19 | | 2903 | 76795 | 2471 | 63219 | 0.967139158 | 0.235200969 |
| 20 | | 1053 | 25831 | 2167 | 35257 | 0.663244407 | 1.72957E-27 |
| 21 | | 401 | 12339 | 1035 | 23021 | 0.722850097 | 3.8441E-08 |
| 22 | | 930 | 26104 | 1501 | 37999 | 0.901918641 | 0.015545636 |

(b) Contingency table used in Fisher's exact test to assess the association between Mutect2-identified mutations and genomic regions, including nucleosomal (well- and non-well-positioned) and non-nucleosomal regions. An odds ratio greater than 1 indicates enrichment of a specific mutation in nucleosomal DNA, whereas an odds ratio less than 1 indicates depletion.

Odds Ratio = (a*d)/(c*b)

| | Specific single nucleotide mutation | Other mutations |
|---|---|---|
| All nucleosomal regions | a | b |
| Inter-ncleosomal regions | c | d |

**Table S8.** (a) Mutect2 quality-passed single nucleotide mutations in well-positioned nucleosomes.

| Chromosome # | Single nucleotide mutation | Elements of contingency table | | | | Odds ratio | P-value |
|---|---|---|---|---|---|---|---|
| | | a | b | c | d | | |
| 1 | T->A | 1611 | 57419 | 11849 | 429093 | 1.016037287 | 0.551366907 |
| 2 | | 2066 | 76332 | 8778 | 305796 | 0.942887546 | 0.017501722 |
| 3 | | 1639 | 54725 | 5922 | 211248 | 1.068359426 | 0.020270199 |
| 4 | | 1142 | 40118 | 4689 | 160217 | 0.972646868 | 0.415934795 |
| 5 | | 1340 | 50126 | 5408 | 192444 | 0.951282354 | 0.109389864 |
| 6 | | 555 | 24065 | 5928 | 220080 | 0.85620843 | 0.000448124 |
| 7 | | 744 | 27672 | 7613 | 269547 | 0.951943251 | 0.214477804 |
| 8 | | 1124 | 39856 | 4480 | 154152 | 0.970384276 | 0.383101239 |
| 9 | | 478 | 18330 | 3886 | 170232 | 1.142362234 | 0.007261863 |
| 10 | | 415 | 13911 | 5346 | 196392 | 1.095934464 | 0.076554187 |
| 11 | | 838 | 33670 | 6540 | 242700 | 0.923619153 | 0.033192221 |
| 12 | | 1445 | 55029 | 5583 | 203323 | 0.956302096 | 0.135808631 |
| 13 | | 528 | 19708 | 2146 | 78942 | 0.985529835 | 0.787454739 |
| 14 | | 839 | 32257 | 3418 | 123626 | 0.94075329 | 0.120226571 |
| 15 | | 724 | 28066 | 3715 | 142497 | 0.989475279 | 0.821550105 |
| 16 | | 258 | 10948 | 4427 | 184263 | 0.9808747 | 0.797107331 |
| 17 | | 948 | 43426 | 4577 | 199125 | 0.949737061 | 0.15558942 |
| 18 | | 269 | 7845 | 1104 | 35120 | 1.090799087 | 0.214603491 |
| 19 | | 682 | 32428 | 2416 | 109862 | 0.956345418 | 0.319229262 |
| 20 | | 220 | 9616 | 1761 | 52711 | 0.684810051 | 5.55797E-08 |
| 21 | | 163 | 5075 | 962 | 30596 | 1.021506508 | 0.794907908 |
| 22 | | 192 | 10166 | 1404 | 54772 | 0.736788121 | 5.71499E-05 |
| 1 | T->C | 3532 | 55498 | 28005 | 412937 | 0.938407745 | 0.000536783 |
| 2 | | 4534 | 73864 | 20400 | 294174 | 0.885162102 | 3.32776E-13 |
| 3 | | 3267 | 53097 | 14343 | 202827 | 0.870091486 | 1.8956E-12 |
| 4 | | 2642 | 38618 | 11280 | 153626 | 0.931748406 | 0.001524111 |
| 5 | | 2963 | 48503 | 12538 | 185314 | 0.90290699 | 1.01865E-06 |
| 6 | | 1424 | 23196 | 15139 | 210869 | 0.855091203 | 2.59008E-08 |
| 7 | | 1400 | 27016 | 15982 | 261178 | 0.846861625 | 3.24141E-09 |
| 8 | | 2347 | 38633 | 9568 | 149064 | 0.946468708 | 0.020575816 |
| 9 | | 1149 | 17659 | 10491 | 163627 | 1.014826976 | 0.640223359 |
| 10 | | 762 | 13564 | 12909 | 188829 | 0.821756755 | 1.59693E-07 |
| 11 | | 1765 | 32743 | 14832 | 234408 | 0.851920275 | 2.8881E-10 |
| 12 | | 3407 | 53067 | 13701 | 195205 | 0.914715984 | 5.65445E-06 |
| 13 | | 1277 | 18959 | 5369 | 75719 | 0.949919792 | 0.112490207 |
| 14 | | 1801 | 31295 | 8253 | 118791 | 0.828343465 | 9.31016E-13 |
| 15 | | 1799 | 26991 | 9046 | 137166 | 1.010653022 | 0.688324442 |
| 16 | | 523 | 10683 | 10494 | 178196 | 0.831314493 | 4.26618E-05 |
| 17 | | 2322 | 42052 | 12227 | 191475 | 0.864704513 | 2.56399E-10 |
| 18 | | 431 | 7683 | 2270 | 33954 | 0.839095755 | 0.001012239 |
| 19 | | 1444 | 31666 | 6024 | 106254 | 0.804330081 | 1.51806E-13 |
| 20 | | 521 | 9315 | 3203 | 51269 | 0.895267403 | 0.022886366 |
| 21 | | 286 | 4952 | 1899 | 29659 | 0.902021598 | 0.121852645 |
| 22 | | 487 | 9871 | 3197 | 52979 | 0.817577497 | 4.24341E-05 |
| 1 | T->G | 1853 | 57177 | 15286 | 425656 | 0.902441292 | 3.416E-05 |
| 2 | | 2915 | 75483 | 11920 | 302654 | 0.980527028 | 0.356765659 |
| 3 | | 2021 | 54343 | 8515 | 208655 | 0.911311493 | 0.000207691 |
| 4 | | 1690 | 39570 | 6910 | 157996 | 0.976536991 | 0.401055266 |
| 5 | | 1805 | 49661 | 7190 | 190662 | 0.963822365 | 0.171782222 |

| | | | | | | | |
|---|---|---|---|---|---|---|---|
| 6 | | 797 | 23823 | 9268 | 216740 | 0.78237491 | 1.77895E-11 |
| 7 | | 763 | 27653 | 8609 | 268551 | 0.860709012 | 7.03551E-05 |
| 8 | | 1424 | 39556 | 5540 | 153092 | 0.994810483 | 0.879980172 |
| 9 | | 732 | 18076 | 5479 | 168639 | 1.246423034 | 8.63613E-08 |
| 10 | | 555 | 13771 | 7276 | 194462 | 1.077133573 | 0.100467889 |
| 11 | | 989 | 33519 | 7274 | 241966 | 0.981490921 | 0.596498479 |
| 12 | | 2120 | 54354 | 7237 | 201669 | 1.086888323 | 0.000995358 |
| 13 | | 808 | 19428 | 3149 | 77939 | 1.029355607 | 0.477771943 |
| 14 | | 1073 | 32023 | 4417 | 122627 | 0.930243001 | 0.036973867 |
| 15 | | 996 | 27794 | 4820 | 141392 | 1.051201773 | 0.160631715 |
| 16 | | 272 | 10934 | 4826 | 183864 | 0.947761845 | 0.422567204 |
| 17 | | 1157 | 43217 | 5458 | 198244 | 0.972400701 | 0.397890664 |
| 18 | | 263 | 7851 | 1296 | 34928 | 0.902816501 | 0.142348784 |
| 19 | | 678 | 32432 | 2421 | 109857 | 0.948612642 | 0.233806933 |
| 20 | | 284 | 9552 | 1615 | 52857 | 0.973092241 | 0.697837954 |
| 21 | | 182 | 5056 | 1106 | 30452 | 0.991117209 | 0.935337996 |
| 22 | | 189 | 10169 | 1373 | 54803 | 0.741852138 | 8.74199E-05 |
| 1 | | 8185 | 50845 | 58116 | 382826 | 1.060415684 | 4.47953E-06 |
| 2 | | 11119 | 67279 | 43093 | 271481 | 1.041163449 | 0.000452562 |
| 3 | | 7809 | 48555 | 28840 | 188330 | 1.050233131 | 0.000381138 |
| 4 | | 5813 | 35447 | 22296 | 142610 | 1.048923612 | 0.002701777 |
| 5 | | 7179 | 44287 | 27056 | 170796 | 1.023297225 | 0.108929101 |
| 6 | | 3327 | 21293 | 30351 | 195657 | 1.00725245 | 0.715821001 |
| 7 | | 4283 | 24133 | 39924 | 237236 | 1.054589171 | 0.00239026 |
| 8 | | 5967 | 35013 | 22031 | 136601 | 1.056687043 | 0.000503584 |
| 9 | | 2515 | 16293 | 22384 | 151734 | 1.046362439 | 0.045160221 |
| 10 | | 2217 | 12109 | 27346 | 174392 | 1.167589452 | 2.07751E-10 |
| 11 | C->A | 5109 | 29399 | 34426 | 214814 | 1.084374664 | 7.36007E-07 |
| 12 | | 7935 | 48539 | 28050 | 180856 | 1.054037742 | 0.000131497 |
| 13 | | 2893 | 17343 | 11093 | 69995 | 1.052548737 | 0.023405308 |
| 14 | | 4532 | 28564 | 16608 | 110436 | 1.055028542 | 0.00304965 |
| 15 | | 3897 | 24893 | 19195 | 127017 | 1.03592163 | 0.061899809 |
| 16 | | 1550 | 9656 | 23477 | 165213 | 1.129629586 | 2.03358E-05 |
| 17 | | 5578 | 38796 | 23930 | 179772 | 1.080117272 | 1.41522E-06 |
| 18 | | 1208 | 6906 | 4893 | 31331 | 1.120055134 | 0.00124955 |
| 19 | | 4121 | 28989 | 13566 | 98712 | 1.034397635 | 0.075213545 |
| 20 | | 1419 | 8417 | 7021 | 47451 | 1.139387537 | 4.03538E-05 |
| 21 | | 817 | 4421 | 4439 | 27119 | 1.128989929 | 0.003730465 |
| 22 | | 1381 | 8977 | 6779 | 49397 | 1.120978785 | 0.00035664 |
| 1 | | 12158 | 46872 | 91118 | 349824 | 0.995850342 | 0.704759043 |
| 2 | | 15892 | 62506 | 62092 | 252482 | 1.03383593 | 0.00084401 |
| 3 | | 11564 | 44800 | 43818 | 173352 | 1.02118958 | 0.073759458 |
| 4 | | 7958 | 33302 | 31332 | 133574 | 1.018749555 | 0.185350604 |
| 5 | | 10697 | 40769 | 40175 | 157677 | 1.029779874 | 0.016652028 |
| 6 | | 5314 | 19306 | 44475 | 181533 | 1.123489134 | 1.79613E-12 |
| 7 | | 5876 | 22540 | 55984 | 221176 | 1.029916343 | 0.056551141 |
| 8 | | 8195 | 32785 | 31985 | 126647 | 0.989742733 | 0.459686957 |
| 9 | C->T | 3875 | 14933 | 38940 | 135178 | 0.900813137 | 2.69043E-08 |
| 10 | | 2634 | 11692 | 40861 | 160877 | 0.886976129 | 5.4763E-08 |
| 11 | | 7429 | 27079 | 52666 | 196574 | 1.023984721 | 0.090270319 |
| 12 | | 11303 | 45171 | 42670 | 166236 | 0.974847001 | 0.031523473 |
| 13 | | 3945 | 16291 | 16063 | 65025 | 0.980286373 | 0.318839486 |
| 14 | | 7146 | 25950 | 26224 | 100820 | 1.058701203 | 0.000159312 |
| 15 | | 6064 | 22726 | 31301 | 114911 | 0.979579261 | 0.194252838 |
| 16 | | 2586 | 8620 | 44267 | 144423 | 0.978762961 | 0.358562783 |
| 17 | | 10680 | 33694 | 48393 | 155309 | 1.017261852 | 0.162723512 |

| | | | | | | | |
|---|---|---|---|---|---|---|---|
| 18 | | 1643 | 6471 | 7251 | 28973 | 1.014522597 | 0.634533129 |
| 19 | | 8419 | 24691 | 27549 | 84729 | 1.048692282 | 0.001001852 |
| 20 | | 2129 | 7707 | 10761 | 43711 | 1.122091864 | 1.95481E-05 |
| 21 | | 1010 | 4228 | 6098 | 25460 | 0.997372465 | 0.95479815 |
| 22 | | 2576 | 7782 | 13258 | 42918 | 1.071559012 | 0.005523702 |
| 1 | | 2176 | 56854 | 16097 | 424845 | 1.010144336 | 0.665673597 |
| 2 | | 2673 | 75725 | 11004 | 303570 | 0.973795909 | 0.230962629 |
| 3 | | 1882 | 54482 | 7147 | 210023 | 1.015101928 | 0.569444281 |
| 4 | | 1385 | 39875 | 5946 | 158960 | 0.928564394 | 0.014782948 |
| 5 | | 1749 | 49717 | 6559 | 191293 | 1.025997593 | 0.348572801 |
| 6 | | 893 | 23727 | 7843 | 218165 | 1.046915168 | 0.200342495 |
| 7 | | 1142 | 27274 | 10468 | 266692 | 1.066752187 | 0.043364328 |
| 8 | | 1433 | 39547 | 5712 | 152920 | 0.97008265 | 0.317695167 |
| 9 | | 655 | 18153 | 5879 | 168239 | 1.032561934 | 0.444979337 |
| 10 | | 580 | 13746 | 7131 | 194607 | 1.151488686 | 0.001647809 |
| 11 | | 1124 | 33384 | 8882 | 240358 | 0.911120362 | 0.003593249 |
| 12 | C->G | 2027 | 54447 | 7212 | 201694 | 1.041158992 | 0.117492837 |
| 13 | | 667 | 19569 | 2724 | 78364 | 0.980543112 | 0.678080335 |
| 14 | | 1157 | 31939 | 4602 | 122442 | 0.963819803 | 0.27409106 |
| 15 | | 915 | 27875 | 5029 | 141183 | 0.921524717 | 0.024925236 |
| 16 | | 414 | 10792 | 6854 | 181836 | 1.017733743 | 0.735600132 |
| 17 | | 1502 | 42872 | 7266 | 196436 | 0.94715679 | 0.061117225 |
| 18 | | 243 | 7871 | 1298 | 34926 | 0.830712066 | 0.008878243 |
| 19 | | 1211 | 31899 | 4163 | 108115 | 0.985931214 | 0.678657562 |
| 20 | | 345 | 9491 | 2875 | 51597 | 0.652369613 | 1.30455E-14 |
| 21 | | 161 | 5077 | 1275 | 30283 | 0.753194993 | 0.000688637 |
| 22 | | 354 | 10004 | 2077 | 54099 | 0.921684576 | 0.171308575 |

(b) Contingency table used in Fisher's exact test to assess the association between Mutect2-identified mutations and genomic regions, including well-positioned nucleosomal and non-nucleosomal regions. An odds ratio greater than 1 indicates enrichment of a specific mutation in nucleosomal DNA, whereas an odds ratio less than 1 indicates depletion.

Odds Ratio = (a*d)/(c*b)

| | Specific single nucleotide mutation | Other mutations |
|---|---|---|
| All nucleosomal regions | a | b |
| Inter-ncleosomal regions | c | d |

**Table S9.** (a) Quality control-passed single nucleotide mutations in linker DNA associated with nucleosomal and non-nucleosomal regions.

| Chromosome # | Single nucleotide mutation | Elements of contingency table | | | | Odds ratio | P-value |
|---|---|---|---|---|---|---|---|
| | | a | b | c | d | | |
| 1 | T->A | 698 | 22774 | 12762 | 463738 | 1.113704695 | 0.007234754 |
| 2 | | 940 | 30656 | 9904 | 351472 | 1.088159273 | 0.015617309 |
| 3 | | 709 | 21225 | 6852 | 244748 | 1.193164528 | 1.57457E-05 |
| 4 | | 580 | 16844 | 5251 | 183491 | 1.203249002 | 4.85557E-05 |
| 5 | | 580 | 19174 | 6168 | 223396 | 1.095585556 | 0.039741421 |
| 6 | | 273 | 9969 | 6210 | 234176 | 1.032670651 | 0.611026358 |
| 7 | | 366 | 10754 | 7991 | 286465 | 1.220060847 | 0.000375041 |
| 8 | | 525 | 15513 | 5079 | 178495 | 1.189354591 | 0.000271462 |
| 9 | | 138 | 7024 | 4226 | 181538 | 0.843980937 | 0.051860222 |
| 10 | | 218 | 5638 | 5543 | 204665 | 1.427677527 | 1.40603E-06 |
| 11 | | 362 | 13482 | 7016 | 262888 | 1.006086751 | 0.912778657 |
| 12 | | 662 | 21216 | 6366 | 237136 | 1.162319003 | 0.00036789 |
| 13 | | 278 | 8698 | 2396 | 89952 | 1.199912017 | 0.005760121 |
| 14 | | 462 | 12862 | 3795 | 143021 | 1.353697106 | 5.48615E-09 |
| 15 | | 350 | 11074 | 4089 | 159489 | 1.23275607 | 0.000313542 |
| 16 | | 100 | 4286 | 4585 | 190925 | 0.971563728 | 0.839953146 |
| 17 | | 468 | 16964 | 5057 | 225587 | 1.230661755 | 3.64343E-05 |
| 18 | | 108 | 3170 | 1265 | 39795 | 1.071772172 | 0.495662805 |
| 19 | | 329 | 12209 | 2769 | 130081 | 1.265921324 | 0.000101988 |
| 20 | | 101 | 3723 | 1880 | 58604 | 0.845663815 | 0.111135684 |
| 21 | | 57 | 1883 | 1068 | 33788 | 0.957669748 | 0.838802711 |
| 22 | | 127 | 3997 | 1469 | 60941 | 1.318127295 | 0.004476332 |
| 1 | T->C | 1699 | 21773 | 29838 | 446662 | 1.168111778 | 3.90602E-09 |
| 2 | | 2291 | 29305 | 22643 | 338733 | 1.169517987 | 1.33212E-11 |
| 3 | | 1616 | 20318 | 15994 | 235606 | 1.171627765 | 9.48558E-09 |
| 4 | | 1258 | 16166 | 12664 | 176078 | 1.081962631 | 0.011073807 |
| 5 | | 1323 | 18431 | 14178 | 215386 | 1.090469301 | 0.003892528 |
| 6 | | 675 | 9567 | 15888 | 224498 | 0.996945102 | 0.967602425 |
| 7 | | 614 | 10506 | 16768 | 277688 | 0.967847253 | 0.452790597 |
| 8 | | 1013 | 15025 | 10902 | 172672 | 1.067851117 | 0.05373583 |
| 9 | | 518 | 6644 | 11122 | 174642 | 1.224238242 | 2.37424E-05 |
| 10 | | 422 | 5434 | 13249 | 196959 | 1.154477697 | 0.00596964 |
| 11 | | 944 | 12900 | 15653 | 254251 | 1.188631864 | 1.13096E-06 |
| 12 | | 1508 | 20370 | 15600 | 227902 | 1.081518246 | 0.005305456 |
| 13 | | 605 | 8371 | 6041 | 86307 | 1.032559812 | 0.461255196 |
| 14 | | 883 | 12441 | 9171 | 137645 | 1.065244156 | 0.086164367 |
| 15 | | 781 | 10643 | 10064 | 153514 | 1.119345948 | 0.003844398 |
| 16 | | 281 | 4105 | 10736 | 184774 | 1.178125391 | 0.009956014 |
| 17 | | 1079 | 16353 | 13470 | 217174 | 1.063810427 | 0.058909576 |
| 18 | | 167 | 3111 | 2534 | 38526 | 0.81613832 | 0.012244894 |
| 19 | | 790 | 11748 | 6678 | 126172 | 1.270514793 | 2.03056E-09 |
| 20 | | 310 | 3514 | 3414 | 57070 | 1.474702079 | 1.80189E-09 |
| 21 | | 134 | 1806 | 2051 | 32805 | 1.186755995 | 0.067669015 |
| 22 | | 251 | 3873 | 3433 | 58977 | 1.113358678 | 0.113620277 |
| 1 | T->G | 832 | 22640 | 16307 | 460193 | 1.03708139 | 0.312228921 |
| 2 | | 1306 | 30290 | 13529 | 347847 | 1.108578544 | 0.000564954 |
| 3 | | 949 | 20985 | 9587 | 242013 | 1.14159802 | 0.000176147 |
| 4 | | 844 | 16580 | 7756 | 180986 | 1.187859572 | 5.70586E-06 |

| | | | | | | | |
|---|---|---|---|---|---|---|---|
| 5 | | 842 | 18912 | 8153 | 221411 | 1.209083747 | 5.3492E-07 |
| 6 | | 329 | 9913 | 9736 | 230650 | 0.78625548 | 1.39118E-05 |
| 7 | | 413 | 10707 | 8959 | 285497 | 1.229204839 | 8.67908E-05 |
| 8 | | 675 | 15363 | 6289 | 177285 | 1.238563106 | 4.37778E-07 |
| 9 | | 282 | 6880 | 5929 | 179835 | 1.243235604 | 0.000640185 |
| 10 | | 205 | 5651 | 7626 | 202582 | 0.963679471 | 0.644885936 |
| 11 | | 485 | 13359 | 7778 | 262126 | 1.223516837 | 3.73021E-05 |
| 12 | | 965 | 20913 | 8392 | 235110 | 1.292756119 | 7.76966E-13 |
| 13 | | 434 | 8542 | 3523 | 88825 | 1.281010635 | 3.69895E-06 |
| 14 | | 527 | 12797 | 4963 | 141853 | 1.177054833 | 0.000598422 |
| 15 | | 419 | 11005 | 5397 | 158181 | 1.115901534 | 0.035215608 |
| 16 | | 146 | 4240 | 4952 | 190558 | 1.325053914 | 0.001376707 |
| 17 | | 602 | 16830 | 6013 | 224631 | 1.336259673 | 1.34123E-10 |
| 18 | | 99 | 3179 | 1460 | 39600 | 0.844669858 | 0.114747254 |
| 19 | | 248 | 12290 | 2851 | 129999 | 0.92011602 | 0.219125564 |
| 20 | | 99 | 3725 | 1800 | 58684 | 0.866475168 | 0.183400353 |
| 21 | | 60 | 1880 | 1228 | 33628 | 0.87396909 | 0.341197006 |
| 22 | | 100 | 4024 | 1462 | 60948 | 1.035986543 | 0.710252541 |
| 1 | C->A | 3275 | 20197 | 63026 | 413474 | 1.063782641 | 0.001452903 |
| 2 | | 4473 | 27123 | 49739 | 311637 | 1.033268381 | 0.052456988 |
| 3 | | 2997 | 18937 | 33652 | 217948 | 1.024985146 | 0.230615721 |
| 4 | | 2422 | 15002 | 25687 | 163055 | 1.024815565 | 0.283320111 |
| 5 | | 2695 | 17059 | 31540 | 198024 | 0.99188504 | 0.714190642 |
| 6 | | 1333 | 8909 | 32345 | 208041 | 0.962371975 | 0.20335522 |
| 7 | | 1821 | 9299 | 42386 | 252070 | 1.164588308 | 9.37455E-09 |
| 8 | | 2325 | 13713 | 25673 | 157901 | 1.042794521 | 0.075332683 |
| 9 | | 907 | 6255 | 23992 | 161772 | 0.977725349 | 0.541504702 |
| 10 | | 788 | 5068 | 28775 | 181433 | 0.980371236 | 0.629871867 |
| 11 | | 2002 | 11842 | 37533 | 232371 | 1.04666491 | 0.066211664 |
| 12 | | 3183 | 18695 | 32802 | 210700 | 1.093642504 | 9.75303E-06 |
| 13 | | 1377 | 7599 | 12609 | 79739 | 1.14595519 | 1.29704E-05 |
| 14 | | 1731 | 11593 | 19409 | 127407 | 0.980147331 | 0.462318385 |
| 15 | | 1617 | 9807 | 21475 | 142103 | 1.091048154 | 0.00191673 |
| 16 | | 674 | 3712 | 24353 | 171157 | 1.276127671 | 2.28506E-08 |
| 17 | | 2285 | 15147 | 27223 | 203421 | 1.127247759 | 3.93163E-07 |
| 18 | | 517 | 2761 | 5584 | 35476 | 1.189634014 | 0.000613244 |
| 19 | | 1585 | 10953 | 16102 | 116748 | 1.049218177 | 0.089019253 |
| 20 | | 523 | 3301 | 7917 | 52567 | 1.051982977 | 0.299695552 |
| 21 | | 319 | 1621 | 4937 | 29919 | 1.192591239 | 0.006237453 |
| 22 | | 508 | 3616 | 7652 | 54758 | 1.005328296 | 0.902492381 |
| 1 | C->T | 4305 | 19167 | 98971 | 377529 | 0.856764321 | 9.47091E-20 |
| 2 | | 5755 | 25841 | 72229 | 289147 | 0.891544655 | 2.12042E-14 |
| 3 | | 3961 | 17973 | 51421 | 200179 | 0.857950566 | 1.75399E-17 |
| 4 | | 2976 | 14448 | 36314 | 152428 | 0.864601244 | 2.10148E-12 |
| 5 | | 3749 | 16005 | 47123 | 182441 | 0.906878853 | 1.82054E-07 |
| 6 | | 2120 | 8122 | 47669 | 192717 | 1.055253645 | 0.031594137 |
| 7 | | 1869 | 9251 | 59991 | 234465 | 0.789609821 | 7.0745E-21 |
| 8 | | 2895 | 13143 | 37285 | 146289 | 0.86423447 | 4.43824E-12 |
| 9 | | 1480 | 5682 | 41335 | 144429 | 0.910116417 | 0.001433073 |
| 10 | | 1048 | 4808 | 42447 | 167761 | 0.86147133 | 1.28312E-05 |
| 11 | | 2693 | 11151 | 57402 | 212502 | 0.894043263 | 2.7332E-07 |
| 12 | | 3863 | 18015 | 50110 | 193392 | 0.827569628 | 1.6495E-25 |
| 13 | | 1552 | 7424 | 18456 | 73892 | 0.836977135 | 5.77973E-10 |
| 14 | | 2559 | 10765 | 30811 | 116005 | 0.895008513 | 1.06365E-06 |
| 15 | | 2155 | 9269 | 35210 | 128368 | 0.847627703 | 1.10445E-11 |
| 16 | | 863 | 3523 | 45990 | 149520 | 0.796405098 | 1.51981E-09 |

| | | | | | | | |
|---|---|---|---|---|---|---|---|
| 17 | | 3715 | 13717 | 55358 | 175286 | 0.857563955 | 4.41766E-16 |
| 18 | | 616 | 2662 | 8278 | 32782 | 0.916394945 | 0.059985601 |
| 19 | | 2795 | 9743 | 33173 | 99677 | 0.861984228 | 2.03704E-11 |
| 20 | | 761 | 3063 | 12129 | 48355 | 0.990499023 | 0.835033805 |
| 21 | | 318 | 1622 | 6790 | 28066 | 0.810376833 | 0.000676974 |
| 22 | | 922 | 3202 | 14912 | 47498 | 0.91716827 | 0.024673456 |
| 1 | | 927 | 22545 | 17346 | 459154 | 1.088399979 | 0.014646137 |
| 2 | | 1033 | 30563 | 12644 | 348732 | 0.932205484 | 0.033272004 |
| 3 | | 735 | 21199 | 8294 | 243306 | 1.017093203 | 0.664655898 |
| 4 | | 632 | 16792 | 6699 | 182043 | 1.022771597 | 0.593022663 |
| 5 | | 688 | 19066 | 7620 | 221944 | 1.051035263 | 0.222894958 |
| 6 | | 391 | 9851 | 8345 | 232041 | 1.103658788 | 0.065273384 |
| 7 | | 477 | 10643 | 11133 | 283323 | 1.14057524 | 0.006840801 |
| 8 | | 586 | 15452 | 6559 | 177015 | 1.023494127 | 0.594802433 |
| 9 | | 256 | 6906 | 6278 | 179486 | 1.059796935 | 0.368725008 |
| 10 | | 247 | 5609 | 7464 | 202744 | 1.196156193 | 0.008178979 |
| 11 | C->G | 436 | 13408 | 9570 | 260334 | 0.88458881 | 0.013106958 |
| 12 | | 758 | 21120 | 8481 | 235021 | 0.994568954 | 0.908039057 |
| 13 | | 242 | 8734 | 3149 | 89199 | 0.784855134 | 0.000251513 |
| 14 | | 500 | 12824 | 5259 | 141557 | 1.049481227 | 0.307540175 |
| 15 | | 390 | 11034 | 5554 | 158024 | 1.005654503 | 0.914899658 |
| 16 | | 129 | 4257 | 7139 | 188371 | 0.799581471 | 0.012760436 |
| 17 | | 567 | 16865 | 8201 | 222443 | 0.911903001 | 0.037083373 |
| 18 | | 132 | 3146 | 1409 | 39651 | 1.180751115 | 0.074496971 |
| 19 | | 522 | 12016 | 4852 | 127998 | 1.146022053 | 0.004393248 |
| 20 | | 118 | 3706 | 3102 | 57382 | 0.58899347 | 2.75197E-09 |
| 21 | | 82 | 1858 | 1354 | 33502 | 1.091993901 | 0.433734098 |
| 22 | | 154 | 3970 | 2277 | 60133 | 1.024424731 | 0.764218649 |

(b) Contingency table used in Fisher's exact test to assess the association between mutations and genomic regions, including linker DNA associated with nucleosomal (well-positioned and non-well-positioned) and flanking DNA associated with non-nucleosomal regions. An odds ratio greater than 1 indicates that a specific mutation is enriched in linker DNA associated with nucleosomal DNA, whereas an odds ratio less than 1 indicates its depletion.

Odds Ratio = (a*d)/(c*b)

| | Specific single nucleotide mutation | Other mutations |
|---|---|---|
| Linker DNA associated with all nucleosomal regions | a | b |
| Flanking DNA associated with inter-ncleosomal regions | c | d |

**Table S10.** (a) Mutect2 quality-passed single nucleotide mutations in linker DNA associated with nucleosomal and non-nucleosomal regions.

| Chromosome # | Single nucleotide mutation | Elements of contingency table | | | | Odds ratio | P-value |
|---|---|---|---|---|---|---|---|
| | | a | b | c | d | | |
| 1 | T->A | 648 | 22774 | 12812 | 463738 | 1.02989143 | 0.469255032 |
| 2 | | 784 | 30656 | 10060 | 351472 | 0.89349747 | 0.002562521 |
| 3 | | 624 | 21225 | 6937 | 244748 | 1.03725216 | 0.389535038 |
| 4 | | 442 | 16844 | 5389 | 183491 | 0.8934775 | 0.024244885 |
| 5 | | 513 | 19174 | 6235 | 223396 | 0.95861358 | 0.372226516 |
| 6 | | 230 | 9969 | 6253 | 234176 | 0.864032731 | 0.030333106 |
| 7 | | 313 | 10754 | 8044 | 286465 | 1.036510752 | 0.533009684 |
| 8 | | 461 | 15513 | 5143 | 178495 | 1.031370401 | 0.532432537 |
| 9 | | 116 | 7024 | 4248 | 181538 | 0.705759162 | 0.000134427 |
| 10 | | 179 | 5638 | 5582 | 204665 | 1.164076996 | 0.052357886 |
| 11 | | 302 | 13482 | 7076 | 262888 | 0.832215036 | 0.001750778 |
| 12 | | 565 | 21216 | 6463 | 237136 | 0.977120838 | 0.627997064 |
| 13 | | 234 | 8698 | 2440 | 89952 | 0.991784808 | 0.94488094 |
| 14 | | 403 | 12862 | 3854 | 143021 | 1.162745429 | 0.005253945 |
| 15 | | 272 | 11074 | 4167 | 159489 | 0.940094733 | 0.338656239 |
| 16 | | 84 | 4286 | 4601 | 190925 | 0.813275493 | 0.06845952 |
| 17 | | 401 | 16964 | 5124 | 225587 | 1.040689221 | 0.43942346 |
| 18 | | 93 | 3170 | 1280 | 39795 | 0.912099517 | 0.430910054 |
| 19 | | 268 | 12209 | 2830 | 130081 | 1.00897901 | 0.871266696 |
| 20 | | 95 | 3723 | 1886 | 58604 | 0.792895842 | 0.029613403 |
| 21 | | 51 | 1883 | 1074 | 33788 | 0.852075465 | 0.308292979 |
| 22 | | 106 | 3997 | 1490 | 60941 | 1.084663498 | 0.429250216 |
| 1 | T->C | 1414 | 21773 | 30123 | 446662 | 0.962968146 | 0.184212062 |
| 2 | | 1891 | 29305 | 23043 | 338733 | 0.948567612 | 0.032169852 |
| 3 | | 1308 | 20318 | 16302 | 235606 | 0.930405449 | 0.014695735 |
| 4 | | 1035 | 16166 | 12887 | 176078 | 0.874764285 | 4.83456E-05 |
| 5 | | 1079 | 18431 | 14422 | 215386 | 0.874308147 | 2.84344E-05 |
| 6 | | 540 | 9567 | 16023 | 224498 | 0.790836362 | 8.38944E-08 |
| 7 | | 501 | 10506 | 16881 | 277688 | 0.784439171 | 7.03508E-08 |
| 8 | | 838 | 15025 | 11077 | 172672 | 0.86941935 | 0.000121281 |
| 9 | | 407 | 6644 | 11233 | 174642 | 0.952396351 | 0.359049488 |
| 10 | | 323 | 5434 | 13348 | 196959 | 0.877086691 | 0.022759811 |
| 11 | | 796 | 12900 | 15801 | 254251 | 0.992890726 | 0.866616233 |
| 12 | | 1256 | 20370 | 15852 | 227902 | 0.886467225 | 5.54686E-05 |
| 13 | | 442 | 8371 | 6204 | 86307 | 0.734546273 | 2.38959E-10 |
| 14 | | 693 | 12441 | 9361 | 137645 | 0.819060797 | 4.80468E-07 |
| 15 | | 617 | 10643 | 10228 | 153514 | 0.870118436 | 0.000986641 |
| 16 | | 232 | 4105 | 10785 | 184774 | 0.9682679 | 0.661921861 |
| 17 | | 877 | 16353 | 13672 | 217174 | 0.851879049 | 5.21107E-06 |
| 18 | | 110 | 3111 | 2591 | 38526 | 0.525749879 | 1.54333E-12 |
| 19 | | 579 | 11748 | 6889 | 126172 | 0.902654244 | 0.021243109 |
| 20 | | 252 | 3514 | 3472 | 57070 | 1.178764206 | 0.015988962 |
| 21 | | 103 | 1806 | 2082 | 32805 | 0.898625619 | 0.320052668 |
| 22 | | 178 | 3873 | 3506 | 58977 | 0.773113525 | 0.000851704 |
| 1 | T->G | 658 | 22640 | 16481 | 460193 | 0.811532506 | 1.075E-07 |
| 2 | | 1100 | 30290 | 13735 | 347847 | 0.919714451 | 0.008297827 |
| 3 | | 813 | 20985 | 9723 | 242013 | 0.964317352 | 0.340113292 |
| 4 | | 658 | 16580 | 7942 | 180986 | 0.904391488 | 0.015182049 |

| | | | | | | | |
|---|---|---|---|---|---|---|---|
| 5 | | 682 | 18912 | 8313 | 221411 | 0.960480005 | 0.32819243 |
| 6 | | 284 | 9913 | 9781 | 230650 | 0.675590344 | 1.16614E-11 |
| 7 | | 339 | 10707 | 9033 | 285497 | 1.000694319 | 0.977579181 |
| 8 | | 552 | 15363 | 6412 | 177285 | 0.993439732 | 0.910359219 |
| 9 | | 238 | 6880 | 5973 | 179835 | 1.041526258 | 0.537971224 |
| 10 | | 157 | 5651 | 7674 | 202582 | 0.733421108 | 8.85683E-05 |
| 11 | | 421 | 13359 | 7842 | 262126 | 1.053395378 | 0.311050573 |
| 12 | | 792 | 20913 | 8565 | 235110 | 1.039567224 | 0.308739539 |
| 13 | | 353 | 8542 | 3604 | 88825 | 1.018510644 | 0.752588681 |
| 14 | | 454 | 12797 | 5036 | 141853 | 0.999310582 | 1 |
| 15 | | 313 | 11005 | 5503 | 158181 | 0.817540158 | 0.000515411 |
| 16 | | 106 | 4240 | 4992 | 190558 | 0.954316907 | 0.697004815 |
| 17 | | 509 | 16830 | 6106 | 224631 | 1.112619217 | 0.024508669 |
| 18 | | 88 | 3179 | 1471 | 39600 | 0.745203108 | 0.007613389 |
| 19 | | 199 | 12290 | 2900 | 129999 | 0.72584386 | 6.53513E-06 |
| 20 | | 89 | 3725 | 1810 | 58684 | 0.774648819 | 0.020307192 |
| 21 | | 43 | 1880 | 1245 | 33628 | 0.617792019 | 0.001390767 |
| 22 | | 85 | 4024 | 1477 | 60948 | 0.87164555 | 0.241767601 |
| 1 | | 2909 | 20197 | 63392 | 413474 | 0.939443374 | 0.002010319 |
| 2 | | 3926 | 27123 | 50286 | 311637 | 0.897045548 | 6.28754E-10 |
| 3 | | 2561 | 18937 | 34088 | 217948 | 0.864668755 | 1.47937E-11 |
| 4 | | 2047 | 15002 | 26062 | 163055 | 0.853679911 | 5.31234E-11 |
| 5 | | 2343 | 17059 | 31892 | 198024 | 0.852814926 | 1.51598E-12 |
| 6 | | 1174 | 8909 | 32504 | 208041 | 0.843434305 | 4.45319E-08 |
| 7 | | 1607 | 9299 | 42600 | 252070 | 1.02256563 | 0.413522299 |
| 8 | | 2023 | 13713 | 25975 | 157901 | 0.896794076 | 9.09368E-06 |
| 9 | | 804 | 6255 | 24095 | 161772 | 0.862988799 | 9.3196E-05 |
| 10 | | 689 | 5068 | 28874 | 181433 | 0.854263686 | 0.000100928 |
| 11 | C->A | 1760 | 11842 | 37775 | 232371 | 0.914250202 | 0.000557486 |
| 12 | | 2842 | 18695 | 33143 | 210700 | 0.966432047 | 0.105320002 |
| 13 | | 1208 | 7599 | 12778 | 79739 | 0.992015347 | 0.820962777 |
| 14 | | 1444 | 11593 | 19696 | 127407 | 0.805724578 | 2.32465E-14 |
| 15 | | 1437 | 9807 | 21655 | 142103 | 0.961536227 | 0.180436668 |
| 16 | | 568 | 3712 | 24459 | 171157 | 1.07077035 | 0.135126722 |
| 17 | | 2014 | 15147 | 27494 | 203421 | 0.983763482 | 0.517181981 |
| 18 | | 452 | 2761 | 5649 | 35476 | 1.028099386 | 0.594925654 |
| 19 | | 1402 | 10953 | 16285 | 116748 | 0.917649036 | 0.003494219 |
| 20 | | 470 | 3301 | 7970 | 52567 | 0.939089976 | 0.223282538 |
| 21 | | 268 | 1621 | 4988 | 29919 | 0.991681949 | 0.919375345 |
| 22 | | 439 | 3616 | 7721 | 54758 | 0.861013822 | 0.003838207 |
| 1 | | 3524 | 19167 | 99752 | 377529 | 0.695841718 | 2.09372E-90 |
| 2 | | 4840 | 25841 | 73144 | 289147 | 0.7404164 | 2.89809E-81 |
| 3 | | 3286 | 17973 | 52096 | 200179 | 0.702523929 | 1.50189E-77 |
| 4 | | 2386 | 14448 | 36904 | 152428 | 0.682109371 | 8.20456E-68 |
| 5 | | 3054 | 16005 | 47818 | 182441 | 0.728021811 | 7.5169E-58 |
| 6 | | 1801 | 8122 | 47988 | 192717 | 0.890508571 | 1.06277E-05 |
| 7 | | 1541 | 9251 | 60319 | 234465 | 0.647497129 | 5.30178E-60 |
| 8 | | 2430 | 13143 | 37750 | 146289 | 0.716483974 | 2.11708E-51 |
| 9 | C->T | 1284 | 5682 | 41531 | 144429 | 0.785861133 | 4.72601E-15 |
| 10 | | 814 | 4808 | 42681 | 167761 | 0.665451435 | 1.05867E-28 |
| 11 | | 2296 | 11151 | 57799 | 212502 | 0.757008516 | 2.38035E-34 |
| 12 | | 3272 | 18015 | 50701 | 193392 | 0.692789039 | 1.87351E-83 |
| 13 | | 1267 | 7424 | 18741 | 73892 | 0.67288882 | 4.49917E-39 |
| 14 | | 2143 | 10765 | 31227 | 116005 | 0.739527932 | 9.62939E-37 |
| 15 | | 1751 | 9269 | 35614 | 128368 | 0.680909329 | 2.40795E-50 |
| 16 | | 766 | 3523 | 46087 | 149520 | 0.70540247 | 3.15977E-19 |

| | | | | | | | |
|---|---|---|---|---|---|---|---|
| 17 | | 3181 | 13717 | 55892 | 175286 | 0.72728078 | 1.10265E-58 |
| 18 | | 496 | 2662 | 8398 | 32782 | 0.727332847 | 8.70259E-11 |
| 19 | | 2359 | 9743 | 33609 | 99677 | 0.718082935 | 2.13287E-46 |
| 20 | | 655 | 3063 | 12235 | 48355 | 0.845145955 | 0.000121934 |
| 21 | | 287 | 1622 | 6821 | 28066 | 0.728053876 | 6.4028E-07 |
| 22 | | 757 | 3202 | 15077 | 47498 | 0.744791892 | 3.29206E-13 |
| 1 | | 820 | 22545 | 17453 | 459154 | 0.956867703 | 0.231592227 |
| 2 | | 861 | 30563 | 12816 | 348732 | 0.766560562 | 1.27574E-14 |
| 3 | | 642 | 21199 | 8387 | 243306 | 0.878548677 | 0.001690574 |
| 4 | | 492 | 16792 | 6839 | 182043 | 0.779909225 | 7.02525E-08 |
| 5 | | 612 | 19066 | 7696 | 221944 | 0.925699829 | 0.071784069 |
| 6 | | 331 | 9851 | 8405 | 232041 | 0.927629786 | 0.194799787 |
| 7 | | 405 | 10643 | 11205 | 283323 | 0.962190206 | 0.477862022 |
| 8 | | 481 | 15452 | 6664 | 177015 | 0.826866616 | 5.18633E-05 |
| 9 | | 214 | 6906 | 6320 | 179486 | 0.88003653 | 0.071396929 |
| 10 | | 211 | 5609 | 7500 | 202744 | 1.016912914 | 0.802019776 |
| 11 | C->G | 379 | 13408 | 9627 | 260334 | 0.764390231 | 1.64457E-07 |
| 12 | | 660 | 21120 | 8579 | 235021 | 0.856091182 | 0.000122861 |
| 13 | | 211 | 8734 | 3180 | 89199 | 0.677644836 | 1.38788E-08 |
| 14 | | 429 | 12824 | 5330 | 141557 | 0.88846009 | 0.020641741 |
| 15 | | 320 | 11034 | 5624 | 158024 | 0.814882024 | 0.000363294 |
| 16 | | 114 | 4257 | 7154 | 188371 | 0.705125318 | 0.000141989 |
| 17 | | 479 | 16865 | 8289 | 222443 | 0.762194432 | 3.85282E-09 |
| 18 | | 116 | 3146 | 1425 | 39651 | 1.025979188 | 0.765904465 |
| 19 | | 452 | 12016 | 4922 | 127998 | 0.978228 | 0.691225229 |
| 20 | | 110 | 3706 | 3110 | 57382 | 0.547649332 | 3.02289E-11 |
| 21 | | 77 | 1858 | 1359 | 33502 | 1.021636247 | 0.856349157 |
| 22 | | 136 | 3970 | 2295 | 60133 | 0.897591193 | 0.246191307 |

(b) Contingency table used in Fisher's exact test to assess the association between Mutect2-identified mutations and genomic regions, including linker DNA associated with nucleosomal (well-positioned and non-well-positioned) and flanking DNA associated with non-nucleosomal regions. An odds ratio greater than 1 indicates that a specific mutation is enriched in linker DNA associated with nucleosomal DNA, whereas an odds ratio less than 1 indicates its depletion.

Odds Ratio = (a*d)/(c*b)

| | Specific single nucleotide mutation | Other mutations |
|---|---|---|
| Linker DNA associated with all nucleosomal regions | a | b |
| Flanking DNA associated with inter-ncleosomal regions | c | d |

**Table S11.** (a) Mutect2 quality-passed single nucleotide mutations in linker DNA associated with well-positioned nucleosomal and non-nucleosomal regions.

| Chromosome # | Single nucleotide mutation | Elements of contingency table | | | | Odds ratio | P-value |
|---|---|---|---|---|---|---|---|
| | | a | b | c | d | | |
| 1 | T->A | 616 | 21602 | 12844 | 464910 | 1.032179766 | 0.445295693 |
| 2 | | 760 | 29357 | 10084 | 352771 | 0.905653268 | 0.008852778 |
| 3 | | 598 | 20257 | 6963 | 245716 | 1.041749013 | 0.344702001 |
| 4 | | 434 | 16120 | 5397 | 184215 | 0.918961389 | 0.09645615 |
| 5 | | 496 | 18213 | 6252 | 224357 | 0.977284124 | 0.639518627 |
| 6 | | 225 | 9475 | 6258 | 234670 | 0.890482346 | 0.096063834 |
| 7 | | 303 | 10226 | 8054 | 286993 | 1.055836129 | 0.361574797 |
| 8 | | 427 | 14717 | 5177 | 179291 | 1.004821478 | 0.918479475 |
| 9 | | 110 | 6646 | 4254 | 181916 | 0.707792181 | 0.000241035 |
| 10 | | 167 | 5434 | 5594 | 204869 | 1.125513277 | 0.141172621 |
| 11 | | 289 | 12833 | 7089 | 263537 | 0.837194314 | 0.002903254 |
| 12 | | 545 | 20208 | 6483 | 238144 | 0.990687747 | 0.857062322 |
| 13 | | 210 | 8292 | 2464 | 90358 | 0.928722372 | 0.322386997 |
| 14 | | 400 | 12373 | 3857 | 143510 | 1.202866705 | 0.00071072 |
| 15 | | 267 | 10649 | 4172 | 159914 | 0.961046986 | 0.550448927 |
| 16 | | 83 | 4175 | 4602 | 191036 | 0.825258896 | 0.090844178 |
| 17 | | 391 | 16105 | 5134 | 226446 | 1.070840572 | 0.19930167 |
| 18 | | 88 | 3065 | 1285 | 39900 | 0.89150126 | 0.336758055 |
| 19 | | 255 | 11596 | 2843 | 130694 | 1.010905977 | 0.868189397 |
| 20 | | 93 | 3537 | 1888 | 58790 | 0.818746316 | 0.067075508 |
| 21 | | 51 | 1819 | 1074 | 33852 | 0.883725787 | 0.447860697 |
| 22 | | 102 | 3795 | 1494 | 61143 | 1.099979364 | 0.358783953 |
| 1 | T->C | 1340 | 20644 | 30197 | 447791 | 0.962548252 | 0.187050428 |
| 2 | | 1843 | 28031 | 23091 | 340007 | 0.968125949 | 0.199149232 |
| 3 | | 1245 | 19394 | 16365 | 236530 | 0.927838058 | 0.013218714 |
| 4 | | 1008 | 15464 | 12914 | 176780 | 0.892300299 | 0.000674733 |
| 5 | | 1044 | 17497 | 14457 | 216320 | 0.892802505 | 0.00053565 |
| 6 | | 502 | 9105 | 16061 | 224960 | 0.772247459 | 1.05319E-08 |
| 7 | | 481 | 9986 | 16901 | 278208 | 0.792885959 | 4.94629E-07 |
| 8 | | 807 | 14238 | 11108 | 173459 | 0.885086086 | 0.000991711 |
| 9 | | 388 | 6281 | 11252 | 175005 | 0.960779362 | 0.463761071 |
| 10 | | 317 | 5232 | 13354 | 197161 | 0.894542888 | 0.057472629 |
| 11 | | 760 | 12282 | 15837 | 254869 | 0.995837778 | 0.939093788 |
| 12 | | 1189 | 19419 | 15919 | 228853 | 0.880229299 | 3.10649E-05 |
| 13 | | 419 | 7968 | 6227 | 86710 | 0.732242645 | 4.78602E-10 |
| 14 | | 673 | 11968 | 9381 | 138118 | 0.827931924 | 2.82965E-06 |
| 15 | | 601 | 10230 | 10244 | 153927 | 0.882762902 | 0.003711486 |
| 16 | | 232 | 3994 | 10785 | 184885 | 0.995775536 | 1 |
| 17 | | 845 | 15524 | 13704 | 218003 | 0.865900908 | 6.04389E-05 |
| 18 | | 103 | 3008 | 2598 | 38629 | 0.509135889 | 4.89131E-13 |
| 19 | | 544 | 11162 | 6924 | 126758 | 0.892226669 | 0.012031573 |
| 20 | | 247 | 3337 | 3477 | 57247 | 1.218677488 | 0.004572082 |
| 21 | | 102 | 1745 | 2083 | 32866 | 0.922279003 | 0.479283513 |
| 22 | | 161 | 3684 | 3523 | 59166 | 0.733948894 | 0.000115294 |
| 1 | T->G | 635 | 21462 | 16504 | 461371 | 0.827112554 | 2.24775E-06 |
| 2 | | 1057 | 29008 | 13778 | 349129 | 0.923329996 | 0.014034113 |
| 3 | | 778 | 20029 | 9758 | 242969 | 0.967186849 | 0.388698944 |
| 4 | | 621 | 15890 | 7979 | 181676 | 0.889849985 | 0.0057736 |

| | | | | | | | |
|---|---|---|---|---|---|---|---|
| 5 | | 655 | 17957 | 8340 | 222366 | 0.972545325 | 0.513264938 |
| 6 | | 271 | 9426 | 9794 | 231137 | 0.67850215 | 5.63082E-11 |
| 7 | | 323 | 10187 | 9049 | 286017 | 1.002184023 | 0.954093538 |
| 8 | | 518 | 14564 | 6446 | 178084 | 0.98261568 | 0.729214217 |
| 9 | | 230 | 6508 | 5981 | 180207 | 1.064824772 | 0.360924893 |
| 10 | | 155 | 5437 | 7676 | 202796 | 0.753176539 | 0.00037733 |
| 11 | | 397 | 12723 | 7866 | 262762 | 1.042340461 | 0.425121669 |
| 12 | | 764 | 19911 | 8593 | 236112 | 1.054322645 | 0.169299961 |
| 13 | | 327 | 8139 | 3630 | 89228 | 0.987577616 | 0.860436487 |
| 14 | | 440 | 12317 | 5050 | 142333 | 1.006843476 | 0.879075729 |
| 15 | | 301 | 10586 | 5515 | 158600 | 0.817696751 | 0.000616184 |
| 16 | | 98 | 4136 | 5000 | 190662 | 0.903523985 | 0.349141917 |
| 17 | | 484 | 15990 | 6131 | 225471 | 1.113156618 | 0.027597256 |
| 18 | | 84 | 3073 | 1475 | 39706 | 0.735835682 | 0.005782311 |
| 19 | | 193 | 11668 | 2906 | 130621 | 0.743495395 | 4.42197E-05 |
| 20 | | 84 | 3542 | 1815 | 58867 | 0.769176494 | 0.020025226 |
| 21 | | 43 | 1816 | 1245 | 33692 | 0.640781629 | 0.003471598 |
| 22 | | 85 | 3818 | 1477 | 61154 | 0.9217802 | 0.513100521 |
| 1 | | 2747 | 19176 | 63554 | 414495 | 0.934279984 | 0.001041862 |
| 2 | | 3763 | 26008 | 50449 | 312752 | 0.896963551 | 1.30146E-09 |
| 3 | | 2448 | 18068 | 34201 | 218817 | 0.866849268 | 8.52061E-11 |
| 4 | | 1938 | 14390 | 26171 | 163667 | 0.842235966 | 3.5312E-12 |
| 5 | | 2204 | 16228 | 32031 | 198855 | 0.843165075 | 1.50815E-13 |
| 6 | | 1133 | 8459 | 32545 | 208491 | 0.858052619 | 1.39959E-06 |
| 7 | | 1518 | 8861 | 42689 | 252508 | 1.013323689 | 0.639465412 |
| 8 | | 1924 | 12988 | 26074 | 158626 | 0.901217258 | 3.41939E-05 |
| 9 | | 749 | 5927 | 24150 | 162100 | 0.848228324 | 2.26198E-05 |
| 10 | | 672 | 4869 | 28891 | 181632 | 0.867679405 | 0.000568323 |
| 11 | C->A | 1681 | 11268 | 37854 | 232945 | 0.918041873 | 0.001276267 |
| 12 | | 2701 | 17821 | 33284 | 211574 | 0.963428074 | 0.083629532 |
| 13 | | 1157 | 7224 | 12829 | 80114 | 1.000164033 | 1 |
| 14 | | 1400 | 11156 | 19740 | 127844 | 0.812742054 | 5.46528E-13 |
| 15 | | 1393 | 9427 | 21699 | 142483 | 0.970288625 | 0.311678052 |
| 16 | | 549 | 3619 | 24478 | 171250 | 1.061300603 | 0.20152665 |
| 17 | | 1920 | 14388 | 27588 | 204180 | 0.987628882 | 0.634432114 |
| 18 | | 441 | 2664 | 5660 | 35573 | 1.040419373 | 0.465782988 |
| 19 | | 1353 | 10380 | 16334 | 117321 | 0.93623236 | 0.028202085 |
| 20 | | 450 | 3134 | 7990 | 52734 | 0.947670708 | 0.308626677 |
| 21 | | 263 | 1566 | 4993 | 29974 | 1.008201009 | 0.89098853 |
| 22 | | 418 | 3437 | 7742 | 54937 | 0.862995483 | 0.005381683 |
| 1 | | 3331 | 18199 | 99945 | 378497 | 0.693151994 | 1.16122E-87 |
| 2 | | 4598 | 24802 | 73386 | 290186 | 0.733070095 | 4.99215E-83 |
| 3 | | 3100 | 17183 | 52282 | 200969 | 0.693489009 | 2.51346E-79 |
| 4 | | 2273 | 13853 | 37017 | 153023 | 0.678283253 | 6.04009E-67 |
| 5 | | 2894 | 15236 | 47978 | 183210 | 0.725328258 | 1.8903E-56 |
| 6 | | 1705 | 7736 | 48084 | 193103 | 0.885108181 | 6.07885E-06 |
| 7 | | 1461 | 8799 | 60399 | 234917 | 0.645805287 | 8.24782E-58 |
| 8 | | 2292 | 12467 | 37888 | 146965 | 0.713123736 | 3.63103E-50 |
| 9 | C->T | 1216 | 5384 | 41599 | 144727 | 0.785769546 | 2.29158E-14 |
| 10 | | 779 | 4638 | 42716 | 167931 | 0.660308685 | 1.31173E-28 |
| 11 | | 2178 | 10640 | 57917 | 213013 | 0.752863597 | 5.01644E-34 |
| 12 | | 3111 | 17174 | 50862 | 194233 | 0.691764286 | 1.87139E-80 |
| 13 | | 1201 | 7070 | 18807 | 74246 | 0.670620971 | 5.63791E-38 |
| 14 | | 2023 | 10395 | 31347 | 116375 | 0.722495421 | 1.36363E-40 |
| 15 | | 1657 | 8938 | 35708 | 128699 | 0.668177434 | 6.48076E-53 |
| 16 | | 745 | 3434 | 46108 | 149609 | 0.703942875 | 5.09788E-19 |

| | | | | | | | |
|---|---|---|---|---|---|---|---|
| 17 | | 3028 | 13042 | 56045 | 175961 | 0.728939061 | 2.44296E-55 |
| 18 | | 476 | 2575 | 8418 | 32869 | 0.721784064 | 6.91938E-11 |
| 19 | | 2193 | 9302 | 33775 | 100118 | 0.698842171 | 4.00717E-51 |
| 20 | | 612 | 2929 | 12278 | 48489 | 0.825178016 | 1.89319E-05 |
| 21 | | 275 | 1573 | 6833 | 28115 | 0.719334083 | 3.66552E-07 |
| 22 | | 715 | 3047 | 15119 | 47653 | 0.739606582 | 3.81294E-13 |
| 1 | C->G | 784 | 21380 | 17489 | 460319 | 0.965166601 | 0.350426506 |
| 2 | | 828 | 29268 | 12849 | 350027 | 0.77067187 | 1.32415E-13 |
| 3 | | 611 | 20239 | 8418 | 244266 | 0.876004343 | 0.001652241 |
| 4 | | 468 | 16089 | 6863 | 182746 | 0.774552183 | 5.60006E-08 |
| 5 | | 581 | 18115 | 7727 | 222895 | 0.925182071 | 0.075172163 |
| 6 | | 308 | 9380 | 8428 | 232512 | 0.905875936 | 0.09555395 |
| 7 | | 382 | 10131 | 11228 | 283835 | 0.953179276 | 0.377485468 |
| 8 | | 455 | 14648 | 6690 | 177819 | 0.825629327 | 7.31709E-05 |
| 9 | | 203 | 6533 | 6331 | 179859 | 0.882761209 | 0.086456434 |
| 10 | | 194 | 5410 | 7517 | 202943 | 0.968130697 | 0.688253497 |
| 11 | | 374 | 12755 | 9632 | 260987 | 0.79449934 | 1.02171E-05 |
| 12 | | 624 | 20131 | 8615 | 236010 | 0.849169455 | 7.96136E-05 |
| 13 | | 205 | 8310 | 3186 | 89623 | 0.693947384 | 1.6568E-07 |
| 14 | | 415 | 12356 | 5344 | 142025 | 0.892623971 | 0.027482854 |
| 15 | | 305 | 10615 | 5639 | 158443 | 0.807329465 | 0.0002541 |
| 16 | | 110 | 4149 | 7158 | 188479 | 0.698104641 | 0.000116477 |
| 17 | | 435 | 16042 | 8333 | 223266 | 0.726527328 | 2.58257E-11 |
| 18 | | 113 | 3039 | 1428 | 39758 | 1.035247202 | 0.723885853 |
| 19 | | 448 | 11397 | 4926 | 128617 | 1.026340422 | 0.593721715 |
| 20 | | 109 | 3519 | 3111 | 57569 | 0.573186437 | 1.39711E-09 |
| 21 | | 76 | 1799 | 1360 | 33561 | 1.04250564 | 0.713383305 |
| 22 | | 132 | 3768 | 2299 | 60335 | 0.919376467 | 0.379010458 |

(b) Contingency table used in Fisher's exact test to assess the association between Mutect2-identified mutations and genomic regions, including linker DNA associated with nucleosomal (well-positioned and non-well-positioned) and flanking DNA associated with non-nucleosomal regions. An odds ratio greater than 1 indicates that a specific mutation is enriched in linker DNA associated with nucleosomal DNA, whereas an odds ratio less than 1 indicates its depletion.

Odds Ratio = (a*d)/(c*b)

| | Specific single nucleotide mutation | Other mutations |
|---|---|---|
| Linker DNA associated with all nucleosomal regions | a | b |
| Flanking DNA associated with inter-ncleosomal regions | c | d |

**Table S12.** (a) Mutational motifs in nucleosomal (well-positioned and non-well-positioned) and non-nucleosomal regions. These analyses are based on mutations that passed quality checks.

| Mutational Motif | Elements of contingency table | | | | P-value | Odds ratio |
|---|---|---|---|---|---|---|
| | a | b | c | d | | |
| **[C>T]**G | 43359 | 14534820 | 29117 | 11419303 | 6.6531E-96 | 1.169937253 |
| G**[C>T]**G | 11667 | 3396106 | 8141 | 2853682 | 4.31504E-38 | 1.204219851 |
| T**[C>A]** | 33594 | 93673661 | 20248 | 63240890 | 1.59048E-37 | 1.120108428 |
| **[C>T]**GG | 12960 | 3964897 | 9186 | 3279111 | 8.70987E-30 | 1.166817056 |
| **[C>A]**T | 35270 | 110452058 | 21347 | 73259575 | 3.29221E-26 | 1.095870389 |
| **[C>T]**GC | 10515 | 3397258 | 7539 | 2854284 | 6.66767E-26 | 1.171828852 |
| **[C>T]**GTT | 1902 | 938832 | 1174 | 825068 | 7.38114E-22 | 1.423784547 |
| AA**[C>T]**G | 2050 | 938684 | 1295 | 824947 | 6.32024E-21 | 1.391203703 |
| T**[C>T]** | 30875 | 93676380 | 19294 | 63241844 | 3.17503E-17 | 1.080336668 |
| TT**[C>G]**C | 312 | 8022137 | 397 | 5572577 | 9.25711E-16 | 0.545921365 |
| TT**[C>T]**G | 6110 | 842088 | 3591 | 580638 | 2.70654E-14 | 1.173204666 |
| **[T>G]**T | 14218 | 155111079 | 8740 | 104484874 | 1.47234E-11 | 1.095816112 |
| C**[C>T]**G | 8468 | 3969389 | 6275 | 3282022 | 4.54063E-11 | 1.115796373 |
| **[C>T]**GAG | 3526 | 1188279 | 2207 | 887949 | 5.62856E-11 | 1.193849482 |
| CG**[C>T]** | 2873 | 3404900 | 2029 | 2859794 | 1.99271E-09 | 1.189279597 |
| **[C>A]**G | 6106 | 14572073 | 4260 | 11444160 | 2.86727E-09 | 1.125666609 |
| T**[C>A]**A | 5274 | 31094242 | 3090 | 20572229 | 7.24342E-08 | 1.129231597 |
| **[C>G]**C | 3826 | 80451957 | 2968 | 54912791 | 1.83781E-07 | 0.879868914 |
| T**[T>G]** | 11709 | 155113588 | 7315 | 104486299 | 4.09909E-07 | 1.078238856 |
| **[C>A]**TG | 7115 | 31738154 | 4416 | 21558565 | 2.40092E-06 | 1.094420013 |
| **[C>A]**GTT | 269 | 940465 | 150 | 826092 | 6.2172E-06 | 1.575240248 |
| ATG**[C>A]** | 738 | 5395600 | 351 | 3390744 | 1.47393E-05 | 1.321309329 |
| **[C>T]**T | 18037 | 110469291 | 11380 | 73269542 | 2.91508E-05 | 1.051245025 |
| C**[C>A]**G | 2280 | 3975577 | 1654 | 3286643 | 5.03524E-05 | 1.139598071 |
| **[C>A]**GG | 1989 | 3975868 | 1432 | 3286865 | 6.69476E-05 | 1.148263803 |
| AA**[C>A]**G | 206 | 940528 | 115 | 826127 | 8.53962E-05 | 1.573419278 |
| **[C>T]**TG | 6297 | 31738972 | 3950 | 21559031 | 8.64473E-05 | 1.082861663 |
| CG**[T>C]**T | 163 | 940571 | 86 | 826156 | 0.000101613 | 1.664790658 |
| ATG**[C>A]**A | 207 | 1943165 | 76 | 1181012 | 0.000135192 | 1.655394028 |
| C**[C>T]** | 19756 | 80436027 | 14059 | 54901700 | 0.000158642 | 0.959135063 |
| **[C>T]**GAA | 2654 | 845544 | 1624 | 582605 | 0.000162697 | 1.126037591 |
| AA**[C>T]** | 3696 | 23154398 | 2282 | 15803032 | 0.000164782 | 1.105409641 |
| TT**[T>G]**T | 3434 | 23832082 | 2149 | 16538594 | 0.00017221 | 1.108920665 |
| **[T>C]**A | 9363 | 104360509 | 5667 | 67270996 | 0.000181742 | 1.065009496 |
| T**[C>T]**GG | 3636 | 891951 | 2415 | 651706 | 0.000288214 | 1.100062758 |
| **[C>T]**GGTA | 422 | 129216 | 202 | 83881 | 0.000349531 | 1.35615206 |
| GC**[C>T]** | 4575 | 18130564 | 3409 | 12472521 | 0.000432086 | 0.923223875 |
| A**[T>A]** | 3604 | 122291547 | 2583 | 80184547 | 0.000571808 | 0.914859953 |
| **[C>A]**GC | 1586 | 3406187 | 1170 | 2860653 | 0.000760534 | 1.13845014 |

**Table S13.** (a) Mutational motifs in nucleosomal and non-nucleosomal regions. These analyses are solely based on Mutect2 quality-passed mutations.

| Mutational Motif | Elements of contingency table | | | | P-value | Odds ratio |
|---|---|---|---|---|---|---|
| | a | b | c | d | | |
| **[C>T]**G | 28046 | 11841372 | 29006 | 14127497 | 5.22799E-65 | 1.153576196 |
| T**[C>T]** | 20376 | 71475703 | 21848 | 85446127 | 6.85088E-29 | 1.114913545 |
| G**[C>T]**G | 7688 | 2769758 | 8140 | 3483851 | 3.55277E-27 | 1.187973401 |
| **[C>T]**GG | 8443 | 3256745 | 8823 | 3991964 | 1.43605E-25 | 1.172960456 |
| **[C>T]**GC | 6887 | 2770559 | 7310 | 3484681 | 6.87647E-24 | 1.18497266 |
| AA**[C>T]**G | 1322 | 751637 | 1303 | 1012667 | 1.52434E-15 | 1.366927708 |
| C**[C>T]**G | 5340 | 3259848 | 5675 | 3995112 | 8.66605E-14 | 1.153206283 |
| CG**[C>T]** | 2072 | 2775374 | 2121 | 3489870 | 3.02986E-11 | 1.22839154 |
| **[C>T]**GTT | 1207 | 751752 | 1269 | 1012701 | 7.87606E-10 | 1.281304332 |
| **[C>T]**GAG | 2298 | 968576 | 2200 | 1108832 | 2.13088E-09 | 1.195802317 |
| G**[C>T]** | 19316 | 52777089 | 20073 | 58226425 | 3.04767E-09 | 1.061645702 |
| AT**[C>T]** | 4085 | 16007332 | 4344 | 19312170 | 7.52028E-09 | 1.134525776 |
| **[T>C]**T | 5645 | 115244204 | 6395 | 144355977 | 3.91305E-08 | 1.105704495 |
| TT**[C>T]**G | 4008 | 674106 | 3957 | 750324 | 9.56892E-08 | 1.127411104 |
| **[C>A]**G | 3922 | 11865496 | 4190 | 14152313 | 7.39771E-07 | 1.116439245 |
| T**[C>T]**C | 3767 | 18930479 | 3941 | 22032233 | 2.96495E-06 | 1.112464339 |
| **[C>A]**GTT | 185 | 752774 | 156 | 1013814 | 1.79521E-05 | 1.597131972 |
| **[T>C]**A | 5801 | 75068858 | 6919 | 96560642 | 2.2691E-05 | 1.078449641 |
| **[C>T]**T | 11938 | 85305935 | 13062 | 98432050 | 2.76824E-05 | 1.054579143 |
| **[C>A]**A | 9977 | 88300508 | 10826 | 101519758 | 3.13292E-05 | 1.059544792 |
| T**[T>G]**TTT | 863 | 7289625 | 1000 | 10190848 | 5.81554E-05 | 1.206468347 |
| TTT**[T>G]**T | 827 | 7289661 | 954 | 10190894 | 5.92151E-05 | 1.211886889 |
| T**[T>C]** | 4782 | 115245067 | 5541 | 144356831 | 8.06344E-05 | 1.08102669 |
| T**[T>G]**TT | 1915 | 17471280 | 2235 | 22899956 | 0.000204222 | 1.123055386 |
| TC**[C>T]** | 3796 | 18930450 | 4062 | 22032112 | 0.000205934 | 1.08763075 |
| **[T>C]**TCC | 378 | 6229217 | 341 | 7365081 | 0.000320956 | 1.310634175 |
| CAT**[C>T]** | 1404 | 4337040 | 1480 | 5225917 | 0.000351871 | 1.143074332 |
| **[C>A]**GC | 1050 | 2776396 | 1133 | 3490858 | 0.000375678 | 1.165225988 |
| **[C>T]**TG | 4337 | 24785508 | 4629 | 28512167 | 0.000405247 | 1.077791224 |
| **[T>C]**C | 4410 | 71491669 | 4904 | 85463071 | 0.000510757 | 1.075006738 |
| CACC**[T>C]** | 90 | 1547114 | 152 | 1654902 | 0.000622275 | 0.633357454 |
| T**[C>T]**GG | 2416 | 733383 | 2422 | 811449 | 0.000631811 | 1.103705437 |
| A**[C>T]**GAG | 434 | 206695 | 378 | 228563 | 0.000726581 | 1.269620384 |
| ATG**[C>A]**A | 138 | 1464264 | 101 | 1659858 | 0.000910227 | 1.548849656 |

(b) Contingency table for Fisher's exact test assessing the association between mutational motifs and DNA regions (well-positioned nucleosomal vs. non-nucleosomal). An odds ratio greater than 1 indicates a mutational motif is enriched in well-positioned nucleosomes, whereas an odds ratio less than 1 indicates it is depleted.

Odds Ratio = (a*d)/(c*b)

| | # Mutations in a specific sequence motif | # Specific sequence motif |
|---|---|---|
| All nucleosomal regions | a | b |
| Inter-nucleosomal regions | c | d |

**Table S14.** (a) Mutational motifs in well-positioned nucleosomal and non-nucleosomal regions. These analyses are based on all mutations in the dataset that passed quality checks.

| Mutational Motif | Elements of contingency table | | | | P-value | Odds ratio |
|---|---|---|---|---|---|---|
| | a | b | c | d | | |
| **[C>T]**G | 10973 | 5430665 | 46079 | 20538204 | 2.58186E-23 | 0.90059967 |
| **[C>A]**T | 8735 | 34609360 | 40725 | 149104165 | 1.51917E-11 | 0.924055429 |
| T**[C>A]** | 8406 | 29959981 | 38347 | 126957320 | 7.20156E-10 | 0.928911212 |
| T**[C>T]**GG | 913 | 345198 | 3925 | 1199634 | 4.05094E-09 | 0.80837265 |
| **[T>C]**G | 2467 | 36844750 | 11651 | 152982201 | 4.22539E-09 | 0.879166716 |
| **[C>T]**CA | 1377 | 9580125 | 6377 | 37761125 | 3.78263E-08 | 0.851120933 |
| CT**[C>T]**G | 709 | 468027 | 3051 | 1610119 | 4.37445E-08 | 0.799449609 |
| **[C>T]**A | 3894 | 36843323 | 17734 | 152976118 | 1.42454E-07 | 0.911704473 |
| **[C>T]**GG | 3404 | 1541791 | 13862 | 5706918 | 5.1914E-07 | 0.908949559 |
| **[C>A]**TG | 1764 | 10255654 | 8416 | 43040807 | 7.24458E-07 | 0.879649984 |
| A**[T>A]** | 817 | 38089890 | 4246 | 164381952 | 7.52257E-07 | 0.830398358 |
| **[C>T]**C | 5057 | 27072685 | 21602 | 107180299 | 9.91823E-07 | 0.926792746 |
| **[C>G]**T | 1417 | 34616678 | 7017 | 149137873 | 1.34287E-06 | 0.870003361 |
| **[C>T]**AG | 1281 | 10256137 | 6186 | 43043037 | 3.73777E-06 | 0.869077101 |
| AG**[C>T]** | 1495 | 7025670 | 7177 | 29619688 | 3.98638E-06 | 0.878195038 |
| A**[T>C]** | 2214 | 38088493 | 10618 | 164375580 | 5.29856E-06 | 0.899867162 |
| C**[C>T]** | 5144 | 27072598 | 21844 | 107180057 | 5.50356E-06 | 0.932293897 |
| T**[C>G]** | 1952 | 29966435 | 9241 | 126986426 | 7.24776E-06 | 0.895123716 |
| TT**[C>T]**G | 1493 | 295671 | 6472 | 1128759 | 8.10861E-06 | 0.880671202 |
| GCA**[T>A]** | 17 | 1632128 | 201 | 7154831 | 1.08102E-05 | 0.370764401 |
| **[C>T]**T | 4448 | 34613647 | 20552 | 149124338 | 2.09491E-05 | 0.932420591 |
| T**[C>T]** | 7721 | 29960666 | 34503 | 126961164 | 2.28109E-05 | 0.948278986 |
| C**[T>C]** | 1906 | 34616189 | 9111 | 149135779 | 3.2499E-05 | 0.901279397 |
| **[C>G]**AG | 400 | 10257018 | 2093 | 43047130 | 3.81802E-05 | 0.802072908 |
| **[T>C]**A | 2188 | 31941602 | 10532 | 139687898 | 3.90681E-05 | 0.9085285 |
| T**[C>G]**A | 575 | 9786955 | 2954 | 41882787 | 4.86109E-05 | 0.833000641 |
| **[T>C]**GA | 678 | 9786852 | 3434 | 41882307 | 4.87575E-05 | 0.8449227 |
| G**[C>T]**G | 3184 | 1339135 | 12644 | 4914474 | 6.41262E-05 | 0.924147414 |
| **[C>T]**GAG | 904 | 467832 | 3594 | 1609576 | 9.05512E-05 | 0.865390097 |
| **[C>G]**A | 1182 | 36846035 | 5546 | 152988306 | 0.000116217 | 0.884922193 |
| C**[C>T]**G | 2179 | 1543016 | 8836 | 5711944 | 0.0001262 | 0.912882821 |
| C**[T>G]** | 1011 | 34617084 | 4956 | 149139934 | 0.000159796 | 0.878867334 |
| **[C>A]**C | 4036 | 27073706 | 17064 | 107184837 | 0.000165394 | 0.936388257 |
| **[C>T]**AGC | 343 | 2403682 | 1716 | 9682475 | 0.000192093 | 0.80516745 |
| **[C>A]**A | 3830 | 36843387 | 16973 | 152976879 | 0.000256651 | 0.936928417 |
| **[T>A]**G | 1222 | 36845995 | 5684 | 152988168 | 0.000292834 | 0.892657158 |
| G**[C>T]** | 7659 | 22392662 | 31730 | 88610852 | 0.000305423 | 0.955175523 |
| G**[C>T]**T | 1303 | 7025862 | 6115 | 29620750 | 0.0003997 | 0.898347554 |
| GG**[C>T]** | 2522 | 6420792 | 10242 | 24176677 | 0.000640695 | 0.927189101 |
| **[T>G]**TGT | 212 | 2755371 | 1147 | 11611046 | 0.000643347 | 0.778867721 |
| **[C>G]**TG | 450 | 10256968 | 2244 | 43046979 | 0.000740899 | 0.841614752 |
| GC**[C>A]** | 1238 | 6422076 | 5181 | 24181738 | 0.000786784 | 0.899744339 |
| CT**[C>A]**C | 317 | 2574146 | 1534 | 10170838 | 0.000865332 | 0.816502397 |
| T**[C>A]**C | 1002 | 7872288 | 4721 | 33092409 | 0.000948523 | 0.892197764 |

(b) Contingency table for Fisher's exact test assessing the association between mutational motifs and DNA regions (well-positioned nucleosomal vs. non-nucleosomal). An odds ratio greater than 1 indicates a mutational motif is enriched in well-positioned nucleosomes, whereas an odds ratio less than 1 indicates it is depleted.

Odds Ratio = (a*d)/(c*b)

| | # Mutations in a specific sequence motif | # Specific sequence motif |
|---|---|---|
| Well-positioned nucleosomal regions | a | b |
| Inter-nucleosomal regions | c | d |

**Table S15.** (a) Mutational motifs in flanking linker DNA associated with nucleosomal (well-positioned and non-well-positioned) and non-nucleosomal regions. These analyses are based on mutations that passed quality checks.

| Mutational Motif | Elements of contingency table | | | | P-value | Odds ratio |
|---|---|---|---|---|---|---|
| | a | b | c | d | | |
| T**[C>T]**G | 14643 | 3493523 | 8387 | 2534072 | 1.16077E-67 | 1.266422923 |
| G**[C>T]**G | 12914 | 3620382 | 8322 | 2818833 | 1.33548E-41 | 1.208225565 |
| T**[C>A]** | 35135 | 96713562 | 19332 | 59165365 | 1.27434E-32 | 1.111842679 |
| GAA**[T>A]** | 248 | 7580433 | 350 | 4718242 | 2.79199E-23 | 0.441031729 |
| **[C>A]**T | 37293 | 114249668 | 20771 | 69277999 | 6.9837E-23 | 1.088705217 |
| T**[C>T]**GA | 4405 | 841279 | 2472 | 584147 | 1.53464E-17 | 1.237312923 |
| AA**[T>A]** | 1391 | 40284577 | 1152 | 23765380 | 3.29211E-17 | 0.712328968 |
| **[T>G]**T | 14797 | 158509795 | 8262 | 98045166 | 7.27364E-14 | 1.107792869 |
| C**[C>T]** | 21611 | 84050174 | 14406 | 51850193 | 6.83375E-13 | 0.925429231 |
| T**[C>T]** | 32921 | 96715776 | 18937 | 59165760 | 1.3677E-11 | 1.06349381 |
| A**[T>A]**G | 952 | 29911665 | 761 | 17479975 | 1.59421E-10 | 0.731059139 |
| T**[T>G]** | 12155 | 158512437 | 6884 | 98046544 | 4.55808E-09 | 1.092151907 |
| C**[T>C]** | 8790 | 114278171 | 5770 | 69293000 | 3.04215E-06 | 0.923717444 |
| **[C>A]**A | 15799 | 118067926 | 9033 | 71653073 | 5.91072E-06 | 1.061452282 |
| C**[T>G]** | 4619 | 114282342 | 2517 | 69296253 | 1.53669E-05 | 1.112744271 |
| C**[C>G]** | 3471 | 84068314 | 2385 | 51862214 | 5.43488E-05 | 0.897811048 |
| **[T>C]**A | 9755 | 106311186 | 5374 | 62700271 | 5.74948E-05 | 1.070582319 |
| T**[C>A]**G | 1176 | 3506990 | 717 | 2541742 | 0.000254563 | 1.188735148 |
| G**[C>A]**G | 1670 | 3631626 | 1136 | 2826019 | 0.000460895 | 1.143963323 |
| **[C>A]**C | 15153 | 84056632 | 9781 | 51854818 | 0.000498901 | 0.955724004 |
| T**[C>T]**GAG | 1284 | 282444 | 754 | 194288 | 0.0005463 | 1.171405617 |
| GG**[C>T]** | 9768 | 19065029 | 6531 | 12087782 | 0.000925418 | 0.948276779 |
| TTG**[C>T]** | 1777 | 6187048 | 940 | 3707639 | 0.001983461 | 1.132852926 |
| GG**[C>G]** | 827 | 19073970 | 613 | 12093700 | 0.003491863 | 0.855387956 |
| C**[C>T]**GCA | 736 | 255897 | 431 | 178033 | 0.00462543 | 1.188053122 |
| TAA**[T>A]** | 197 | 9601591 | 154 | 5521875 | 0.004659669 | 0.735679872 |
| CACCA**[T>C]** | 105 | 653581 | 104 | 441528 | 0.005908241 | 0.682047767 |
| TC**[C>T]** | 6298 | 25089721 | 3995 | 15087848 | 0.00863585 | 0.948019654 |
| AAA**[T>G]**G | 94 | 3950095 | 31 | 2222490 | 0.008902809 | 1.70607624 |
| GAA**[T>C]** | 480 | 7580201 | 359 | 4718233 | 0.009512491 | 0.832233993 |

(b) Contingency table for Fisher's exact test assessing the association between mutational motifs and flanking DNA regions (nucleosomal vs. non-nucleosomal). An odds ratio greater than 1 indicates a mutational motif is enriched in flanking regions of nucleosomes, whereas an odds ratio less than 1 indicates it is depleted.

Odds Ratio = (a*d)/(c*b)

| | # Mutations in a specific sequence motif | # Specific sequence motif |
|---|---|---|
| Flanking DNA of nucleosomal regions | a | b |
| Flanking DNA of inter-nucleosomal regions | c | d |

**Table S16.** (a) Mutational motifs in flanking linker DNA associated with nucleosomal (well-positioned and non-well-positioned) and non-nucleosomal regions. These analyses are solely based on Mutect2 quality-passed mutations.

| Mutational Motif | Elements of contingency table | | | | P-value | Odds ratio |
|---|---|---|---|---|---|---|
| | a | b | c | d | | |
| T**[C>A]** | 7577 | 21648686 | 38795 | 134415003 | 5.49367E-51 | 1.212656324 |
| **[T>G]**T | 3326 | 39866471 | 16015 | 219726409 | 2.88906E-12 | 1.144642235 |
| G**[C>T]**G | 1849 | 628410 | 13979 | 5625199 | 2.50417E-11 | 1.184010652 |
| **[C>A]**A | 3148 | 25959931 | 17655 | 163860335 | 1.68886E-09 | 1.125478768 |
| T**[T>G]** | 2711 | 39867086 | 13160 | 219729264 | 2.93104E-09 | 1.135395154 |
| T**[C>A]**C | 824 | 5124584 | 4666 | 34804696 | 2.54655E-06 | 1.199393541 |
| **[C>G]**A | 1044 | 25962035 | 5684 | 163872306 | 1.61005E-05 | 1.159346522 |
| **[T>C]**T | 2011 | 39867786 | 10029 | 219732395 | 5.22583E-05 | 1.105163187 |
| C**[C>A]** | 3547 | 16296998 | 23890 | 117955208 | 7.30955E-05 | 1.07461908 |
| **[T>A]**T | 984 | 39868813 | 4750 | 219737674 | 0.000186099 | 1.141754432 |
| T**[C>G]** | 1667 | 21654596 | 9364 | 134444434 | 0.000199628 | 1.10526632 |
| **[C>T]**A | 3146 | 25959933 | 18482 | 163859508 | 0.000223031 | 1.074429272 |
| **[T>C]**A | 2323 | 29289223 | 10397 | 142340277 | 0.000371747 | 1.085828213 |
| **[T>C]**G | 2063 | 25383065 | 12352 | 165340841 | 0.000452999 | 1.087922667 |
| T**[T>C]** | 1714 | 39868083 | 8609 | 219733815 | 0.000499397 | 1.097310796 |
| TC**[C>G]** | 173 | 5125235 | 897 | 34808465 | 0.001586794 | 1.309859604 |
| G**[C>G]**G | 87 | 630172 | 532 | 5638646 | 0.001618709 | 1.463266223 |
| **[C>A]**C | 2709 | 16297836 | 18391 | 117960707 | 0.002010547 | 1.066132258 |
| **[T>A]**A | 818 | 29290728 | 3524 | 142347150 | 0.002164376 | 1.128069908 |
| GA**[T>G]**G | 33 | 1363604 | 105 | 8202099 | 0.002202556 | 1.890433398 |
| T**[C>T]**C | 1087 | 5124321 | 6712 | 34802650 | 0.003995728 | 1.099900965 |
| G**[C>A]**G | 250 | 630009 | 1849 | 5637329 | 0.005760371 | 1.209844976 |

(b) Contingency table for Fisher's exact test assessing the association between mutational motifs and flanking DNA regions (nucleosomal vs. non-nucleosomal). An odds ratio greater than 1 indicates a mutational motif is enriched in flanking regions of nucleosomes, whereas an odds ratio less than 1 indicates it is depleted.

Odds Ratio = (a*d)/(c*b)

| | # Mutations in a specific sequence motif | # Specific sequence motif |
|---|---|---|
| Flanking DNA of nucleosomal regions | a | b |
| Flanking DNA of inter-nucleosomal regions | c | d |

**Table S17.** (a) Mutational motifs in flanking linker DNA associated with well-positioned nucleosomal and non-nucleosomal regions. These analyses are solely based on Mutect2 quality-passed mutations.

| Mutational Motif | Elements of contingency table | | | | P-value | Odds ratio |
|---|---|---|---|---|---|---|
| | a | b | c | d | | |
| **[C>A]**T | 29486 | 89067524 | 20850 | 90130669 | 0 | 1.43107705 |
| C**[C>T]** | 18286 | 67752954 | 9925 | 66852952 | 0 | 1.817944221 |
| **[C>T]**T | 15598 | 89081412 | 10115 | 90141514 | 2.2352E-272 | 1.560417401 |
| **[C>T]**C | 16688 | 67754552 | 10968 | 66851651 | 1.379E-244 | 1.501241317 |
| C**[T>C]** | 7373 | 89089637 | 3932 | 90147811 | 2.9132E-242 | 1.897399234 |
| **[T>C]**G | 8987 | 92111659 | 5275 | 93424061 | 4.5482E-226 | 1.727970862 |
| C**[C>A]** | 16564 | 67754676 | 11337 | 66851040 | 3.8715E-201 | 1.441570778 |
| **[C>A]**A | 12651 | 92107995 | 8672 | 93420536 | 7.48E-177 | 1.479621429 |
| **[C>T]**A | 13012 | 92107634 | 8992 | 93420242 | 3.4015E-175 | 1.467685886 |
| **[C>A]**C | 12444 | 67758796 | 8762 | 66853667 | 3.7009E-131 | 1.401252198 |
| **[T>G]**G | 2494 | 92118152 | 1320 | 93427956 | 2.25707E-85 | 1.916258739 |
| **[C>G]**T | 4983 | 89092027 | 3330 | 90148217 | 3.25995E-78 | 1.514136243 |
| **[C>G]**A | 4094 | 92116552 | 2710 | 93426504 | 6.55585E-68 | 1.532184173 |
| **[C>G]**C | 3557 | 67767683 | 2204 | 66860171 | 1.00131E-67 | 1.592271496 |
| C**[C>G]** | 2878 | 67768362 | 1713 | 66860626 | 1.64351E-63 | 1.657589078 |
| **[T>A]**G | 4033 | 92116613 | 2794 | 93426410 | 6.12025E-55 | 1.463974527 |
| C**[T>G]** | 3571 | 89093439 | 2471 | 90149050 | 1.45797E-48 | 1.462286721 |
| C**[T>A]** | 4281 | 89092729 | 3126 | 90148447 | 3.13363E-44 | 1.385709651 |
| CAT**[C>T]** | 1818 | 4533307 | 1220 | 4595664 | 2.25525E-29 | 1.510661587 |
| CAT**[C>A]** | 1264 | 4372505 | 950 | 4558023 | 2.03415E-14 | 1.3869783 |
| CAT**[C>G]** | 455 | 4269180 | 349 | 4499409 | 8.42335E-06 | 1.374032408 |

(b) Contingency table for Fisher's exact test assessing the association between mutational motifs and flanking DNA regions (nucleosomal vs. non-nucleosomal). An odds ratio greater than 1 indicates a mutational motif is enriched in flanking regions of nucleosomes, whereas an odds ratio less than 1 indicates it is depleted.

Odds Ratio = (a*d)/(c*b)

| | # Mutations in a specific sequence motif | # Specific sequence motif |
|---|---|---|
| Flanking DNA of well-positioned nucleosomal regions | a | b |
| Flanking DNA of inter-nucleosomal regions | c | d |

**Table S18.** (a) Mutational motifs across different cancer types within nucleosomal DNA (well-positioned and non-well-positioned) and non-nucleosomal regions. These analyses are based mutations that passed quality filters. CCRCC: Renal cell carcinoma; HNSC: head and neck squamous cell carcinoma; UCEC: Urothelial carcinoma; EA: Endometrioid adenocarcinoma; PRCC: Papillary renal cell carcinoma; SCC: Squamous cell carcinoma; ADE: Adenocarcinoma; IDC: Infiltrating duct carcinoma; GBM: GBMoblastoma; ONC: Oncocytoma.

| Mutational Motif | Cancer type | Elements of contingency table | | | | P-value | Odds ratio |
|---|---|---|---|---|---|---|---|
| | | a | b | c | d | | |
| **[C>T]**G | CCRCC | 230 | 6172356 | 144747 | 45534690 | 0 | 0.011722215 |
| | EA | 3808 | 6172356 | 141169 | 45534690 | 0 | 0.198998141 |
| | PRCC | 6 | 6172356 | 144971 | 45534690 | 0 | 0.000305324 |
| | SCC | 478 | 6172356 | 144499 | 45534690 | 0 | 0.024403632 |
| | ADE | 350 | 6172356 | 144627 | 45534690 | 0 | 0.017852954 |
| | IDC | 282 | 6172356 | 144695 | 45534690 | 0 | 0.01437762 |
| | GBM | 846 | 6172356 | 144131 | 45534690 | 0 | 0.043301644 |
| | ONC | 7 | 6172356 | 144970 | 45534690 | 0 | 0.000356214 |
| G**[C>T]**G | CCRCC | 35 | 724817 | 19771 | 5492871 | 0 | 0.01341561 |
| | EA | 504 | 724817 | 19302 | 5492871 | 0 | 0.197878794 |
| | SCC | 71 | 724817 | 19735 | 5492871 | 0 | 0.027264168 |
| | ADE | 47 | 724817 | 19759 | 5492871 | 0 | 0.018026189 |
| | IDC | 40 | 724817 | 19766 | 5492871 | 0 | 0.015336005 |
| | GBM | 90 | 724817 | 19716 | 5492871 | 0 | 0.034593518 |
| T**[C>A]** | CCRCC | 145 | 37302236 | 106868 | 275497054 | 0 | 0.0100208 |
| | EA | 4236 | 37302236 | 102777 | 275497054 | 0 | 0.30439822 |
| | PRCC | 6 | 37302236 | 107007 | 275497054 | 0 | 0.000414115 |
| | SCC | 494 | 37302236 | 106519 | 275497054 | 0 | 0.034251687 |
| | ADE | 791 | 37302236 | 106222 | 275497054 | 0 | 0.054997646 |
| | IDC | 45 | 37302236 | 106968 | 275497054 | 0 | 0.003106996 |
| | GBM | 35 | 37302236 | 106978 | 275497054 | 0 | 0.002416327 |
| **[C>A]**T | CCRCC | 31 | 10458405 | 24539 | 73492524 | 0 | 0.008877334 |
| **[C>T]**GC | CCRCC | 69 | 724817 | 17983 | 5492871 | 0 | 0.029077561 |
| | EA | 1331 | 724817 | 16721 | 5492871 | 6.89569E-81 | 0.603235425 |
| | SCC | 215 | 724817 | 17837 | 5492871 | 0 | 0.091345609 |
| | ADE | 153 | 724817 | 17899 | 5492871 | 0 | 0.064778918 |
| | IDC | 107 | 724817 | 17945 | 5492871 | 0 | 0.045186775 |
| | GBM | 300 | 724817 | 17752 | 5492871 | 0 | 0.128069288 |
| T**[C>T]** | CCRCC | 191 | 37302236 | 99702 | 275497054 | 0 | 0.014148539 |
| | EA | 2023 | 37302236 | 97870 | 275497054 | 0 | 0.1526611 |
| | SCC | 970 | 37302236 | 98923 | 275497054 | 0 | 0.072419671 |
| | ADE | 678 | 37302236 | 99215 | 275497054 | 0 | 0.050470133 |
| | IDC | 106 | 37302236 | 99787 | 275497054 | 0 | 0.00784538 |
| | GBM | 148 | 37302236 | 99745 | 275497054 | 0 | 0.010958539 |
| | ONC | 6 | 37302236 | 99887 | 275497054 | 0 | 0.000443634 |
| TT**[C>T]**G | CCRCC | 12 | 345852 | 19572 | 2715890 | 0 | 0.004814685 |
| | EA | 1080 | 345852 | 18504 | 2715890 | 3.3756E-172 | 0.458331831 |
| | SCC | 43 | 345852 | 19541 | 2715890 | 0 | 0.017279993 |
| | ADE | 30 | 345852 | 19554 | 2715890 | 0 | 0.012047794 |
| | IDC | 17 | 345852 | 19567 | 2715890 | 0 | 0.006822547 |
| | GBM | 47 | 345852 | 19537 | 2715890 | 0 | 0.018891301 |
| **[T>G]**T | CCRCC | 23 | 30187424 | 22935 | 229342499 | 0 | 0.007618818 |
| | EA | 392 | 30187424 | 22566 | 229342499 | 0 | 0.131974483 |
| | SCC | 33 | 30187424 | 22925 | 229342499 | 0 | 0.010936115 |
| | ADE | 41 | 30187424 | 22917 | 229342499 | 0 | 0.013592038 |
| | GBM | 8 | 30187424 | 22950 | 229342499 | 0 | 0.002648291 |
| **[C>T]**GAG | CCRCC | 17 | 253727 | 5715 | 1813982 | 3.5456E-291 | 0.021266645 |
| | EA | 321 | 253727 | 5411 | 1813982 | 1.63337E-64 | 0.424124917 |
| | SCC | 34 | 253727 | 5698 | 1813982 | 9.6833E-266 | 0.042660188 |
| | ADE | 15 | 253727 | 5717 | 1813982 | 2.1763E-294 | 0.018758122 |
| | IDC | 13 | 253727 | 5719 | 1813982 | 6.8394E-298 | 0.016251354 |

| | | | | | | | |
|---|---|---|---|---|---|---|---|
| | GBM | 29 | 253727 | 5703 | 1813982 | 1.255E-272 | 0.036354729 |
| CG**[C>T]** | EA | 6 | 724817 | 4895 | 5492871 | 3.2765E-250 | 0.009289013 |
| **[C>A]**G | CCRCC | 23 | 6172356 | 20489 | 45534690 | 0 | 0.008281299 |
| | EA | 146 | 6172356 | 20366 | 45534690 | 0 | 0.052885734 |
| | SCC | 212 | 6172356 | 20300 | 45534690 | 0 | 0.077042655 |
| | ADE | 366 | 6172356 | 20146 | 45534690 | 0 | 0.134024339 |
| | IDC | 15 | 6172356 | 20497 | 45534690 | 0 | 0.00539874 |
| | GBM | 19 | 6172356 | 20493 | 45534690 | 0 | 0.006839738 |
| **[C>G]**C | CCRCC | 26 | 16650939 | 6768 | 118586034 | 0 | 0.027359478 |
| | EA | 50 | 16650939 | 6744 | 118586034 | 1.1224E-303 | 0.052801621 |
| | SCC | 64 | 16650939 | 6730 | 118586034 | 1.5401E-286 | 0.06772667 |
| | ADE | 69 | 16650939 | 6725 | 118586034 | 6.4338E-281 | 0.073072105 |
| T**[T>G]** | CCRCC | 90 | 30187424 | 18934 | 229342499 | 0 | 0.036112589 |
| | EA | 1729 | 30187424 | 17295 | 229342499 | 1.18733E-29 | 0.759508979 |
| | SCC | 89 | 30187424 | 18935 | 229342499 | 0 | 0.035709452 |
| | ADE | 42 | 17933979 | 10955 | 134946194 | 0 | 0.028848344 |
| **[C>A]**TG | CCRCC | 30 | 6423100 | 11501 | 46830475 | 0 | 0.019018205 |
| | EA | 646 | 6423100 | 10885 | 46830475 | 7.0062E-122 | 0.432701065 |
| | SCC | 139 | 6423100 | 11392 | 46830475 | 0 | 0.088960805 |
| | ADE | 323 | 6423100 | 11208 | 46830475 | 1.7486E-284 | 0.210115591 |
| | IDC | 6 | 3927710 | 6498 | 28385363 | 0 | 0.006673084 |
| ATG**[C>A]** | EA | 26 | 1064580 | 1063 | 7713540 | 1.07757E-31 | 0.177221136 |
| **[C>T]**T | CCRCC | 40 | 10458405 | 12064 | 73492524 | 0 | 0.023299488 |
| **[C>T]**TG | CCRCC | 71 | 6423100 | 10176 | 46830475 | 0 | 0.050870397 |
| | EA | 638 | 6423100 | 9609 | 46830475 | 8.03665E-87 | 0.484090282 |
| | SCC | 210 | 6423100 | 10037 | 46830475 | 0 | 0.152545447 |
| | ADE | 185 | 6423100 | 10062 | 46830475 | 0 | 0.134051382 |
| | IDC | 32 | 6423100 | 10215 | 46830475 | 0 | 0.022839968 |
| | GBM | 38 | 6423100 | 10209 | 46830475 | 0 | 0.027138402 |
| CG**[T>C]**T | SCC | 6 | 193703 | 243 | 1565145 | 7.2697E-07 | 0.199509329 |
| ATG**[C>A]**A | SCC | 10 | 379199 | 273 | 2742299 | 6.16761E-07 | 0.26490184 |
| | ADE | 13 | 379199 | 270 | 2742299 | 2.31483E-05 | 0.348198752 |
| C**[C>T]** | EA | 21 | 16650939 | 33793 | 118586034 | 0 | 0.004425755 |
| | SCC | 9 | 16650939 | 33805 | 118586034 | 0 | 0.001896079 |
| | ADE | 8 | 16650939 | 33806 | 118586034 | 0 | 0.001685354 |
| | IDC | 6 | 16650939 | 33808 | 118586034 | 0 | 0.00126394 |
| | GBM | 9 | 16650939 | 33805 | 118586034 | 0 | 0.001896079 |
| **[C>T]**GAA | CCRCC | 16 | 345852 | 8535 | 2715890 | 0 | 0.014721035 |
| | EA | 412 | 345852 | 8139 | 2715890 | 2.11037E-98 | 0.397509994 |
| | SCC | 27 | 345852 | 8524 | 2715890 | 0 | 0.024873804 |
| | ADE | 16 | 345852 | 8535 | 2715890 | 0 | 0.014721035 |
| | IDC | 20 | 345852 | 8531 | 2715890 | 0 | 0.018409922 |
| | GBM | 54 | 345852 | 8497 | 2715890 | 0 | 0.049905686 |
| AA**[C>T]** | CCRCC | 47 | 4572439 | 5931 | 34367837 | 6.9174E-248 | 0.059562678 |
| | EA | 399 | 4572439 | 5579 | 34367837 | 3.32909E-39 | 0.537552411 |
| | SCC | 95 | 4572439 | 5883 | 34367837 | 1.5771E-197 | 0.121374942 |
| | ADE | 64 | 4572439 | 5914 | 34367837 | 2.441E-228 | 0.081339769 |
| | IDC | 35 | 4572439 | 5943 | 34367837 | 2.72E-263 | 0.044265624 |
| | GBM | 88 | 4572439 | 5890 | 34367837 | 5.6825E-204 | 0.112297906 |
| TT**[T>G]**T | CCRCC | 8 | 4616078 | 5575 | 35730805 | 8.7167E-277 | 0.01110746 |
| **[T>C]**A | CCRCC | 37 | 19668412 | 14993 | 151926552 | 0 | 0.0190624 |
| T**[C>T]**GG | CCRCC | 16 | 191531 | 6035 | 1345850 | 0 | 0.018629461 |
| A**[T>A]** | CCRCC | 60 | 46933268 | 12529 | 357907384 | 0 | 0.03651949 |
| | EA | 266 | 46933268 | 12323 | 357907384 | 0 | 0.164609559 |
| | SCC | 133 | 46933268 | 12456 | 357907384 | 0 | 0.081425963 |
| | ADE | 126 | 46933268 | 12463 | 357907384 | 0 | 0.077097059 |
| | IDC | 19 | 46933268 | 12570 | 357907384 | 0 | 0.011526785 |
| | GBM | 21 | 34256257 | 8663 | 250934718 | 0 | 0.017757091 |
| **[C>A]**GC | CCRCC | 21 | 724817 | 2735 | 5492871 | 1.2374E-114 | 0.058187941 |
| | EA | 48 | 724817 | 2708 | 5492871 | 4.71834E-87 | 0.13432709 |
| | SCC | 78 | 724817 | 2678 | 5492871 | 1.57904E-64 | 0.220726796 |
| | ADE | 142 | 724817 | 2614 | 5492871 | 5.78181E-32 | 0.411674333 |
| | GBM | 7 | 724817 | 2749 | 5492871 | 1.3583E-134 | 0.019297201 |

(b) Contingency table for Fisher's exact test assessing the association between mutational motifs across different cancer types and DNA regions (nucleosomal vs. non-nucleosomal). An odds ratio greater than 1 indicates that a mutational motif is enriched in a specific cancer type and within nucleosomal regions, whereas an odds ratio less than 1 indicates depletion.

Odds Ratio = (a*d)/(c*b)

| | # Mutational motif in a specific cancer type | #Mutations |
|---|---|---|
| Nucleosomal regions | a | b |
| Inter-nucleosomal regions | c | d |

**Table S19.** (a) Mutational motifs across different cancer types within nucleosomal DNA and non-nucleosomal regions. These analyses are solely based on Mutect2 mutations that passed quality filters. CCRCC: Renal cell carcinoma; HNSC: head and neck squamous cell carcinoma; UCEC: Urothelial carcinoma; EA: Endometrioid adenocarcinoma; PRCC: Papillary renal cell carcinoma; SCC: Squamous cell carcinoma; ADE: Adenocarcinoma; IDC: Infiltrating duct carcinoma; GBM: GBMoblastoma; ONC: Oncocytoma.

| Mutational Motif | Cancer type | Elements of contingency table | | | | P-value | Odds ratio |
|---|---|---|---|---|---|---|---|
| | | a | b | c | d | | |
| **[C>T]**G | CCRCC | 230 | 6172356 | 144747 | 45534690 | 0 | 0.011722215 |
| | EA | 3808 | 6172356 | 141169 | 45534690 | 0 | 0.198998141 |
| | PRCC | 6 | 6172356 | 144971 | 45534690 | 0 | 0.000305324 |
| | SCC | 478 | 6172356 | 144499 | 45534690 | 0 | 0.024403632 |
| | ADE | 350 | 6172356 | 144627 | 45534690 | 0 | 0.017852954 |
| | IDC | 282 | 6172356 | 144695 | 45534690 | 0 | 0.01437762 |
| | GBM | 846 | 6172356 | 144131 | 45534690 | 0 | 0.043301644 |
| | ONC | 7 | 6172356 | 144970 | 45534690 | 0 | 0.000356214 |
| T**[C>T]** | CCRCC | 191 | 37302236 | 99702 | 275497054 | 0 | 0.014148539 |
| | EA | 2023 | 37302236 | 97870 | 275497054 | 0 | 0.1526611 |
| | SCC | 970 | 37302236 | 98923 | 275497054 | 0 | 0.072419671 |
| | ADE | 678 | 37302236 | 99215 | 275497054 | 0 | 0.050470133 |
| | IDC | 106 | 37302236 | 99787 | 275497054 | 0 | 0.00784538 |
| | GBM | 148 | 37302236 | 99745 | 275497054 | 0 | 0.010958539 |
| | ONC | 6 | 37302236 | 99887 | 275497054 | 0 | 0.000443634 |
| G**[C>T]**G | CCRCC | 35 | 724817 | 19771 | 5492871 | 0 | 0.01341561 |
| | EA | 504 | 724817 | 19302 | 5492871 | 0 | 0.197878794 |
| | SCC | 71 | 724817 | 19735 | 5492871 | 0 | 0.027264168 |
| | ADE | 47 | 724817 | 19759 | 5492871 | 0 | 0.018026189 |
| | IDC | 40 | 724817 | 19766 | 5492871 | 0 | 0.015336005 |
| | GBM | 90 | 724817 | 19716 | 5492871 | 0 | 0.034593518 |
| **[C>T]**GC | CCRCC | 69 | 724817 | 17983 | 5492871 | 0 | 0.029077561 |
| | EA | 1331 | 724817 | 16721 | 5492871 | 6.89569E-81 | 0.603235425 |
| | SCC | 215 | 724817 | 17837 | 5492871 | 0 | 0.091345609 |
| | ADE | 153 | 724817 | 17899 | 5492871 | 0 | 0.064778918 |
| | IDC | 107 | 724817 | 17945 | 5492871 | 0 | 0.045186775 |
| | GBM | 300 | 724817 | 17752 | 5492871 | 0 | 0.128069288 |
| CG**[C>T]** | EA | 6 | 724817 | 4895 | 5492871 | 3.2765E-250 | 0.009289013 |
| **[C>T]**GAG | CCRCC | 17 | 253727 | 5715 | 1813982 | 3.5456E-291 | 0.021266645 |
| | EA | 321 | 253727 | 5411 | 1813982 | 1.63337E-64 | 0.424124917 |
| | SCC | 34 | 253727 | 5698 | 1813982 | 9.6833E-266 | 0.042660188 |
| | ADE | 15 | 253727 | 5717 | 1813982 | 2.1763E-294 | 0.018758122 |
| | IDC | 13 | 253727 | 5719 | 1813982 | 6.8394E-298 | 0.016251354 |
| | GBM | 29 | 253727 | 5703 | 1813982 | 1.255E-272 | 0.036354729 |
| G**[C>T]** | CCRCC | 47 | 27428808 | 91493 | 194392176 | 0 | 0.003640674 |
| | EA | 879 | 27428808 | 90661 | 194392176 | 0 | 0.068713207 |
| | SCC | 139 | 27428808 | 91401 | 194392176 | 0 | 0.010777938 |
| | ADE | 118 | 27428808 | 91422 | 194392176 | 0 | 0.009147515 |
| | IDC | 32 | 27428808 | 91508 | 194392176 | 0 | 0.002478351 |
| | GBM | 53 | 27428808 | 91487 | 194392176 | 0 | 0.004105711 |
| **[T>C]**T | CCRCC | 57 | 30187424 | 15028 | 229342499 | 0 | 0.028815898 |
| | EA | 432 | 30187424 | 14653 | 229342499 | 0 | 0.223983323 |
| | SCC | 111 | 30187424 | 14974 | 229342499 | 0 | 0.056317535 |
| | ADE | 82 | 30187424 | 15003 | 229342499 | 0 | 0.041523526 |
| | IDC | 39 | 30187424 | 15046 | 229342499 | 0 | 0.019692554 |
| | GBM | 26 | 30187424 | 15059 | 229342499 | 0 | 0.013117036 |
| TT**[C>T]**G | CCRCC | 12 | 345852 | 19572 | 2715890 | 0 | 0.004814685 |
| | EA | 1080 | 345852 | 18504 | 2715890 | 3.3756E-172 | 0.458331831 |
| | SCC | 43 | 345852 | 19541 | 2715890 | 0 | 0.017279993 |
| | ADE | 30 | 345852 | 19554 | 2715890 | 0 | 0.012047794 |
| | IDC | 17 | 345852 | 19567 | 2715890 | 0 | 0.006822547 |

| | | | | | | | |
|---|---|---|---|---|---|---|---|
| | GBM | 47 | 345852 | 19537 | 2715890 | 0 | 0.018891301 |
| **[C>A]**G | CCRCC | 23 | 6172356 | 20489 | 45534690 | 0 | 0.008281299 |
| | EA | 146 | 6172356 | 20366 | 45534690 | 0 | 0.052885734 |
| | SCC | 212 | 6172356 | 20300 | 45534690 | 0 | 0.077042655 |
| | ADE | 366 | 6172356 | 20146 | 45534690 | 0 | 0.134024339 |
| | IDC | 15 | 6172356 | 20497 | 45534690 | 0 | 0.00539874 |
| | GBM | 19 | 6172356 | 20493 | 45534690 | 0 | 0.006839738 |
| T**[C>T]**C | CCRCC | 43 | 4906109 | 8915 | 36020933 | 0 | 0.035413176 |
| | EA | 242 | 4906109 | 8716 | 36020933 | 4.9379E-225 | 0.203852438 |
| | SCC | 266 | 4906109 | 8692 | 36020933 | 5.8482E-209 | 0.224687899 |
| | ADE | 118 | 4906109 | 8840 | 36020933 | 0 | 0.098004836 |
| | IDC | 19 | 4906109 | 8939 | 36020933 | 0 | 0.01560567 |
| | GBM | 25 | 4906109 | 8933 | 36020933 | 0 | 0.020547569 |
| **[T>C]**A | CCRCC | 138 | 39336824 | 29709 | 303853104 | 0 | 0.035880248 |
| | EA | 1099 | 39336824 | 28748 | 303853104 | 0 | 0.295293876 |
| | SCC | 359 | 39336824 | 29488 | 303853104 | 0 | 0.094040194 |
| | ADE | 231 | 39336824 | 29616 | 303853104 | 0 | 0.060249017 |
| | IDC | 69 | 39336824 | 29778 | 303853104 | 0 | 0.017898554 |
| | GBM | 68 | 39336824 | 29779 | 303853104 | 0 | 0.017638562 |
| **[C>T]**T | CCRCC | 40 | 10458405 | 12064 | 73492524 | 0 | 0.023299488 |
| **[C>A]**A | CCRCC | 174 | 46086368 | 48053 | 334318519 | 0 | 0.026267376 |
| | EA | 640 | 46086368 | 47587 | 334318519 | 0 | 0.097561754 |
| | SCC | 549 | 46086368 | 47678 | 334318519 | 0 | 0.083529959 |
| | ADE | 992 | 46086368 | 47235 | 334318519 | 0 | 0.15234763 |
| | IDC | 74 | 46086368 | 48153 | 334318519 | 0 | 0.011147984 |
| | GBM | 66 | 46086368 | 48161 | 334318519 | 0 | 0.009941145 |
| T**[T>C]** | CCRCC | 93 | 30187424 | 12806 | 229342499 | 0 | 0.05517317 |
| | EA | 894 | 30187424 | 12005 | 229342499 | 4.23156E-71 | 0.565762086 |
| | SCC | 177 | 30187424 | 12722 | 229342499 | 0 | 0.105700334 |
| | ADE | 143 | 30187424 | 12756 | 229342499 | 0 | 0.085168699 |
| | IDC | 59 | 30187424 | 12840 | 229342499 | 0 | 0.034909648 |
| | GBM | 73 | 30187424 | 12826 | 229342499 | 0 | 0.043240441 |
| T**[T>G]**TT | CCRCC | 7 | 4616078 | 4991 | 35730805 | 2.7531E-248 | 0.010856257 |
| | EA | 277 | 4616078 | 4721 | 35730805 | 8.44762E-47 | 0.454166849 |
| | SCC | 10 | 4616078 | 4988 | 35730805 | 1.5931E-242 | 0.015518267 |
| TC**[C>T]** | EA | 9 | 4906109 | 9833 | 36020933 | 0 | 0.006720077 |
| **[T>C]**TCC | CCRCC | 13 | 1610683 | 976 | 11968680 | 8.62426E-37 | 0.098975958 |
| | EA | 60 | 1610683 | 929 | 11968680 | 1.1636E-09 | 0.47992317 |
| | SCC | 21 | 1610683 | 968 | 11968680 | 9.75283E-30 | 0.161205598 |
| | ADE | 10 | 1610683 | 979 | 11968680 | 6.66343E-40 | 0.075902047 |
| **[C>A]**GC | CCRCC | 21 | 724817 | 2735 | 5492871 | 1.2374E-114 | 0.058187941 |
| | EA | 48 | 724817 | 2708 | 5492871 | 4.71834E-87 | 0.13432709 |
| | SCC | 78 | 724817 | 2678 | 5492871 | 1.57904E-64 | 0.220726796 |
| | ADE | 142 | 724817 | 2614 | 5492871 | 5.78181E-32 | 0.411674333 |
| | GBM | 7 | 724817 | 2749 | 5492871 | 1.3583E-134 | 0.019297201 |
| **[C>T]**TG | CCRCC | 71 | 6423100 | 10176 | 46830475 | 0 | 0.050870397 |
| | EA | 638 | 6423100 | 9609 | 46830475 | 8.03665E-87 | 0.484090282 |
| | SCC | 210 | 6423100 | 10037 | 46830475 | 0 | 0.152545447 |
| | ADE | 185 | 6423100 | 10062 | 46830475 | 0 | 0.134051382 |
| | IDC | 32 | 6423100 | 10215 | 46830475 | 0 | 0.022839968 |
| | GBM | 38 | 6423100 | 10209 | 46830475 | 0 | 0.027138402 |
| **[T>C]**C | CCRCC | 88 | 37302236 | 23399 | 275497054 | 0 | 0.027775857 |
| | EA | 540 | 37302236 | 22947 | 275497054 | 0 | 0.173800066 |
| | SCC | 116 | 37302236 | 23371 | 275497054 | 0 | 0.036657495 |
| | ADE | 108 | 37302236 | 23379 | 275497054 | 0 | 0.034117714 |
| | IDC | 49 | 37302236 | 23438 | 275497054 | 0 | 0.015440367 |
| | GBM | 85 | 37302236 | 23402 | 275497054 | 0 | 0.026825514 |
| CACC**[T>C]** | EA | 22 | 401101 | 278 | 2796427 | 0.005094207 | 0.551731306 |
| | SCC | 10 | 401101 | 290 | 2796427 | 4.06205E-08 | 0.240409566 |
| T**[C>T]**GG | CCRCC | 16 | 191531 | 6035 | 1345850 | 0 | 0.018629461 |
| | EA | 567 | 191531 | 5484 | 1345850 | 5.37569E-14 | 0.726512675 |
| | SCC | 55 | 191531 | 5996 | 1345850 | 2.9387E-261 | 0.064455302 |
| | ADE | 45 | 191531 | 6006 | 1345850 | 1.6336E-273 | 0.05264835 |

| | | | | | | | |
|---|---|---|---|---|---|---|---|
| | IDC | 25 | 191531 | 6026 | 1345850 | 3.3888E-301 | 0.029152007 |
| | GBM | 34 | 191531 | 6017 | 1345850 | 3.0317E-288 | 0.039706032 |
| A**[C>T]**GAG | EA | 91 | 53780 | 966 | 379515 | 0.000102427 | 0.664771533 |
| | SCC | 18 | 53780 | 1039 | 379515 | 6.35009E-38 | 0.122254571 |
| | ADE | 8 | 53780 | 1049 | 379515 | 2.3973E-48 | 0.053817392 |
| | IDC | 7 | 53780 | 1050 | 379515 | 1.07995E-49 | 0.04704537 |
| | GBM | 15 | 53780 | 1042 | 379515 | 1.24117E-40 | 0.101585492 |
| ATG**[C>A]**A | SCC | 10 | 379199 | 273 | 2742299 | 6.16761E-07 | 0.26490184 |
| | ADE | 13 | 379199 | 270 | 2742299 | 2.31483E-05 | 0.348198752 |

(b) Contingency table for Fisher's exact test assessing the association between mutational motifs across different cancer types and DNA regions (nucleosomal vs. non-nucleosomal). An odds ratio greater than 1 indicates that a mutational motif is enriched in a specific cancer type and within nucleosomal regions, whereas an odds ratio less than 1 indicates depletion.

Odds Ratio = (a*d)/(c*b)

| | # Mutational motif in a specific cancer type | #Mutations |
|---|---|---|
| Nucleosomal regions | a | b |
| Inter-nucleosomal regions | c | d |

**Table S20.** (a) Mutational motifs across different cancer types within well-positioned nucleosomal DNA and non-nucleosomal regions. These analyses are based solely on mutations that passed Mutect2 quality filters for various cancer types, including clear cell renal cell carcinoma (CCRCC), glioblastoma (GBM), head and neck squamous cell carcinoma (HNSC), lung squamous cell carcinoma (LUSC), lung adenocarcinoma (LUAD), pancreatic ductal adenocarcinoma (PDAC), and uterine corpus endometrial carcinoma (UCEC).

| Mutational Motif | Cancer type | Elements of contingency table | | | | P-value | Odds ratio |
|---|---|---|---|---|---|---|---|
| | | a | b | c | d | | |
| **[C>T]**G | LUSC | 321 | 10816156 | 144656 | 40890890 | 0 | 0.008389232 |
| | LUAD | 525 | 10816156 | 144452 | 40890890 | 0 | 0.013740083 |
| | HNSC | 442 | 10816156 | 144535 | 40890890 | 0 | 0.011561199 |
| | UCEC | 5583 | 10816156 | 139394 | 40890890 | 0 | 0.151417882 |
| | GBM | 1227 | 10816156 | 143750 | 40890890 | 0 | 0.032269358 |
| | PDAC | 478 | 10816156 | 144499 | 40890890 | 0 | 0.01250595 |
| | CCRCC | 418 | 10816156 | 144559 | 40890890 | 0 | 0.010931626 |
| **[C>A]**T | LUSC | 244 | 34596050 | 56373 | 149041908 | 0 | 0.018646639 |
| | LUAD | 622 | 34596050 | 55995 | 149041908 | 0 | 0.047854526 |
| | HNSC | 98 | 34596050 | 56519 | 149041908 | 0 | 0.007469878 |
| | UCEC | 5093 | 34596050 | 51524 | 149041908 | 0 | 0.425839546 |
| | GBM | 46 | 34596050 | 56571 | 149041908 | 0 | 0.003503046 |
| | PDAC | 44 | 34596050 | 56573 | 149041908 | 0 | 0.003350621 |
| | CCRCC | 88 | 34596050 | 56529 | 149041908 | 0 | 0.006706459 |
| T**[C>A]** | LUSC | 523 | 59456977 | 106490 | 253342313 | 0 | 0.020926556 |
| | LUAD | 1167 | 59456977 | 105846 | 253342313 | 0 | 0.046978734 |
| | HNSC | 210 | 59456977 | 106803 | 253342313 | 0 | 0.008378008 |
| | UCEC | 6133 | 59456977 | 100880 | 253342313 | 0 | 0.259043559 |
| | GBM | 64 | 59456977 | 106949 | 253342313 | 0 | 0.002549812 |
| | PDAC | 72 | 59456977 | 106941 | 253342313 | 0 | 0.002868753 |
| | CCRCC | 237 | 59456977 | 106776 | 253342313 | 0 | 0.009457571 |
| T**[C>T]**GG | LUSC | 43 | 343734 | 6008 | 1193647 | 0 | 0.024853751 |
| | LUAD | 83 | 343734 | 5968 | 1193647 | 0 | 0.048295059 |
| | HNSC | 59 | 343734 | 5992 | 1193647 | 0 | 0.034192718 |
| | UCEC | 764 | 343734 | 5287 | 1193647 | 3.32613E-83 | 0.501807869 |
| | GBM | 59 | 343734 | 5992 | 1193647 | 0 | 0.034192718 |
| | PDAC | 32 | 343734 | 6019 | 1193647 | 0 | 0.018462013 |
| | CCRCC | 42 | 343734 | 6009 | 1193647 | 0 | 0.024271717 |
| **[T>C]**G | LUSC | 214 | 72921280 | 34742 | 307483607 | 0 | 0.025973271 |
| | LUAD | 286 | 72921280 | 34670 | 307483607 | 0 | 0.034784028 |
| | HNSC | 101 | 72921280 | 34855 | 307483607 | 0 | 0.012218671 |
| | UCEC | 1171 | 72921280 | 33785 | 307483607 | 0 | 0.146150618 |
| | GBM | 75 | 72921280 | 34881 | 307483607 | 0 | 0.009066507 |
| | PDAC | 131 | 72921280 | 34825 | 307483607 | 0 | 0.015861631 |
| | CCRCC | 227 | 72921280 | 34729 | 307483607 | 0 | 0.027561399 |
| **[C>T]**CA | LUSC | 6 | 9573625 | 8929 | 37732952 | 0 | 0.002648456 |
| | LUAD | 8 | 9573625 | 8927 | 37732952 | 0 | 0.003532066 |
| | HNSC | 8 | 9573625 | 8927 | 37732952 | 0 | 0.003532066 |
| | UCEC | 22 | 9573625 | 8913 | 37732952 | 0 | 0.009728439 |
| CT**[C>T]**G | LUSC | 22 | 465889 | 4581 | 1601820 | 0 | 0.016511771 |
| | LUAD | 50 | 465889 | 4553 | 1601820 | 0 | 0.037757533 |
| | HNSC | 29 | 465889 | 4574 | 1601820 | 0 | 0.021798825 |
| | UCEC | 348 | 465889 | 4255 | 1601820 | 3.5724E-163 | 0.281197163 |
| | GBM | 27 | 465889 | 4576 | 1601820 | 0 | 0.020286588 |
| | PDAC | 12 | 465889 | 4591 | 1601820 | 0 | 0.008986803 |
| | CCRCC | 18 | 465889 | 4585 | 1601820 | 0 | 0.013497844 |
| **[C>T]**A | LUSC | 455 | 72921280 | 49342 | 307483607 | 0 | 0.03888323 |
| | LUAD | 785 | 72921280 | 49012 | 307483607 | 0 | 0.067535935 |
| | HNSC | 468 | 72921280 | 49329 | 307483607 | 0 | 0.040004719 |
| | UCEC | 1522 | 72921280 | 48275 | 307483607 | 0 | 0.132941341 |
| | GBM | 148 | 72921280 | 49649 | 307483607 | 0 | 0.012569526 |

| | | | | | | | |
|---|---|---|---|---|---|---|---|
| | PDAC | 158 | 72921280 | 49639 | 307483607 | 0 | 0.013421521 |
| | CCRCC | 308 | 72921280 | 49489 | 307483607 | 0 | 0.026242773 |
| | LUSC | 175 | 10248102 | 11356 | 43005473 | 0 | 0.064668525 |
| | LUAD | 459 | 10248102 | 11072 | 43005473 | 0 | 0.173967009 |
| | HNSC | 68 | 10248102 | 11463 | 43005473 | 0 | 0.024893784 |
| **[C>A]**TG | UCEC | 958 | 10248102 | 10573 | 43005473 | 9.7728E-236 | 0.380231039 |
| | GBM | 20 | 10248102 | 11511 | 43005473 | 0 | 0.00729117 |
| | PDAC | 31 | 10248102 | 11500 | 43005473 | 0 | 0.011312124 |
| | CCRCC | 65 | 10248102 | 11466 | 43005473 | 0 | 0.023789302 |
| | LUSC | 22 | 38081020 | 6165 | 164339306 | 0 | 0.015400062 |
| | LUAD | 29 | 38081020 | 6158 | 164339306 | 0 | 0.020323158 |
| | HNSC | 17 | 38081020 | 6170 | 164339306 | 0 | 0.011890405 |
| A**[T>A]** | UCEC | 133 | 38081020 | 6054 | 164339306 | 0 | 0.09480737 |
| | GBM | 11 | 38081020 | 6176 | 164339306 | 0 | 0.007686317 |
| | PDAC | 9 | 38081020 | 6178 | 164339306 | 0 | 0.006286769 |
| | CCRCC | 25 | 38081020 | 6162 | 164339306 | 0 | 0.017508591 |
| | LUSC | 195 | 54176761 | 60786 | 215208799 | 0 | 0.012743186 |
| | LUAD | 339 | 54176761 | 60642 | 215208799 | 0 | 0.022206145 |
| | HNSC | 293 | 54176761 | 60688 | 215208799 | 0 | 0.019178374 |
| **[C>T]**C | UCEC | 1145 | 54176761 | 59836 | 215208799 | 0 | 0.076013358 |
| | GBM | 147 | 54176761 | 60834 | 215208799 | 0 | 0.009598822 |
| | PDAC | 89 | 54176761 | 60892 | 215208799 | 0 | 0.005805996 |
| | CCRCC | 184 | 54176761 | 60797 | 215208799 | 0 | 0.012022164 |
| | LUSC | 177 | 34596050 | 9582 | 149041908 | 0 | 0.079579093 |
| | LUAD | 304 | 34596050 | 9455 | 149041908 | 0 | 0.138514085 |
| | HNSC | 166 | 34596050 | 9593 | 149041908 | 0 | 0.07454792 |
| **[C>G]**T | UCEC | 90 | 34596050 | 9669 | 149041908 | 0 | 0.040099858 |
| | GBM | 50 | 34596050 | 9709 | 149041908 | 0 | 0.022185917 |
| | PDAC | 36 | 34596050 | 9723 | 149041908 | 0 | 0.01595086 |
| | CCRCC | 90 | 34596050 | 9669 | 149041908 | 0 | 0.040099858 |
| | LUSC | 27 | 7020556 | 10259 | 29596560 | 0 | 0.01109503 |
| | LUAD | 43 | 7020556 | 10243 | 29596560 | 0 | 0.017697463 |
| | HNSC | 31 | 7020556 | 10255 | 29596560 | 0 | 0.012743706 |
| AG**[C>T]** | UCEC | 352 | 7020556 | 9934 | 29596560 | 0 | 0.149378549 |
| | GBM | 36 | 7020556 | 10250 | 29596560 | 0 | 0.014806362 |
| | PDAC | 14 | 7020556 | 10272 | 29596560 | 0 | 0.005745697 |
| | CCRCC | 20 | 7020556 | 10266 | 29596560 | 0 | 0.008212936 |
| | LUSC | 269 | 76162040 | 31256 | 328678612 | 0 | 0.037140843 |
| | LUAD | 391 | 76162040 | 31134 | 328678612 | 0 | 0.054196933 |
| | HNSC | 208 | 76162040 | 31317 | 328678612 | 0 | 0.028662631 |
| A**[T>C]** | UCEC | 1038 | 76162040 | 30487 | 328678612 | 0 | 0.146931712 |
| | GBM | 132 | 76162040 | 31393 | 328678612 | 0 | 0.018145711 |
| | PDAC | 110 | 76162040 | 31415 | 328678612 | 0 | 0.015110836 |
| | CCRCC | 252 | 76162040 | 31273 | 328678612 | 0 | 0.034774738 |
| | LUSC | 318 | 27052755 | 33401 | 107095832 | 0 | 0.037690224 |
| | LUAD | 491 | 27052755 | 33228 | 107095832 | 0 | 0.058497642 |
| | HNSC | 357 | 27052755 | 33362 | 107095832 | 0 | 0.042362073 |
| C**[C>T]** | UCEC | 1522 | 27052755 | 32197 | 107095832 | 0 | 0.187137281 |
| | GBM | 360 | 27052755 | 33359 | 107095832 | 0 | 0.042721899 |
| | PDAC | 187 | 27052755 | 33532 | 107095832 | 0 | 0.022077161 |
| | CCRCC | 269 | 27052755 | 33450 | 107095832 | 0 | 0.031835907 |
| | LUSC | 452 | 59456977 | 25557 | 253342313 | 0 | 0.075358712 |
| | LUAD | 713 | 59456977 | 25296 | 253342313 | 0 | 0.120099883 |
| | HNSC | 410 | 59456977 | 25599 | 253342313 | 0 | 0.068244203 |
| T**[C>G]** | UCEC | 135 | 59456977 | 25874 | 253342313 | 0 | 0.022231824 |
| | GBM | 74 | 59456977 | 25935 | 253342313 | 0 | 0.012157671 |
| | PDAC | 90 | 59456977 | 25919 | 253342313 | 0 | 0.014795484 |
| | CCRCC | 174 | 59456977 | 25835 | 253342313 | 0 | 0.028697607 |
| | LUSC | 251 | 34596050 | 29166 | 149041908 | 0 | 0.037074793 |
| | LUAD | 389 | 34596050 | 29028 | 149041908 | 0 | 0.057731703 |
| **[C>T]**T | HNSC | 304 | 34596050 | 29113 | 149041908 | 0 | 0.044985081 |
| | UCEC | 1077 | 34596050 | 28340 | 149041908 | 0 | 0.163718495 |
| | GBM | 106 | 34596050 | 29311 | 149041908 | 0 | 0.015579629 |

| | | | | | | | |
|---|---|---|---|---|---|---|---|
| | PDAC | 109 | 34596050 | 29308 | 149041908 | 0 | 0.016022202 |
| | CCRCC | 172 | 34596050 | 29245 | 149041908 | 0 | 0.025337205 |
| T**[C>T]** | LUSC | 583 | 59456977 | 99310 | 253342313 | 0 | 0.025013846 |
| | LUAD | 991 | 59456977 | 98902 | 253342313 | 0 | 0.042694653 |
| | HNSC | 906 | 59456977 | 98987 | 253342313 | 0 | 0.038999132 |
| | UCEC | 2765 | 59456977 | 97128 | 253342313 | 0 | 0.121298546 |
| | GBM | 234 | 59456977 | 99659 | 253342313 | 0 | 0.010004704 |
| | PDAC | 212 | 59456977 | 99681 | 253342313 | 0 | 0.00906209 |
| | CCRCC | 354 | 59456977 | 99539 | 253342313 | 0 | 0.015153568 |
| C**[T>C]** | LUSC | 52 | 34596050 | 13990 | 149041908 | 0 | 0.016012809 |
| | LUAD | 85 | 34596050 | 13957 | 149041908 | 0 | 0.026236671 |
| | HNSC | 30 | 34596050 | 14012 | 149041908 | 0 | 0.009223654 |
| | UCEC | 150 | 34596050 | 13892 | 149041908 | 0 | 0.046516644 |
| | GBM | 21 | 34596050 | 14021 | 149041908 | 0 | 0.006452413 |
| | PDAC | 25 | 34596050 | 14017 | 149041908 | 0 | 0.007683637 |
| | CCRCC | 45 | 34596050 | 13997 | 149041908 | 0 | 0.013850308 |
| **[T>C]**A | LUSC | 323 | 63871152 | 29524 | 279318776 | 0 | 0.047843474 |
| | LUAD | 373 | 63871152 | 29474 | 279318776 | 0 | 0.05534331 |
| | HNSC | 190 | 63871152 | 29657 | 279318776 | 0 | 0.028017009 |
| | UCEC | 1700 | 63871152 | 28147 | 279318776 | 0 | 0.264126629 |
| | GBM | 115 | 63871152 | 29732 | 279318776 | 0 | 0.016914887 |
| | PDAC | 106 | 63871152 | 29741 | 279318776 | 0 | 0.015586395 |
| | CCRCC | 222 | 63871152 | 29625 | 279318776 | 0 | 0.032771023 |
| **[T>C]**GA | LUSC | 40 | 19569923 | 9759 | 83631154 | 0 | 0.017515948 |
| | LUAD | 60 | 19569923 | 9739 | 83631154 | 0 | 0.026327878 |
| | HNSC | 41 | 19569923 | 9758 | 83631154 | 0 | 0.017955687 |
| | UCEC | 479 | 19569923 | 9320 | 83631154 | 0 | 0.219633496 |
| | GBM | 17 | 19569923 | 9782 | 83631154 | 0 | 0.007426774 |
| | PDAC | 31 | 19569923 | 9768 | 83631154 | 0 | 0.013562352 |
| | CCRCC | 52 | 19569923 | 9747 | 83631154 | 0 | 0.022798766 |
| G**[C>T]**G | LUSC | 46 | 1331707 | 19760 | 4885981 | 0 | 0.008541103 |
| | LUAD | 79 | 1331707 | 19727 | 4885981 | 0 | 0.014692955 |
| | HNSC | 56 | 1331707 | 19750 | 4885981 | 0 | 0.01040313 |
| | UCEC | 842 | 1331707 | 18964 | 4885981 | 0 | 0.162901557 |
| | GBM | 155 | 1331707 | 19651 | 4885981 | 0 | 0.028939441 |
| | PDAC | 79 | 1331707 | 19727 | 4885981 | 0 | 0.014692955 |
| | CCRCC | 62 | 1331707 | 19744 | 4885981 | 0 | 0.011521251 |
| **[C>T]**GAG | LUSC | 19 | 465889 | 5713 | 1601820 | 0 | 0.011434591 |
| | LUAD | 35 | 465889 | 5697 | 1601820 | 0 | 0.021122877 |
| | HNSC | 20 | 465889 | 5712 | 1601820 | 0 | 0.012038519 |
| | UCEC | 497 | 465889 | 5235 | 1601820 | 1.0437E-169 | 0.326415639 |
| | GBM | 39 | 465889 | 5693 | 1601820 | 0 | 0.023553458 |
| | PDAC | 33 | 465889 | 5699 | 1601820 | 0 | 0.019908867 |
| | CCRCC | 32 | 465889 | 5700 | 1601820 | 0 | 0.019302181 |
| **[C>G]**A | LUSC | 271 | 72921280 | 15642 | 307483607 | 0 | 0.073054117 |
| | LUAD | 477 | 72921280 | 15436 | 307483607 | 0 | 0.130302063 |
| | HNSC | 238 | 72921280 | 15675 | 307483607 | 0 | 0.064023158 |
| | UCEC | 108 | 72921280 | 15805 | 307483607 | 0 | 0.028813561 |
| | GBM | 59 | 72921280 | 15854 | 307483607 | 0 | 0.015692092 |
| | PDAC | 56 | 72921280 | 15857 | 307483607 | 0 | 0.014891371 |
| | CCRCC | 135 | 72921280 | 15778 | 307483607 | 0 | 0.036078585 |
| C**[T>G]** | LUSC | 11 | 34596050 | 6926 | 149041908 | 0 | 0.006842142 |
| | LUAD | 34 | 34596050 | 6903 | 149041908 | 0 | 0.021218903 |
| | HNSC | 16 | 34596050 | 6921 | 149041908 | 0 | 0.009959396 |
| | UCEC | 273 | 34596050 | 6664 | 149041908 | 3.4748E-304 | 0.176485709 |
| | GBM | 7 | 34596050 | 6930 | 149041908 | 0 | 0.004351577 |
| | PDAC | 11 | 34596050 | 6926 | 149041908 | 0 | 0.006842142 |
| | CCRCC | 23 | 34596050 | 6914 | 149041908 | 0 | 0.014331127 |
| **[C>A]**C | LUSC | 321 | 54176761 | 46485 | 215208799 | 0 | 0.027430845 |
| | LUAD | 787 | 54176761 | 46019 | 215208799 | 0 | 0.067933586 |
| | HNSC | 95 | 54176761 | 46711 | 215208799 | 0 | 0.008078885 |
| | UCEC | 321 | 54176761 | 46485 | 215208799 | 0 | 0.027430845 |
| | GBM | 53 | 54176761 | 46753 | 215208799 | 0 | 0.004503118 |

| | | | | | | | |
|---|---|---|---|---|---|---|---|
| | PDAC | 45 | 54176761 | 46761 | 215208799 | 0 | 0.003822748 |
| | CCRCC | 124 | 54176761 | 46682 | 215208799 | 0 | 0.010551622 |
| **[C>A]**A | LUSC | 624 | 72921280 | 47603 | 307483607 | 0 | 0.055273625 |
| | LUAD | 1457 | 72921280 | 46770 | 307483607 | 0 | 0.13135901 |
| | HNSC | 219 | 72921280 | 48008 | 307483607 | 0 | 0.019235265 |
| | UCEC | 933 | 72921280 | 47294 | 307483607 | 0 | 0.083184667 |
| | GBM | 99 | 72921280 | 48128 | 307483607 | 0 | 0.008673713 |
| | PDAC | 118 | 72921280 | 48109 | 307483607 | 0 | 0.010342448 |
| | CCRCC | 272 | 72921280 | 47955 | 307483607 | 0 | 0.023916779 |
| **[T>A]**G | LUSC | 83 | 36831550 | 7763 | 152922881 | 0 | 0.044391619 |
| | LUAD | 197 | 36831550 | 7649 | 152922881 | 0 | 0.106933564 |
| | HNSC | 37 | 36831550 | 7809 | 152922881 | 0 | 0.019672465 |
| | UCEC | 60 | 36831550 | 7786 | 152922881 | 0 | 0.031995531 |
| | GBM | 13 | 36831550 | 7833 | 152922881 | 0 | 0.006890769 |
| | PDAC | 12 | 36831550 | 7834 | 152922881 | 0 | 0.006359898 |
| | CCRCC | 58 | 36831550 | 7788 | 152922881 | 0 | 0.030921071 |
| G**[C>T]** | LUSC | 113 | 44749150 | 91427 | 177071834 | 0 | 0.004890673 |
| | LUAD | 177 | 44749150 | 91363 | 177071834 | 0 | 0.007665979 |
| | HNSC | 118 | 44749150 | 91422 | 177071834 | 0 | 0.005107354 |
| | UCEC | 1347 | 44749150 | 90193 | 177071834 | 0 | 0.059096187 |
| | GBM | 82 | 44749150 | 91458 | 177071834 | 0 | 0.003547781 |
| | PDAC | 67 | 44749150 | 91473 | 177071834 | 0 | 0.002898322 |
| | CCRCC | 92 | 44749150 | 91448 | 177071834 | 0 | 0.003980873 |
| GG**[C>T]** | LUSC | 143 | 6412663 | 15022 | 24143689 | 0 | 0.035840453 |
| | LUAD | 189 | 6412663 | 14976 | 24143689 | 0 | 0.047515049 |
| | HNSC | 205 | 6412663 | 14960 | 24143689 | 0 | 0.051592608 |
| | UCEC | 1877 | 6412663 | 13288 | 24143689 | 1.8044E-168 | 0.531826366 |
| | GBM | 308 | 6412663 | 14857 | 24143689 | 0 | 0.078052138 |
| | PDAC | 126 | 6412663 | 15039 | 24143689 | 0 | 0.031544002 |
| | CCRCC | 167 | 6412663 | 14998 | 24143689 | 0 | 0.041922612 |
| **[T>G]**TGT | LUAD | 7 | 2753854 | 1638 | 11604367 | 4.5412E-138 | 0.018007967 |
| | UCEC | 199 | 2753854 | 1446 | 11604367 | 1.68075E-14 | 0.579916315 |
| **[C>G]**TG | LUSC | 122 | 10248102 | 2991 | 43005473 | 3.8588E-143 | 0.171168446 |
| | LUAD | 156 | 10248102 | 2957 | 43005473 | 3.719E-119 | 0.221387738 |
| | HNSC | 47 | 10248102 | 3066 | 43005473 | 5.3805E-214 | 0.064328881 |
| | UCEC | 63 | 10248102 | 3050 | 43005473 | 3.5477E-196 | 0.086680418 |
| | GBM | 17 | 10248102 | 3096 | 43005473 | 3.1585E-255 | 0.023042429 |
| | PDAC | 12 | 10248102 | 3101 | 43005473 | 1.0923E-263 | 0.016239018 |
| | CCRCC | 58 | 10248102 | 3055 | 43005473 | 1.345E-201 | 0.079670413 |
| T**[C>A]**C | LUSC | 115 | 7865466 | 6277 | 33061576 | 0 | 0.077009589 |
| | LUAD | 265 | 7865466 | 6127 | 33061576 | 1.2845E-280 | 0.181801343 |
| | HNSC | 40 | 7865466 | 6352 | 33061576 | 0 | 0.026469674 |
| | UCEC | 154 | 7865466 | 6238 | 33061576 | 0 | 0.103770628 |
| | GBM | 14 | 7865466 | 6378 | 33061576 | 0 | 0.00922662 |
| | PDAC | 14 | 7865466 | 6378 | 33061576 | 0 | 0.00922662 |
| | CCRCC | 46 | 7865466 | 6346 | 33061576 | 0 | 0.030468906 |

(b) Contingency table for Fisher's exact test assessing the association between mutational motifs across different cancer types and DNA regions (nucleosomal vs. non-nucleosomal). An odds ratio greater than 1 indicates that a mutational motif is enriched in a specific cancer type and within nucleosomal regions, whereas an odds ratio less than 1 indicates depletion.

Odds Ratio = (a*d)/(c*b)

| | # Mutational motif in a specific cancer type | #Mutations |
|---|---|---|
| Well-positioned nucleosomal regions | a | b |
| Inter-nucleosomal regions | c | d |

**Table S21.** (a) The association between nucleosomal structural features and mutations across different cancer types, including clear cell renal cell carcinoma (CCRCC), glioblastoma (GBM), head and neck squamous cell carcinoma (HNSC), lung squamous cell carcinoma (LUSC), lung adenocarcinoma (LUAD), pancreatic ductal adenocarcinoma (PDAC), and uterine corpus endometrial carcinoma (UCEC).

| Cancer type | Elements of contingency table | | | | P-value | Odds ratio |
|---|---|---|---|---|---|---|
| | a | b | c | d | | |
| CCRCC | 192748 | 153788 | 172842 | 146040 | 3.26589E-31 | 1.0589854 |
| GBM | 104830 | 83574 | 88310 | 73743 | 1.03507E-11 | 1.047430736 |
| HNSC | 148412 | 127286 | 132765 | 114787 | 0.147223833 | 1.008085771 |
| LUSC | 202180 | 172498 | 183515 | 162224 | 6.53814E-14 | 1.036090438 |
| LUAD | 405628 | 330422 | 366482 | 313551 | 8.01387E-48 | 1.050302773 |
| PDAC | 126182 | 100288 | 102692 | 88934 | 4.07046E-43 | 1.089631501 |
| UCEC | 1031036 | 912540 | 986513 | 850575 | 6.74322E-37 | 0.974163202 |

(b) Contingency table used in Fisher's exact test to assess the association between the presence or absence of mutations minor groove sites across different cancer types. An odds ratio greater than 1 indicates that the mutation is enriched in minor groove facing out sites, whereas an odds ratio less than 1 indicates it is depleted.

Odds Ratio = (a*d)/(c*b)

| | Minor grooves facing out the histone octamer | Minor grooves facing the histone octamer |
|---|---|---|
| Nucleosomal DNA | a | b |
| Inter-nucleosomal DND | c | d |

**Table S22.** (a) Methylation motifs across different cancer types within well-positioned nucleosomal DNA and non-nucleosomal regions. We included clear cell renal cell carcinoma (CCRCC), glioblastoma (GBM), head and neck squamous cell carcinoma (HNSC), lung squamous cell carcinoma (LUSC), lung adenocarcinoma (LUAD), pancreatic ductal adenocarcinoma (PDAC), and uterine corpus endometrial carcinoma (UCEC) for this analysis.

| Methylation motif | Cancer type | Elements of contingency table | | | | Odds ratio | P-value |
|---|---|---|---|---|---|---|---|
| | | a | b | c | d | | |
| CG | CCRCC | 93081 | 5348557 | 417513 | 20166770 | 0.84060265 | 0 |
| | GBM | 101780 | 5339858 | 445636 | 20138647 | 0.861356153 | 0 |
| | HNSC | 83306 | 5358332 | 376136 | 20208147 | 0.835272636 | 0 |
| | LUSC | 89660 | 5351978 | 400400 | 20183883 | 0.844491079 | 0 |
| | LUAD | 103043 | 5338595 | 448172 | 20136111 | 0.867206242 | 0 |
| | PDAC | 107675 | 5333963 | 457160 | 20127123 | 0.888747431 | 2.7745E-266 |
| | UCEC | 100967 | 5340671 | 438238 | 20146045 | 0.869087281 | 0 |
| CGG | CCRCC | 23635 | 1521560 | 100561 | 5620219 | 0.868140824 | 3.49077E-86 |
| | GBM | 26371 | 1518824 | 108800 | 5611980 | 0.895584084 | 2.83345E-58 |
| | HNSC | 21109 | 1524086 | 90917 | 5629863 | 0.857651627 | 2.97512E-91 |
| | LUSC | 22985 | 1522210 | 97470 | 5623310 | 0.871146101 | 7.52152E-80 |
| | LUAD | 27070 | 1518125 | 110901 | 5609879 | 0.901983837 | 2.3118E-52 |
| | PDAC | 29053 | 1516142 | 115871 | 5604909 | 0.926925684 | 8.90119E-31 |
| | UCEC | 26195 | 1519000 | 107563 | 5613217 | 0.899931709 | 5.36722E-53 |
| GCG | CCRCC | 948 | 1341371 | 3023 | 4924095 | 1.151191835 | 0.000187174 |
| | GBM | 1218 | 1341101 | 3781 | 4923337 | 1.182602216 | 5.13718E-07 |
| | HNSC | 858 | 1341461 | 2677 | 4924441 | 1.176570091 | 4.4455E-05 |
| | LUSC | 981 | 1341338 | 3174 | 4923944 | 1.134584802 | 0.000615971 |
| | LUAD | 1468 | 1340851 | 4494 | 4922624 | 1.199248357 | 2.8089E-09 |
| | PDAC | 1946 | 1340373 | 5933 | 4921185 | 1.204238501 | 2.58383E-12 |
| | UCEC | 1353 | 1340966 | 4190 | 4922928 | 1.185467061 | 8.36854E-08 |
| CGAA | CCRCC | 4166 | 292998 | 18617 | 1116614 | 0.852801547 | 7.45543E-21 |
| | GBM | 4447 | 292717 | 19515 | 1115716 | 0.868568929 | 1.51297E-17 |
| | HNSC | 3779 | 293385 | 16974 | 1118257 | 0.848587077 | 3.18169E-20 |
| | LUSC | 3990 | 293174 | 17791 | 1117440 | 0.854813352 | 1.72955E-19 |
| | LUAD | 4419 | 292745 | 19322 | 1115909 | 0.871788665 | 1.38151E-16 |
| | PDAC | 4391 | 292773 | 19171 | 1116060 | 0.873122521 | 3.6061E-16 |
| | UCEC | 4358 | 292806 | 19140 | 1116091 | 0.867890484 | 2.12212E-17 |
| CGTT | CCRCC | 7520 | 301635 | 34519 | 1423255 | 1.027923085 | 0.033107064 |
| | GBM | 7882 | 301273 | 36065 | 1421709 | 1.03133794 | 0.014639323 |
| | HNSC | 6399 | 302756 | 29346 | 1428428 | 1.028794877 | 0.042123534 |
| | LUAD | 7648 | 301507 | 35108 | 1422666 | 1.027891661 | 0.031913218 |
| | PDAC | 7376 | 301779 | 33493 | 1424281 | 1.03937801 | 0.003103454 |
| | UCEC | 7597 | 301558 | 34433 | 1423341 | 1.041370736 | 0.001663241 |
| **GCG (Flanking inker)** | CCRCC | 157 | 301334 | 3814 | 5964132 | 0.81473819 | 0.01157579 |
| | GBM | 1218 | 1341101 | 3781 | 4923337 | 1.182602216 | 5.13718E-07 |
| | HNSC | 858 | 1341461 | 2677 | 4924441 | 1.176570091 | 4.4455E-05 |
| | LUSC | 981 | 1341338 | 3174 | 4923944 | 1.134584802 | 0.000615971 |
| | LUAD | 1468 | 1340851 | 4494 | 4922624 | 1.199248357 | 2.8089E-09 |
| | PDAC | 1946 | 1340373 | 5933 | 4921185 | 1.204238501 | 2.58383E-12 |
| | UCEC | 1353 | 1340966 | 4190 | 4922928 | 1.185467061 | 8.36854E-08 |

(b) Contingency table for Fisher's exact test assessing the association between methylation motifs across different cancer types and DNA regions (nucleosomal vs. non-nucleosomal). An odds ratio greater than 1 indicates that a methylation motif is enriched in a specific cancer type and within well-positioned nucleosomal regions, whereas an odds ratio less than 1 indicates depletion.

Odds Ratio = (a*d)/(c*b)

| | # Methylation motif in a specific cancer type | #Methylations |
|---|---|---|
| Well-positioned nucleosomal regions | a | b |
| Non-nucleosomal regions | c | d |

**Table S23.** (a) Associations between CpG methylation and nucleosomal structural features across cancer types. We included clear cell renal cell carcinoma (CCRCC), glioblastoma (GBM), head and neck squamous cell carcinoma (HNSC), lung squamous cell carcinoma (LUSC), lung adenocarcinoma (LUAD), pancreatic ductal adenocarcinoma (PDAC), and uterine corpus endometrial carcinoma (UCEC) for this analysis.

| Cancer type | Elements of contingency table | | | | P-value | Odds ratio |
|---|---|---|---|---|---|---|
| | a | b | c | d | | |
| CCRCC | 362525782 | 309866774 | 332230013 | 281369200 | 0 | 0.990835549 |
| GBM | 119765080 | 101684088 | 110506186 | 93356300 | 1.5075E-144 | 0.995025595 |
| HNSC | 90185078 | 77754336 | 80907477 | 71776133 | 0 | 1.028967047 |
| LUSC | 120319750 | 103633482 | 111047118 | 97252483 | 0 | 1.01678759 |
| LUAD | 268499552 | 231995148 | 245077645 | 212282275 | 3.91574E-81 | 1.002477648 |
| PDAC | 116916846 | 101003396 | 106184042 | 91394509 | 7.57532E-78 | 0.996327152 |
| UCEC | 143444256 | 122831198 | 130801532 | 112782055 | 0 | 1.006935489 |

(b) Contingency table used in Fisher's exact test to assess the association between the presence or absence of methylations in minor groove sites across different cancer types. An odds ratio greater than 1 indicates that methylation is enriched in minor groove facing out sites, whereas an odds ratio less than 1 indicates it is depleted.

Odds Ratio = (a*d)/(c*b)

| | Minor grooves facing out the histone octamer | Minor grooves facing the histone octamer |
|---|---|---|
| Nucleosomal DNA | a | b |
| Inter-nucleosomal DND | c | d |

**Table S24.** (a) Associations between CpG methylation in regions close to the dyad position (at most 20 bases distance) and nucleosomal structural features across all cancer types.

| Cancer type | Elements of contingency table | | | | P-value | Odds ratio |
|---|---|---|---|---|---|---|
| | a | b | c | d | | |
| All cancer types | 710909884 | 1559514882 | 707510571 | 1364462539 | 0 | 0.879131239 |

(b) Contingency table used in Fisher's exact test to assess the association between the presence or absence of methylations in regions close to the dyad position across different cancer types. An odds ratio greater than 1 indicates that methylation is enriched in regions close to the dyad position, whereas an odds ratio less than 1 indicates it is depleted.

Odds Ratio = (a*d)/(c*b)

| | Regions close to the dyad position | Other regions |
|---|---|---|
| Nucleosomal DNA | a | b |
| Inter-nucleosomal DND | c | d |
